# Bifurcation dynamics in a shared network beyond sensory areas characterize conscious auditory perception independently of report

**DOI:** 10.64898/2026.06.17.732590

**Authors:** J Boyer, N Beraud, B Beranger, T Hardy, B Türker, H Gouyette, A Lopez-Persem, C Sergent

## Abstract

The neural correlates of conscious perception remain debated, particularly regarding the role of extra-sensory regions such as the prefrontal cortex. One promising approach is to study the dynamical properties of neural processing in task-related and task-free contexts. A previous EEG study showed that conscious perception is associated with all-or-none late activations, giving rise to bifurcation dynamics even without a task. Here we used fMRI and near-threshold auditory stimulation to ask which brain networks give rise to these bifurcations, and how they differ depending on task. In both contexts, stimulus intensity modulated activity within broad networks spanning sensory and extra-sensory regions, including the prefrontal cortex. These networks showed both shared and distinct components. Single-trial modelling further revealed bifurcation dynamics beyond primary sensory cortices and enabled single-trial prediction of conscious perception in both contexts. These findings resolve previous conflicting results and reveal common networks and dynamics underlying conscious perception irrespective of task.

## Introduction

What differentiates the neural correlates of “conscious access”, i.e. consciously perceiving a given stimulus, from its unconscious processing? Classically this has been investigated by contrasting brain activity according to participants’ explicit report of either perceiving a presented stimulus (i.e., conscious processing) or not (i.e., unconscious processing). These studies probed the visual modality^1^, but also other modalities, notably audition^2–14^. This approach has proven particularly fruitful: it has demonstrated that the brain’s response can be drastically different depending on the reported experience, even for the exact same external stimulation^15^. These differences involve not only sensory areas but also a wider network of extra-sensory regions, often including the prefrontal cortex (PFC)^16,17^, in association with differences during late stages of processing^18^. Such observations were in good accordance with some models of conscious perception such as the Global Neuronal Workspace framework^19,20^. According to this framework, conscious access relies on a late, sustained, broad activity involving distant but strongly connected higher-order areas, notably in the parietal and frontal lobes, and occurring beyond the initial stages of sensory analysis. The activity of this broad neural network, or “global workspace”, in response to a stimulus would critically enable that stimulus to become available to many distributed computational resources, way beyond the automatic routes specialized in its processing, and the resulting functional properties would correspond to conscious experience.

However, this classical experimental approach, which uses report as a marker of conscious perception, has shown its limits. There is indeed a risk of confusion between the actual neural correlates of conscious perception and those associated with reporting the stimulus (task-related activity). It has notably been argued that the distributed areas showing late sustained activity could in fact reflect task-related activity and not conscious perception *per se*^15,21–23^. Consequently, so-called ’no-report’ paradigms have been proposed to try and solve this issue^24^: the idea is to probe conscious processing in situations where participants are not requested to report it. Neuroimaging studies using these types of paradigms have yielded discordant results regarding the brain regions involved in such task-free conscious perception^5,9,13,25–31^. The involvement of the PFC in particular has become a central topic of debate^32,33^. Some studies suggested that prefrontal activations were abolished^13,28,31^ or even deactivated^26^ during task-free conscious processing while others suggested that conscious contents were still reflected in the activity of prefrontal areas even in the absence of report^30,34^. Several recent studies introduced interesting nuances: Hatamimajoumerd and colleagues (2022), with visual stimuli, and Dellert and colleagues (2025), with auditory stimuli, found no significant activity in the prefrontal cortex in task-free conditions when using classical univariate fMRI analysis; yet they showed that the use of more complex techniques, respectively multivariate pattern analysis (MVPA) and Bayesian approaches, led to a change in conclusion^5,29^.

One persistent problem that all these studies face is the need to rely on indirect markers of awareness, which are then used to contrast putatively conscious and unconscious processing. This circularity might limit the identification of intrinsic neural mechanisms underlying conscious access. Here we explore a complementary approach which could help overcome this fundamental limitation by inverting the logic: it consists in characterizing the dynamics of the brain’s response to stimulation around threshold, with no a priori labelling of trials as being conscious or not, in search for properties that would indicate a transition between different ways of processing the same stimulus. Indeed, several previous studies on reported perception have suggested that conscious access might be linked to the all-or-none triggering of some specific neural events, beyond initial sensory processing^18,35–37^. In a previous EEG study, we developed a dynamical approach inspired from these observations to question the neural correlates of conscious access independent of report^10^. We analysed the single-trial dynamics of EEG responses elicited by auditory stimuli of various intensities around threshold in both an Active condition, where participants had to report the stimulus, and a Passive condition without report. We tested whether the intensity of EEG responses followed unimodal dynamics (Figure 1A left), with activity across trials grouping around a single mean, scaling with stimulus intensity, or bifurcation dynamics (Figure 1A right) where activity for very low or very high stimulus intensity still follows unimodal distributions (Noise and Level 5 in Figure 1A), but where activity for threshold intensities shows bimodal distributions, suggesting two possible equilibria across trials for the same external stimulation. Characterizing these dynamics involved not only looking at mean response but also assessing the variability of responses across trials as a function of stimulus intensity, which distinguishes better these two types of dynamics, as shown in Figure 1A. The results showed that, in both Active and Passive settings, early EEG waveforms followed unimodal dynamics, while later waveforms, beyond 300 milliseconds post-stimulus, showed bifurcation dynamics suggesting that, around threshold intensities, the same stimulus could elicit a strong, sustained activity on some trials, and not on others. In the Active condition, the presence or absence of these late activations matched participants’ conscious report. In the Passive condition, the presence of these late activities predicted that, if presented with a mind-wandering probe^38^, participants would mention that they had the sound on their mind^10^.

**Figure 1.**
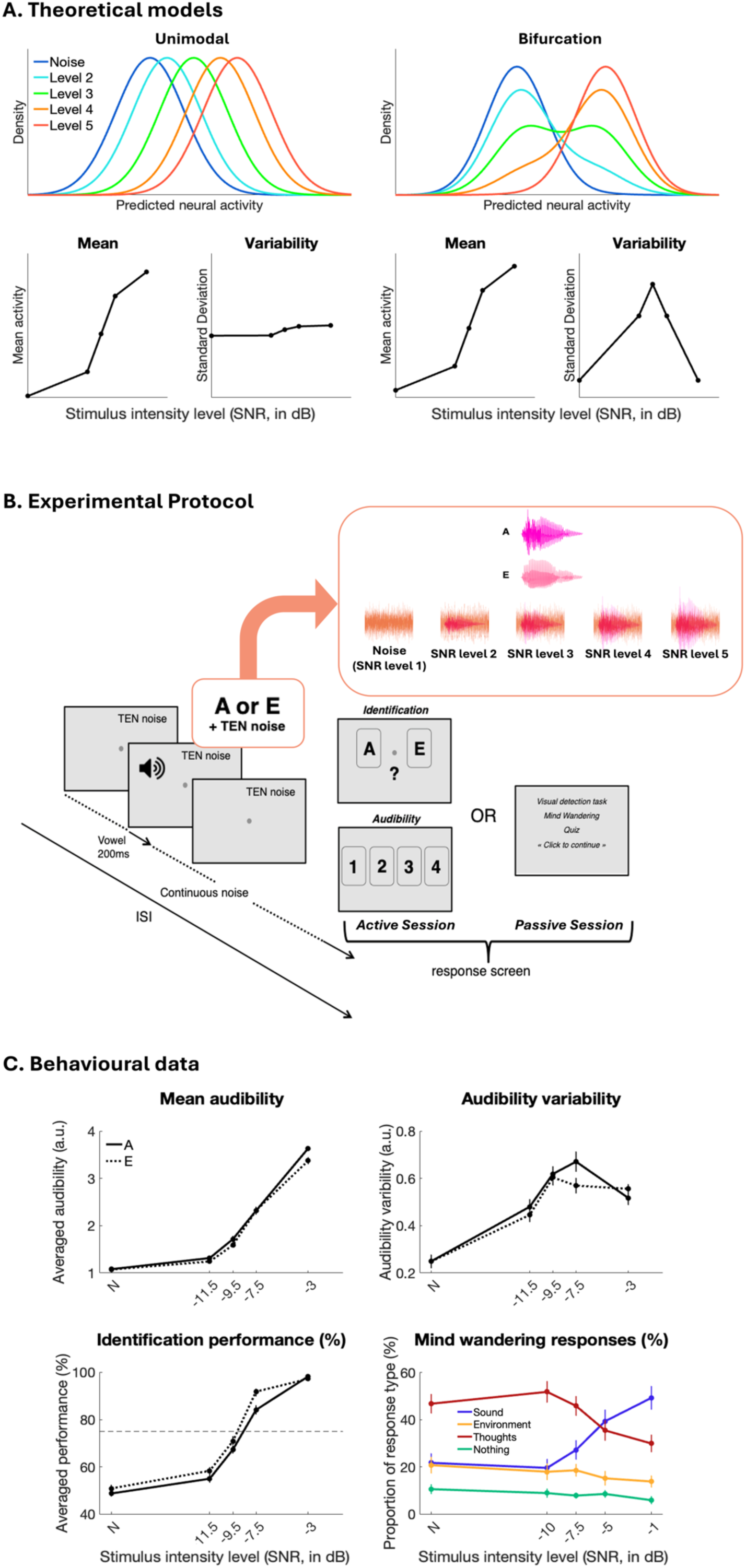
Theoretical predictions, experimental setting and behavioural results. A. Theoretical predictions for trial-by-trial dynamics of two possible models relating stimulus intensity and neural activity. For each model, simulated single-trial neural activity distribution as well as average and inter-trial variability are represented as a function of the stimulus intensity. According to the unimodal model (left panel), predicted neural activity progressively increases with stimulus strength with only unimodal distributions. On the other hand, the bifurcation model (right panel) predicts low neural activity in response to low levels of stimulus, high neural activity in response to high levels, and either level of activity across trials in response to stimuli around perception threshold. Stimulation around threshold thus yields a bimodal distribution of activity across trials, resulting in increased inter-trial variability compared to other levels of stimulation. B. Experimental paradigm: subjects heard a continuous background noise. The French vowel "A" or "E" (200 milliseconds) was superimposed to a threshold equalizing noise (see Methods) at a jittered delay of 1 to 3 seconds relative to the start of the trial that was marked by the onset of the fixation point. The vowels were played at five possible intensity levels (from no vowel, just noise (N) to the highest vowel intensity, measured in Signal to Noise Ratio - SNR - relative to the background sound; the specific intensities varied across Active and Passive conditions, see panel C). A response screen appeared with a jittered delay of 4 to 6 seconds relative to vowel onset, containing either an identification task followed by audibility rating (Active condition) or one of four distracting tasks (Passive condition: a visual detection task, mind-wandering probe, trivia quiz or just ’click to continue’ screen appeared randomly across trials). C. Behavioural results across the 27 participants included in fMRI analyses. Top left panel: averaged audibility rating as a function of stimulus intensity in the Active condition, separately for the two vowels. Top right panel: inter-trial variability of audibility as a function of stimulus intensity in the Active condition, separately for the two vowels.

Interestingly, the EEG topographies of these late activations were different in the Active and Passive conditions, suggesting different underlying networks, but the poor spatial resolution of EEG did not allow to determine which brain circuits give rise to these bifurcations, and how they differ depending on task demands. In the present study we leverage the spatial resolution of functional MRI to investigate where and how these processes occur in the brain. In adapting this dynamical approach to fMRI we pursue two main goals: (1) identify networks that encode auditory stimuli and map their overlaps and differences across the Active and Passive conditions; specifically, we are interested in testing if extra-sensory regions, notably in the prefrontal cortex, are also implicated in spontaneous conscious perception during passive listening, (2) apply the single-trial dynamics approach to fMRI data and characterize the neuroanatomy of the bifurcation dynamics we observed in EEG, with the specific prediction that bifurcation dynamics emerge from the joint activity of extra primary sensory regions while primary sensory regions might display more progressive, unimodal dynamics. If observed, this bifurcation should correspond to the split between conscious versus non-conscious processing in both cases. Finally, we also used concomitant eye-tracking to investigate whether the pupil size reflected these brain dynamics to some extend and could thus be used as a simple marker for future clinical use.

## Results

We adapted the protocol from Sergent and colleagues^10^ to fMRI constraints (Figure 1B): participants were presented with a continuous background noise on which spoken vowels "A" or "E" with variable intensity were superimposed. Depending on condition (either Active or Passive), they were instructed to either identify the vowels and rate their audibility (Active condition) or respond to distractive questions in the visual modality without paying attention to the sound (Passive condition). Importantly, 1/4th of Passive condition trials’ distractive questions were mind-wandering probes, randomly sampling participants’ current mental content. A total of 27 participants underwent both the Active and Passive conditions during two different fMRI sessions, in a randomized order. See Methods and Figure 1B for detailed description of the experimental protocol.

## 1. Behavioural response functions and behavioural evidence of bifurcation in Active and Passive conditions

Presenting auditory stimuli in an fMRI experiment can lead to several technical issues and pitfalls^39^, due to the need for MRI-compatible material and, of course, the noise of the scanner. As our experiment was based on playing stimuli precisely around the auditory threshold, it was crucial to deliver them in a carefully controlled way. This was achieved thanks to a behavioural pilot study (supplementary Figure S1) and individualized staircase procedures in the scanner (see Methods). Behavioural data obtained in the scanner confirmed that our chosen intensity values spanned the desired part of the psychometric function (Figure 1C). The bottom left panel of Figure 1C shows vowel identification performance in the Active condition: performance was at chance in the absence of a vowel (50%), vowels played at intermediate intensities (-11.5, -9.5 and - 7.5dB) spanned the steep part of the psychometric function, around the 75% threshold, and performance was close to ceiling for the highest intensity (-3dB). These psychometric curves were similar for the two vowels "A" and "E". Mean audibility across trials during the Active session showed a similar profile and were also similar across both vowels (Figure 1C top left panel). Finally, the inter-trial variability of audibility ratings showed the non-monotonous profile expected from underlying bifurcation distributions (Figure 1C top right panel), with a peak in variability for stimulation around threshold that could be linked with a mixture of “Heard” and “Not heard” trials.

During the Passive session, on the other hand, we had no direct report to verify threshold intensity, but we could use the responses to the mind-wandering probes that occurred randomly in 1/4^th^ of the trials, asking participants what was on their mind (Figure 1B). Figure 1C, bottom right panel shows that the proportion of trials where participants answered that they had "the *headphones’* sound" in their mind increased as a function of vowel intensity, with a trade-oO against the answer "My thoughts", and, to a lesser extent, against the two other types of answers “The environment” and “Nothing”, which also declined slightly with increasing vowel intensity. Note that the proportion of “The sound” answers did not go down to zero when no vowel was presented: this should, at least in part, be due to the fact that this answer includes having the sounds of the headphones on one’s mind (including the experimental background noise), not necessarily the vowel, to avoid a hidden task on the vowels. This profile of results stands in favour of the mind-wandering probes constituting a reliable though indirect measure of conscious perception of the stimulus in the Passive condition. Moreover, the switch between a majority of answers "The sound" versus "My thoughts" happened between the 3rd and 4th intensity levels (-7.5 and -5dB), confirming a shift of 2 to 3 dB in the auditory threshold in the Passive relative to the Active condition, that was correctly anticipated in the shifted intensity range. These behavioural results align well with those observed in the corresponding EEG study that inspired the current protocol^10^.

Bottom left panel: averaged identification performance as a function of stimulus intensity (% of correctly selecting "A" or "E", compared to threshold level 75%). Bottom right panel: proportion of response type selected by the participants following mind-wandering probes as a function of stimulus intensity in the Passive sessions. Error bars represent the standard error to the mean (SEM) across participants.

## 2. Networks responding to stimulus intensity in the Active and Passive conditions

To delineate the brain networks showing increased (or decreased) activation as a function of stimulus intensity in the Active and Passive conditions, we used a univariate whole-brain parametric modulation approach. The generalized linear model we tested included stimulus intensity as a parametric modulator of a categorical boxcar regressor modelling stimulus presentation (200 milliseconds between onset and offset; GLM1, see Methods). Consequently, the following results correspond to networks exhibiting activity that varied proportionally with stimulus intensity across the Active and Passive conditions.

In the Active condition, this parametric modulation uncovered a wide network of areas (see Figure 2A, Table 1 and supplementary Figure S2). As expected, increased intensity of the auditory stimulus induced increased activations in the bilateral superior temporal gyri, including the primary auditory areas as well as non-primary auditory areas located in Brodmann area 22 and in left middle temporal gyrus (MTG). We also found broad effects in frontal regions, including motor-planning regions (notably label 1), wide dorsolateral prefrontal regions bilaterally, as well as bilateral anterior insula and middle cingulate gyrus (labels 4 and 2). In the parietal lobe we found significant effects in bilateral lateral regions and precuneus (label 8). Finally, we observed effects in visual and subcortical regions, including cerebellum and thalami. The network that showed significant negative effects of stimulus intensity (supplementary Figure S6A and Table S1) was mostly the Default Mode Network (DMN). We also found negative effects in the lateral occipital cortex (LOC), the right post-central gyrus, and the left superior temporal gyrus.

**Figure 2.**
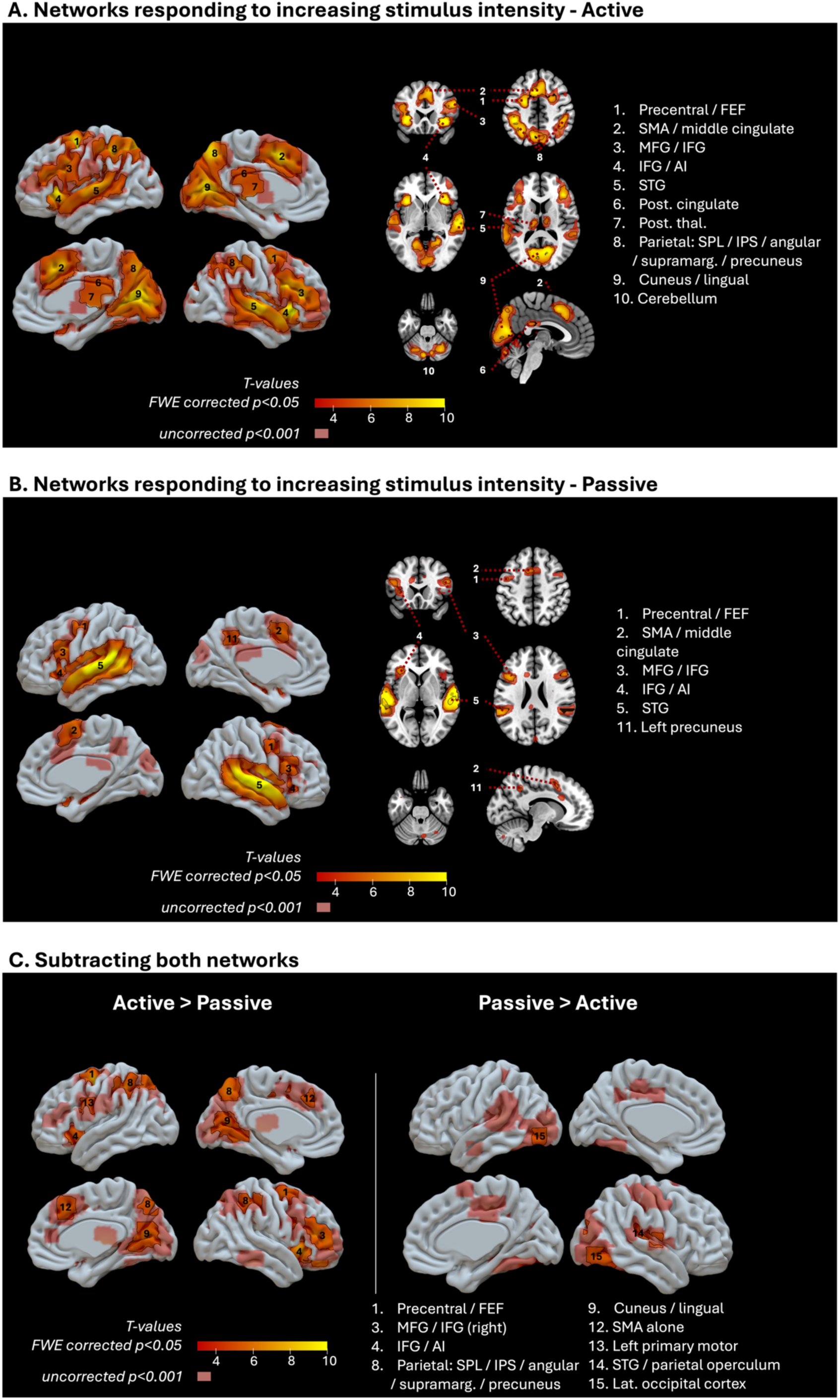
Networks encoding the auditory stimulus intensity in Active and Passive conditions. Statistical maps relating the canonical hemodynamic response to the auditory stimulus intensity in the Active condition (A) and Passive condition (B), and their contrasts (C). All slices are displayed in neurological convention (left hemisphere is on the left). Both slices and 3D surfaces display corrected activations delimited by a black contour (family-wise error correction for multiple comparisons, FWEc, p<0.05), and uncorrected ones (p<0.001) in a less opaque shade. FEF = frontal eyefield; SMA = supplementary motor area; MFG = middle frontal gyrus; IFG = inferior frontal gyrus; AI = anterior insula; post. cingulate = posterior cingulate cortex; post. thal. = posterior thalamus; SPL = superior parietal lobule; IPS = intraparietal sulcus; supramarg. = supramarginal gyrus; STG = superior temporal gyrus. Exhaustive results are displayed in Tables 1 & 2 and supplementary Figures S2 to S5.

**Table 1.** Networks encoding the auditory stimulus intensity in Active and Passive conditions. The clusters and sub-clusters peaks listed here result from whole-brain parametric modulation analyses and survived cluster-level FWE correction (p < 0.05). X, y and z coordinates refer to the Montreal Neurologic Institute (MNI) space. The last column indicates to which network(s) the clusters might belong according to Yeo et al. 2011^42^, NA = Not applicable (corresponding to subcortical areas that were not attributed to one of the networks described by Yeo et al). Uncorrected p-values are not displayed here. FEF = frontal eyefield; SMA = supplementary motor area; DMN = default mode network.

| ACTIVE |  |  |  |  |  |  |  |  |  |  |
| --- | --- | --- | --- | --- | --- | --- | --- | --- | --- | --- |
| Cluster label | Cluster size (voxels) | Side | Peak coordinates (mm) |  |  | Peak T-value | Z-score of peak T-value | Peak p-value (FWEc) | Cluster p-value (FWEc) | Network(s) of interest |
|  |  |  | x | y | z |  |  |  |  |  |
| Anterior insula | 276 | R | 37 | 23 | -7 | 13,6632 | > 10 | < 0.001 | < 0.001 | Saliency |
| Superior parietal lobule / supramarginal gyrus | 4028 | L | -41 | -38 | 41 | 13,5458 | > 10 | < 0.001 | < 0.001 | Dorsal attention / control / visual |
|  |  | R | 17 | -70 | 8 | 10,9537 | 7,7138 | < 0.001 |  |  |
|  |  | L | -19 | -70 | 6 | 10,7574 | 7,6359 | < 0.001 |  |  |
| Superior frontal gyrus (precentral / FEF / SMA) / middle cingulate / anterior insula | 2565 | L | -24 | -5 | 56 | 13,2968 | > 10 | < 0.001 | < 0.001 | Saliency / control / DMN |
|  |  | L | -9 | 13 | 46 | 11,6336 | > 10 | < 0.001 |  |  |
|  |  | L | -34 | 18 | -5 | 10,3447 | 7,4654 | < 0.001 |  |  |
| Superior temporal gyrus | 1010 | R | 57 | -18 | -5 | 11,1599 | 7,7907 | < 0.001 | < 0.001 | Auditory / DMN |
|  |  | R | 52 | 10 | -15 | 9,757 | 7,209 | < 0.001 |  |  |
|  |  | R | 67 | -23 | 6 | 9,7263 | 7,1952 | < 0.001 |  |  |
| Middle / inferior frontal gyrus | 819 | R | 49 | 18 | 26 | 10,2314 | 7,4171 | < 0.001 | < 0.001 | Saliency / control / DMN |
|  |  | R | 42 | 33 | 16 | 10,1318 | 7,3743 | < 0.001 |  |  |
|  |  | R | 37 | 20 | 8 | 5,5542 | 4,8567 | 0,0218 |  |  |
| Superior parietal lobule / supramarginal gyrus | 396 | R | 39 | -40 | 41 | 9,7711 | 7,2153 | < 0.001 | < 0.001 | Dorsal attention / control |
|  |  | R | 37 | -50 | 41 | 7,7167 | 6,1927 | < 0.001 |  |  |
|  |  | R | 34 | -63 | 41 | 6,3477 | 5,3821 | 0,002 |  |  |
| Cerebellum | 380 | R | 9 | -75 | -25 | 9,0741 | 6,8915 | < 0.001 | < 0.001 | NA |
|  |  | R | 32 | -60 | -30 | 8,9333 | 6,8234 | < 0.001 |  |  |
|  |  | R | 34 | -53 | -32 | 8,168 | 6,4357 | < 0.001 |  |  |
| Superior temporal gyrus | 673 | L | -59 | -18 | 3 | 8,8634 | 6,7892 | < 0.001 | < 0.001 | Auditory / DMN |
|  |  | L | -61 | -25 | 8 | 8,6718 | 6,6943 | < 0.001 |  |  |
|  |  | L | -54 | -33 | 6 | 8,4527 | 6,5835 | < 0.001 |  |  |
| Cerebellum | 108 | L | -9 | -80 | -30 | 8,8513 | 6,7833 | < 0.001 | < 0.001 | NA |
| Cerebellum | 107 | L | -26 | -68 | -30 | 7,6007 | 6,1283 | < 0.001 | < 0.001 | NA |
| Thalamus | 43 | L | -14 | -20 | 11 | 7,5663 | 6,1091 | < 0.001 | < 0.001 | NA |
| Cerebellum | 104 | L | 24 | -65 | -50 | 7,558 | 6,1044 | < 0.001 | < 0.001 | NA |
|  |  | R | 27 | -75 | -50 | 6,9191 | 5,7344 | < 0.001 |  |  |
| Brainstem (tectum of midbrain) | 5 | L | -4 | -35 | -7 | 6,7744 | 5,6472 | < 0.001 | 0,0078 | NA |
| Thalamus | 18 | R | 12 | -15 | 13 | 6,3027 | 5,3534 | 0,0023 | < 0.001 | NA |
| Posterior cingulate (inferior part) | 50 | R | 4 | -33 | 26 | 6,2941 | 5,3479 | 0,0023 | < 0.001 | Control / (DMN) |
| Cerebellum | 29 | L | -31 | -73 | -47 | 6,0587 | 5,1957 | 0,0048 | < 0.001 | NA |
| Middle / inferior temporal gyrus | 9 | R | 59 | -35 | -15 | 5,797 | 5,022 | 0,0106 | 0,0032 | DMN |
| Frontal pole | 6 | R | 24 | 48 | -12 | 5,6758 | 4,94 | 0,0152 | 0,0061 | Control / DMN |
| Paracingulate | 5 | L | -14 | 23 | 28 | 5,5283 | 4,8388 | 0,0236 | 0,0078 | DMN |
| Cerebellum | 2 | R | 19 | -53 | -22 | 5,4159 | 4,7607 | 0,0327 | 0,0183 | NA |
| Middle frontal gyrus (precentral) | 1 | R | 39 | 0 | 38 | 5,3387 | 4,7064 | 0,0409 | 0,0266 | Control |
| Middle frontal gyrus (precentral) | 1 | R | 42 | 3 | 41 | 5,2893 | 4,6715 | 0,0471 | 0,0266 | Control |
| PASSIVE |  |  |  |  |  |  |  |  |  |  |
| Cluster label | Cluster size (voxels) | Side | Peak coordinates (mm) |  |  | Peak T-value | Z-score of peak T-value | Peak p-value (FWEc) | Cluster p-value (FWEc) | Network(s) of interest |
|  |  |  | x | y | z |  |  |  |  |  |
| Superior / middle temporal gyrus | 1836 | R | 64 | -13 | 8 | 17,4837 | > 10 | < 0.001 | < 0.001 | Auditory / DMN |
|  |  | R | 67 | -23 | 6 | 14,1988 | > 10 | < 0.001 |  |  |
|  |  | R | 59 | -15 | -5 | 14,0521 | > 10 | < 0.001 |  |  |
| Superior temporal gyrus | 1833 | L | -64 | -28 | 11 | 15,8157 | > 10 | < 0.001 | < 0.001 | Auditory / DMN |
|  |  | L | -61 | -43 | 13 | 15,4229 | > 10 | < 0.001 |  |  |
|  |  | L | -59 | -18 | 3 | 13,463 | > 10 | < 0.001 |  |  |
| Middle / inferior frontal gyrus / anterior insula | 192 | L | -41 | 15 | 23 | 7,9725 | 6,3318 | < 0.001 | < 0.001 | Saliency / control / DMN |
|  |  | L | -36 | 25 | 3 | 6,5001 | 5,4781 | 0,0012 |  |  |
| Superior frontal gyrus (SMA) / middle cingulate | 39 | R | 9 | 3 | 63 | 7,8165 | 6,2474 | < 0.001 | < 0.001 | Saliency / DMN |
| Middle / inferior frontal gyrus | 45 | R | 49 | 20 | 23 | 6,8723 | 5,7064 | < 0.001 | < 0.001 | Saliency / control / DMN |
| Superior frontal gyrus (SMA) / middle cingulate | 50 | L | -9 | 10 | 48 | 6,5821 | 5,5292 | 0,001 | < 0.001 | Saliency / DMN |
| Superior frontal gyrus (precentral / FEF) | 17 | L | -44 | -3 | 48 | 6,475 | 5,4624 | 0,0013 | < 0.001 | Control |
| Superior frontal gyrus (precentral / FEF) | 12 | R | 49 | 3 | 46 | 6,1235 | 5,238 | 0,0039 | 0,0018 | Control |
| Precuneus | 7 | L | -14 | -50 | 43 | 5,7875 | 5,0157 | 0,0109 | 0,0049 | DMN |
| Anterior insula | 5 | R | 37 | 28 | 1 | 5,5916 | 4,8824 | 0,0196 | 0,0078 | Saliency |

In conclusion, the network responding to increasing stimulation in the Active condition encompassed, as expected, both auditory sensory, salience, and control fronto-parietal networks, with concurrent negative response of the DMN activity.

In the Passive condition (see Figure 2B, Table 1 and supplementary Figure S3) auditory regions also showed strong modulation by increasing stimulation strength. As in the Active condition, these temporal activations extended inferiorly and posteriorly to include the middle temporal gyrus and the temporoparietal junction (TPJ) possibly with a broader extent than in the Active condition (label 5 in Figure 2B; see also Table 2, Passive minus Active contrast). We also found a significant frontal effect in the Passive condition but with notable differences with the Active condition: they were much less strong and less extended (Table 1 and Table 2, Active minus Passive contrast). As shown in Figure 2B, we found significant activations in the bilateral precentral regions / frontal eye fields, label 1 (FEF; a region with high inter-individual variability, making it difficult to label precisely at group level^40,41^), middle / inferior frontal gyri (MFG / IFG; more precisely dorsal Brodmann area 44, label 3), a cluster including the supplementary motor area (SMA) and middle cingulate cortex (label 2), and a cluster encompassing the anterior insula (AI) and a part of the inferior frontal gyrus (label 4). In the parietal lobe, the only significant activation was found in the left precuneus (label 11), while the broad bilateral parietal activations found in the Active condition were absent. If we consider areas that were activated at a lower threshold, uncorrected for multiple comparisons, this network extended to posterior cingulate, middle cingulate and some parts of the cerebellum that seemed to match those observed in the Active condition. Negative effects were found mostly in regions belonging to the DMN, but still with some differences with the deactivated network found in the Active condition (supplementary Figure S6B and Table S1).

**Table 2.** DiCerences between Active and Passive networks. The clusters and sub-clusters peaks listed here result from whole-brain parametric modulation analyses and survived cluster-level FWE correction (p < 0.05). X, y and z coordinates refer to the Montreal Neurologic Institute (MNI) space. The last column indicates network(s) the clusters belong to, according to Yeo et al. 2011^42^ unless specified otherwise (NA = Not applicable). Uncorrected p-values are not displayed here. SMA = supplementary motor area; DMN = default mode network.

| ACTIVE MINUS PASSIVE |  |  |  |  |  |  |  |  |  |  |
| --- | --- | --- | --- | --- | --- | --- | --- | --- | --- | --- |
| Cluster label | Cluster size (voxels) | Side | Peak coordinates (mm) |  |  | Peak T-value | Z-score of peak T-value | Peak p-value (FWEc) | Cluster p-value (FWEc) | Network(s) of interest |
| Superior frontal gyrus (precentral) | 289 | L | -21 | -8 | 56 | 10,53 | 7,54 | < 0.001 | < 0.001 | Somato-motor / dorsal attention |
| Anterior insula | 113 | R | 36 | 20 | -7 | 9,8 | 7,23 | < 0.001 | < 0.001 | Salience |
| Superior parietal lobule / supramarginal gyrus | 175 | L | -41 | -38 | 43 | 9,23 | 6,97 | < 0.001 | < 0.001 | Dorsal attention / control |
|  |  | L | -41 | -50 | 58 | 5,57 | 4,87 | 0,021 |  |  |
| Superior frontal gyrus (precentral) | 134 | R | 26 | 2 | 53 | 7,86 | 6,27 | < 0.001 | < 0.001 | Somato-motor / dorsal attention |
| Superior parietal lobule / supramarginal gyrus | 124 | R | 39 | -40 | 40 | 7,66 | 6,16 | < 0.001 | < 0.001 | Dorsal attention / control |
| Precuneus | 107 | L | -8 | -65 | 50 | 7,1 | 5,84 | < 0.001 | < 0.001 | DMN |
| Middle / inferior frontal gyrus | 149 | R | 42 | 32 | 13 | 6,89 | 5,72 | < 0.001 | < 0.001 | Salience / control / DMN |
|  |  | R | 49 | 35 | 28 | 5,64 | 4,92 | 0,017 |  |  |
| Cuneus / lingual / intracalcarine cortex | 392 | R | 16 | -70 | 8 | 6,76 | 5,64 | 0,001 | < 0.001 | Visual |
|  |  | R | 12 | -68 | 18 | 6,56 | 5,52 | 0,001 |  |  |
|  |  | L | -18 | -68 | 6 | 6,15 | 5,26 | 0,004 |  |  |
| Middle / inferior frontal gyrus | 28 | L | -48 | 0 | 28 | 6,6 | 5,54 | 0,001 | < 0.001 | Salience / control / DMN |
| Frontal pole | 4 | R | 22 | 42 | -17 | 6,21 | 5,29 | 0,003 | 0,01 | Control / DMN |
| Superior frontal gyrus (SMA) / middle cingulate | 41 | R | 9 | 25 | 48 | 6,14 | 5,25 | 0,004 | < 0.001 | Salience / control / DMN |
| Anterior insula | 24 | L | -36 | 18 | -4 | 5,99 | 5,15 | 0,006 | < 0.001 | Salience |
| Superior parietal lobule / supramarginal gyrus | 11 | L | -31 | -55 | 46 | 5,85 | 5,06 | 0,009 | 0,002 | Dorsal attention / control |
| Cuneus / lingula / intracalcarine cortex | 7 | R | 6 | -75 | 43 | 5,58 | 4,88 | 0,02 | 0,005 | Visual |
| Middle / inferior frontal gyrus | 2 | R | 42 | 5 | 26 | 5,47 | 4,8 | 0,028 | 0,018 | Salience / control / DMN |
| Cerebellum | 2 | L | -8 | -80 | -30 | 5,39 | 4,74 | 0,035 | 0,018 | NA |
| Cerebellum | 1 | L | -24 | -68 | -30 | 5,35 | 4,72 | 0,039 | 0,027 | NA |
| PASSIVE MINUS ACTIVE |  |  |  |  |  |  |  |  |  |  |
| Cluster label | Cluster size (voxels) | Side | Peak coordinates (mm) |  |  | Peak T-value | Z-score of peak T-value | Peak p-value (FWEc) | Cluster p-value (FWEc) | Network(s) of interest |
| Lateral occipital cortex | 12 | L | -41 | -80 | -4 | 5,77 | 5 | 0,012 | 0,002 | Visual |
| Lateral occipital cortex | 29 | R | 42 | -78 | -4 | 5,58 | 4,88 | 0,02 | < 0.001 | Visual |
|  |  | R | 42 | -65 | -12 | 5,58 | 4,87 | 0,02 |  |  |
| Superior temporal gyrus / parietal operculum (temporo-parietal junction) | 7 | R | 42 | -28 | 20 | 5,53 | 4,84 | 0,023 | 0,005 | Auditory / DMN / salience |
| Superior temporal gyrus | 1 | R | 64 | -12 | 10 | 5,38 | 4,73 | 0,037 | 0,027 | Auditory |
| Lateral occipital cortex | 2 | R | 32 | -82 | 23 | 5,27 | 4,66 | 0,049 | 0,018 | Visual |

In conclusion, as in the Active condition, the network whose activity scaled with the intensity of the auditory stimulation in the Passive condition comprised both auditory and extra-auditory regions, although the latter were much less extended.

We next investigated which regions showed a stronger effect of stimulus intensity in the Active than in the Passive condition (Active minus Passive parametric modulation, Figure 2C left, Table 2 and supplementary Figure S4). This contrast revealed a fronto-parietal network in line with task-related effects (lateral parietal, dorsolateral prefrontal, anterior cingulate and premotor regions) as well as medial occipital areas previously described. Notably, we found a clear effect in left precentral (label 13) probably corresponding to the motor preparation of right-hand key press. The main focus of greater effect in the Active condition in the prefrontal cortex seemed to relate to the right anterior part.

When contrasting parametric modulation in the Passive minus the Active condition (Figure 2C right, Table 2 and supplementary Figure S5), very few effects survived multiple comparison, suggesting that most of the Passive network (Figure 2B) is included in the Active network (Figure 2A). The significant differences that survived correction for multiple comparison were a small region in the right superior temporal gyrus (STG, label 15) and bilateral regions in the lateral occipital cortex (LOC, label 15). The latter might be explained rather by a negative effect in the Active condition (supplementary Figure S6A) than by an actual stronger positive effect in the Passive condition. Extending to uncorrected statistics, we found broader, bilateral STG activations for Passive versus Active conditions. Among the other uncorrected results, we can highlight the temporo-parietal junctions (TPJ), mostly on the left.

## 3. A common network for Active and Passive listening with differing time courses of the BOLD response according to task-relevance

### 3.1. Conjunction of Active and Passive networks

Figure 3A displays on the same map, in blue and yellow, the networks responding to increasing stimulus intensity in the Active and Passive conditions respectively and, superimposed in green, the statistically significant conjunction of both, corrected for multiple comparisons (parametric modulation; GLM1, see Methods). This conjunction encompasses bilateral superior temporal gyri (auditory regions) and part of the middle temporal gyri, bilateral middle and inferior frontal gyri and anterior insulae as well as a medial cluster of SMA and middle cingulate cortex. Beyond the auditory cortex, these regions belong to the salience network, control fronto-parietal network and the DMN^42^, with some of them actually standing at the intersection between all three (see Table 3). For detailed regions and networks where an effect was found and corresponding statistics, see Table 3 and supplementary Figure S7.

**Figure 3.**
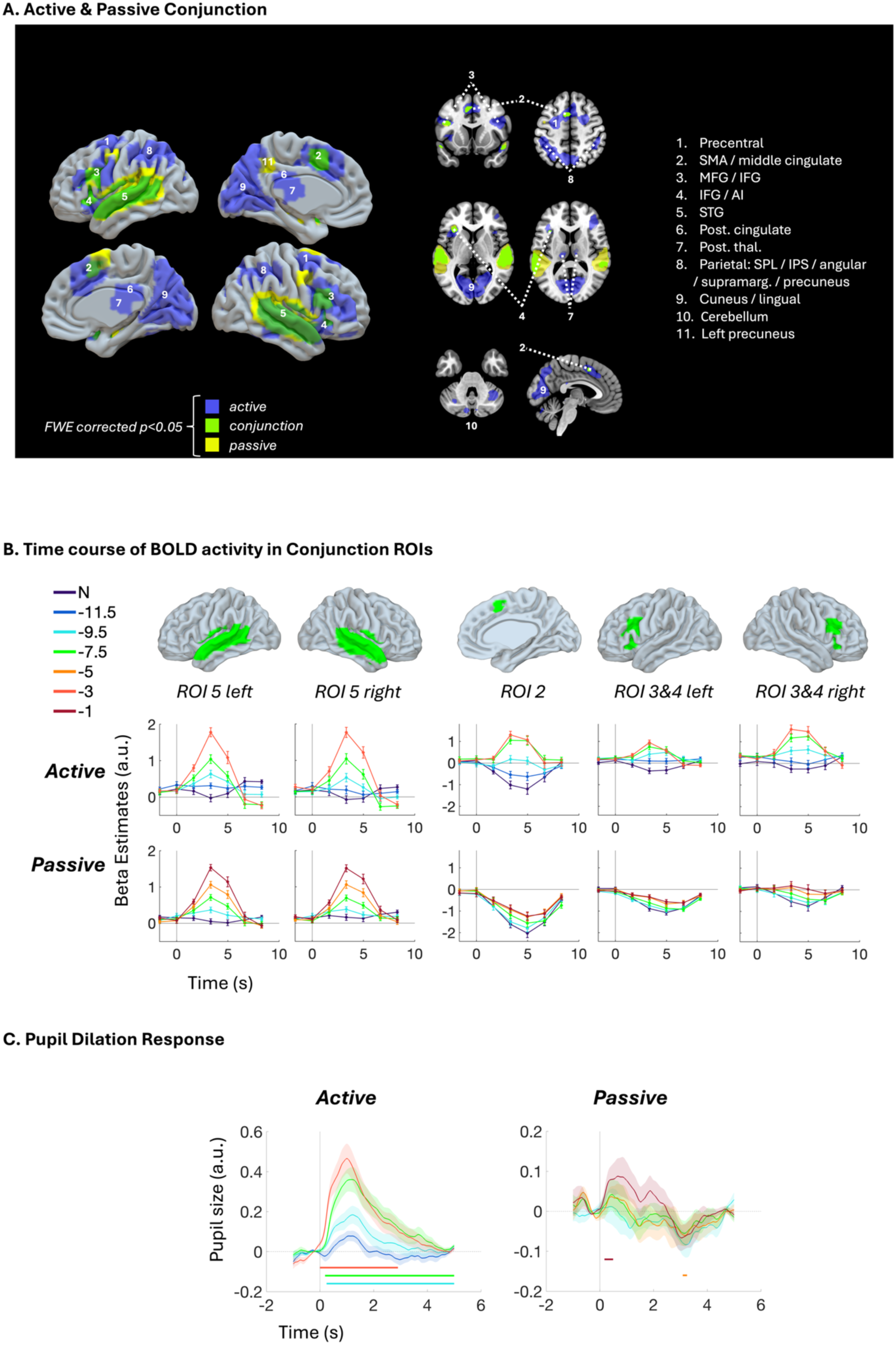
Shared network between the Active and Passive conditions, BOLD time courses within this network and associated pupil response. A: parametric modulation conjunction analysis results (in green) displaying the regions significantly responding to stimulus intensity in both the Active and Passive conditions (corrected: FWE, p<0.05). Slices are presented in neurological convention (left hemisphere is on the left). Exhaustive results are displayed in Table 3 and supplementary Figure S7. SMA = supplementary motor area; MFG = middle frontal gyrus; IFG = inferior frontal gyrus; AI = anterior insula; post. cingulate = posterior cingulate cortex; post. thal. = posterior thalamus; SPL = superior parietal lobule; IPS = intraparietal sulcus; supramarg. = supramarginal gyrus; STG = superior temporal gyrus. B: Time course of peri-stimulus fMRI activity in five conjunction-based regions of interest (depicted in the first row), shown separately for each stimulus intensity (SNR level in dB) in the Active (second row) and Passive (third row) conditions. N stands for Noise (SNR level 1). C: Time course of peri-stimulus mean pupil diameter (z-scored) shown separately for each stimulus intensity in the Active (right panel) and Passive (left panel) conditions. Shaded regions indicate ± SEM across participants (N = 23 for Active; N = 25 for Passive). Color-coded horizontal markers at the base of each plot indicate time points where the pupil response for a given intensity level was significantly different from 0 (t-tests, p < 0.05, uncorrected). Vertical and dashed lines indicate stimulus onset and the zero-baseline, respectively. Colour legend is the same as displayed for B.

**Table 3.**
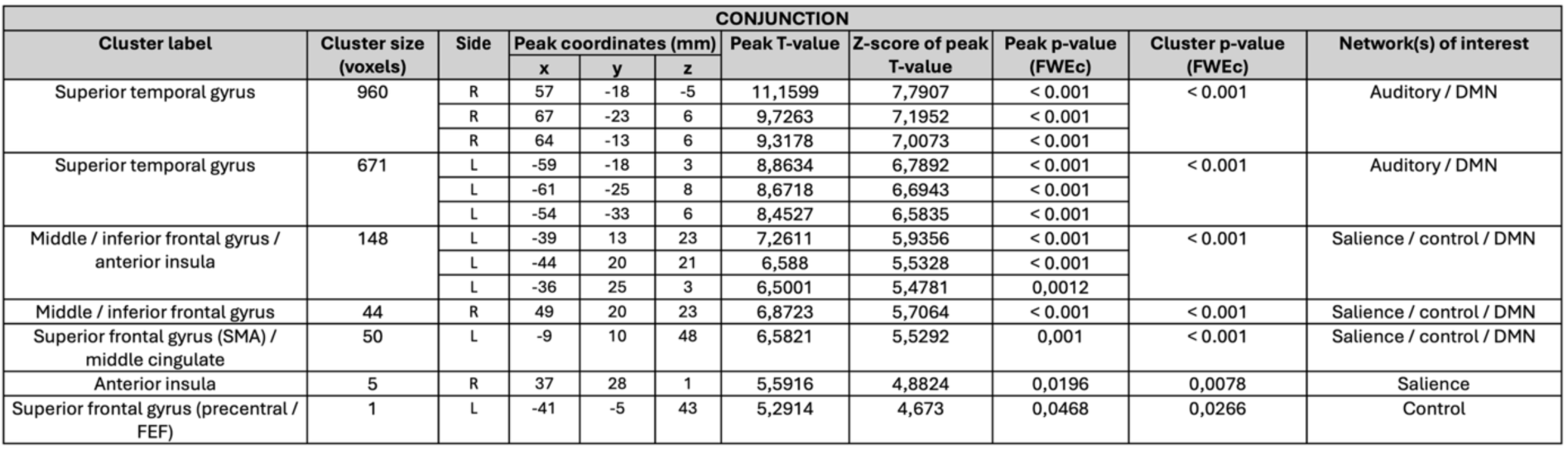
Conjunction between Active and Passive networks. The clusters and sub-clusters peaks listed here result from whole-brain parametric modulation analyses and survived cluster-level FWE correction (p < 0.05). X, y and z coordinates refer to the Montreal Neurologic Institute (MNI) space. The last column indicates network(s) the clusters belong to, according to Yeo et al. 2011^42^ unless specified otherwise (NA = Not applicable). Uncorrected p-values are not displayed here.

### 3.2. Time course of BOLD response within conjunction ROIs

To investigate the time-course of BOLD activations as a function of stimulus intensity without placing any assumption on the form, timing or amplitude of the BOLD response, we modeled the fMRI data using a GLM with a finite impulse response basis set (FIR, GLM2; see Methods). We conducted this analysis in the five main regions of the conjunction network shown in Figure 3A. This analysis does not aim at showing that these regions respond to stimulus intensity, since this was already demonstrated by the parametric modulation analysis. Instead, it aims at bringing precision on the way their activity is modulated by stimulus intensity.

As shown in Figure 3B, we found very similar time courses in the left and right superior temporal gyri (auditory regions) for both conditions, with a BOLD response that is almost flat on trials where no stimulus was presented, and whose amplitude increases gradually as a function of vowel intensity, reaching a peak around 4 seconds post stimulus onset. As expected, this response function was shifted in the Passive relative to the Active condition, with equivalent stimulation intensities yielding smaller responses in the Passive than in the Active condition. This matches the shifted auditory threshold observed in behaviour (Figure 1C). Beyond the auditory cortex, the patterns of responses observed in the three other regions of interest (ROIs) were clearly distinct in the Active and in the Passive condition. Indeed, both the PFC / insula and the SMA / middle cingulate clusters revealed negative values when no stimulus was played, much more markedly in the Passive condition, with a progressive alleviation of this relative deactivation. As discussed by Piazza, Dehaene and colleagues in another fMRI study, the BOLD signal being a relative measure, the 0 reference point corresponds to the mean activity of that region; hence, beta estimates are a relative, not an absolute, measure of activation and negative values indicate a lower signal intensity with respect to the mean activation of the region across the session^43^; see Discussion. These relative deactivations were strongest in the Passive condition, with only the right PFC / insula cluster becoming slightly positive for maximal stimulus intensity, while in the Active condition, beta estimates tended to become positive at threshold stimulus intensity or before. Furthermore, thanks to the FIR approach, we found a slightly longer latency in the rise of BOLD signal for the extra-temporal ROIs in the Passive than the Active condition (around 5 seconds post-stimulus, versus 4 seconds post-stimulus in the Active condition and in the STG cluster for both conditions).

In conclusion, although all these areas show parametric modulation with increasing stimulation strength, in details their activation patterns differ: auditory areas show similar positive patterns for both Active and Passive conditions; on the other hand, activations in extra-sensory areas emerge on top of a relative negative activity in the Passive condition while this is much less drastic in the Active condition. While we have been able to show that these regions are significantly activated in both the Active and Passive conditions, they do not have the same starting point, which might in part explain why some previous studies had difficulties finding significant activations in these regions in the absence of a task (see Discussion).

## 4. Pupil size is also modulated by the intensity of the auditory stimulus

Finally, by recording eye-tracking data simultaneously to fMRI scanning, we were able to characterize the pupil dilation response (changes in pupil diameter over time, locked to the stimulus) as a function of stimulus intensity in order to test whether it would show a similar pattern as the one observed in the brain. As shown in Figure 3C, this analysis revealed a clear intensity effect in the Active condition. The Passive condition showed signs of a similar effect, although noisier.

## 5. Networks associated with conscious and unconscious processing in the Active and Passive conditions

Although the aim of the present study is to overcome the classical procedure of contrasting conscious versus non-conscious processing using indirect labelling in the task-free condition, we also performed these classical contrasts to serve as a complementary analysis and a reference point before diving into a more agnostic description of single-trial dynamics.

### 5.1. Networks associated with non-conscious processing in the Active and Passive conditions

We first tested the network associated with non-conscious processing in the Active condition by contrasting trials reported as “Not heard”, when the stimulus was present at any intensity, with trials where the stimulus was absent (GLM3a; see Methods). In the Passive condition, we relied on the answers to mind-wandering probes (1/4^th^ of the trials) and contrasted trials where the mind-wandering content was anything but “The sound” (which we summarize by “Other”), when the stimulus was present at any intensity, and trials where the stimulus was absent (GLM3b; see Methods). In both the Active and Passive condition this contrast essentially showed bilateral activations of auditory regions (Figure 4A, Table 4 and supplementary Figures S8 & S9).

**Figure 4.**
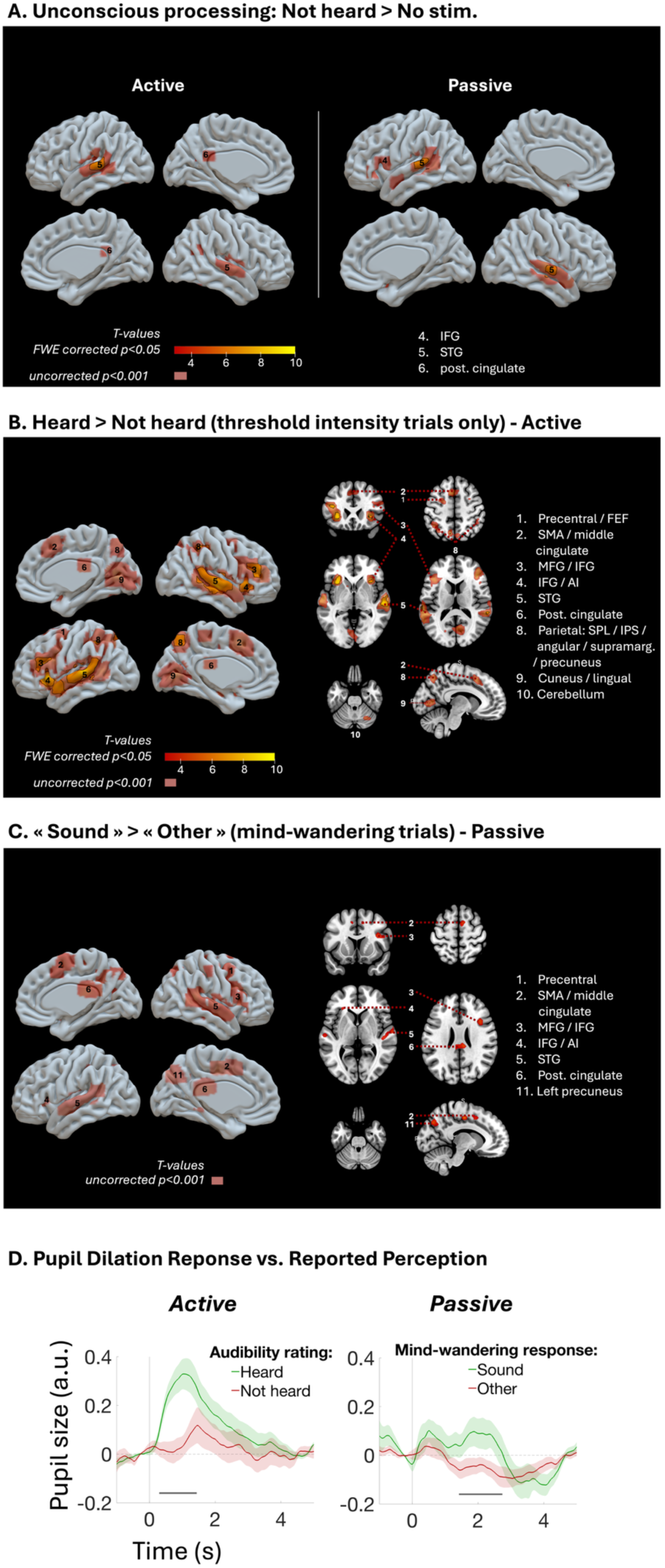
Contrasting (non-)conscious and stimulus absent conditions in the Active and Passive conditions: brain activations and pupil responses. A: T-maps contrasting "Stimulus present but not heard" with "No stimulus" in the Active and Passive conditions. B: T-map contrasting "Heard" (audibility ratings 2, 3 or 4) and "Not Heard" (audibility 1) in the Active condition, restricted to the trials at threshold intensities (based on behavioural data; 1/5^th^ of all trials). C: T-map contrasting trials with mind-wandering response "The sound" with other responses to mind-wandering probes (therefore limited to 1/4^th^ of all trials) in the Passive condition. Only uncorrected binarized t-maps are displayed for this contrast (p < 0.001).

**Table 4.** Regions associated with non-conscious processing in the Active and Passive conditions. The clusters and sub-clusters peaks listed here show the full results of the whole-brain analyses shown in Figure 4A; the sub-clusters that did not survive cluster-level FWE correction (p < 0.05) are displayed in grey and italics (p < 0.001). X, y and z coordinates refer to the Montreal Neurologic Institute (MNI) space. The last column indicates network(s) the clusters belong to, according to Yeo et al. 2011^42^ ; NA = Not applicable.

| ACTIVE - Not heard > No stimulus |  |  |  |  |  |  |  |  |  |  |  |
| --- | --- | --- | --- | --- | --- | --- | --- | --- | --- | --- | --- |
| Cluster label | Cluster size (voxels) | Side | Peak coordinates (mm) |  |  | Peak T-value | Z-score of peak T-value | Peak p-value (FWEc) | Peak p-value (unc.) | Cluster p-value (FWEc) | Network(s) of interest |
|  |  |  | x | y | z |  |  |  |  |  |  |
| Superior temporal gyrus | 336 | L | -58 | -28 | 8 | 8,06 | 5,44 | 0,002 | < 0.001 | < 0.001 | Auditory |
|  |  | L | -66 | -30 | 18 | 5,15 | 4,13 | 0,454 | < 0.001 |  |  |
| White matter | 44 | L | -24 | -2 | 20 | 5,71 | 4,42 | 0,191 | < 0.001 | 0,264 | NA |
|  |  | L | -24 | 8 | 23 | 4,67 | 3,85 | 0,77 | < 0.001 |  |  |
| Cerebellum | 18 | L | -24 | -48 | -47 | 4,96 | 4,02 | 0,576 | < 0.001 | 0,785 | NA |
| Superior temporal gyrus | 120 | R | 62 | -18 | 6 | 4,69 | 3,87 | 0,752 | < 0.001 | 0,01 | Auditory |
|  |  | R | 62 | -8 | 0 | 4,57 | 3,79 | 0,826 | < 0.001 |  |  |
| Cerebellum | 28 | R | 19 | -70 | -32 | 4,53 | 3,77 | 0,846 | < 0.001 | 0,542 | NA |
|  |  | R | 14 | -78 | -27 | 3,83 | 3,32 | 0,998 | < 0.001 |  |  |
| White matter | 14 | L | -26 | -20 | 20 | 4,4 | 3,69 | 0,907 | < 0.001 | 0,876 | NA |
| White matter | 5 | L | -24 | -52 | 23 | 4,11 | 3,5 | 0,982 | < 0.001 | 0,992 | NA |
| Cerebellum | 2 | R | 9 | -50 | -42 | 4,01 | 3,43 | 0,992 | < 0.001 | 0,999 | NA |
| Cerebellum | 9 | R | 46 | -62 | -27 | 3,92 | 3,38 | 0,996 | < 0.001 | 0,959 | NA |
| Angular gyrus / lateral occipital cortex | 4 | R | 42 | -60 | 26 | 3,88 | 3,35 | 0,998 | < 0.001 | 0,995 | DMN |
| Putamen | 1 | R | 29 | -15 | 3 | 3,74 | 3,25 | 1 | 0,001 | 1 | NA |
| Posterior cingulate | 3 | L | -6 | -45 | 23 | 3,73 | 3,25 | 1 | 0,001 | 0,998 | DMN |
| Cerebellum | 1 | L | -44 | -72 | -24 | 3,63 | 3,18 | 1 | 0,001 | 1 | NA |
| White matter | 1 | R | 32 | -15 | 18 | 3,6 | 3,16 | 1 | 0,001 | 1 | NA |
| Cerebellum | 1 | R | 14 | -60 | -37 | 3,53 | 3,11 | 1 | 0,001 | 1 | NA |
| PASSIVE - Other > No stimulus |  |  |  |  |  |  |  |  |  |  |  |
| Cluster label | Cluster size (voxels) | Side | Peak coordinates (mm) |  |  | Peak T-value | Z-score of peak T-value | Peak p-value (FWEc) | Peak p-value (unc.) | Cluster p-value (FWEc) | Network(s) of interest |
|  |  |  | x | y | z |  |  |  |  |  |  |
| Superior temporal gyrus | 289 | L | -64 | -32 | 8 | 6,97 | 5,01 | 0,019 | < 0.001 | 0,019 | Auditory |
|  |  | L | -61 | -48 | 8 | 5,04 | 4,06 | 0,492 | < 0.001 | 0,492 |  |
|  |  | L | -56 | -48 | 20 | 4,05 | 3,46 | 0,984 | < 0.001 | 0,984 |  |
| Superior temporal gyrus | 214 | R | 64 | -15 | -2 | 6,48 | 4,8 | 0,045 | < 0.001 | 0,045 | Auditory |
|  |  | R | 64 | -22 | 6 | 6,42 | 4,77 | 0,05 | < 0.001 | 0,05 |  |
| Temporal pole | 7 | R | 49 | 8 | -14 | 4,3 | 3,62 | 0,928 | < 0.001 | 0,928 | DMN |
| Inferior frontal gyrus | 27 | L | -48 | 18 | 8 | 4,26 | 3,6 | 0,942 | < 0.001 | 0,942 | DMN |
| Temporal pole | 12 | L | -48 | 0 | -17 | 4,11 | 3,5 | 0,976 | < 0.001 | 0,976 | DMN |
| Frontal pole | 7 | L | -44 | 32 | -4 | 4 | 3,43 | 0,989 | < 0.001 | 0,989 | DMN |
| White matter | 2 | R | 19 | -12 | 30 | 3,98 | 3,42 | 0,99 | < 0.001 | 0,99 | NA |

All slices are displayed in neurological convention. Both slices and 3D surfaces display corrected activations delimited by a black contour (FWEc, p<0.05), and uncorrected ones (p<0.001) in a less opaque shade. FEF = frontal eyefield; SMA = supplementary motor area; MFG = middle frontal gyrus; IFG = inferior frontal gyrus; AI = anterior insula; post. cingulate = posterior cingulate cortex; post. thal. = posterior thalamus; SPL = superior parietal lobule; IPS = intraparietal sulcus; supramarg. = supramarginal gyrus; STG = superior temporal gyrus. Exhaustive fMRI results are displayed in Tables 4, 5 & 6 and supplementary Figures S8-11. D: Pupil response evoked by the auditory stimuli as a function of awareness (using the same criterion as in the fMRI contrasts) in the Active (right panel) and Passive (left panel) conditions. Mean pupil diameter (z-scored) is plotted as a function of time relative to stimulus onset (t = 0s). Shaded regions indicate ± SEM across participants (N = 23 for Active; N = 25 for Passive). Horizontal markers at the base of each plot indicate time points where the pupil response was significantly different across responses (Paired t-tests, p < 0.05, uncorrected).

**Table 5.** Regions associated with the conscious versus non-conscious contrast in the Active condition. The clusters and sub-clusters peaks listed here show the full results of the whole-brain analyses shown in Figure 4B; the sub-clusters that did not survive cluster-level FWE correction (p < 0.05) are displayed in grey and italics (p < 0.001). X, y and z coordinates refer to the Montreal Neurologic Institute (MNI) space. The last column indicates network(s) the clusters belong to, according to Yeo et al. 2011^42^ ; NA = Not applicable.

| ACTIVE - Heard > Not heard at threshold |  |  |  |  |  |  |  |  |  |  |  |
| --- | --- | --- | --- | --- | --- | --- | --- | --- | --- | --- | --- |
| Cluster label | Cluster size (voxels) | Side | Peak coordinates (mm) |  |  | Peak T-value | Z-score of peak T-value | Peak p-value (FWEc) | Peak p-value (unc.) | Cluster p-value (FWEc) | Network(s) of interest |
|  |  |  | x | y | z |  |  |  |  |  |  |
| Superior temporal gyrus | 916 | R | 56 | -20 | -4 | 9,7 | 5,99 | < 0.001 | < 0.001 | < 0.001 | Auditory |
|  |  | R | 69 | -22 | 3 | 7,47 | 5,22 | 0,007 | < 0.001 |  |  |
|  |  | R | 62 | -38 | 13 | 7,12 | 5,08 | 0,016 | < 0.001 |  |  |
| Middle / inferior frontal gyrus / anterior insula | 1077 | L | -31 | 20 | 0 | 9,11 | 5,81 | < 0.001 | < 0.001 | < 0.001 | Salience / control / DMN |
|  |  | L | -41 | 25 | 18 | 8,17 | 5,49 | 0,002 | < 0.001 |  |  |
|  |  | L | -41 | 8 | 28 | 5,93 | 4,54 | 0,13 | < 0.001 |  |  |
| Superior temporal gyrus | 1159 | L | -68 | -28 | 6 | 9,08 | 5,8 | < 0.001 | < 0.001 | < 0.001 | Auditory |
|  |  | L | -54 | 5 | -12 | 7,37 | 5,18 | 0,009 | < 0.001 |  |  |
|  |  | L | -58 | -35 | 8 | 6,95 | 5 | 0,021 | < 0.001 |  |  |
| Superior parietal lobule / supramarginal gyrus | 186 | R | 42 | -40 | 43 | 8,4 | 5,56 | 0,001 | < 0.001 | 0,001 | Dorsal attention / control / visual |
|  |  | R | 36 | -50 | 33 | 4,56 | 3,79 | 0,828 | < 0.001 |  |  |
| Middle / inferior frontal gyrus / anterior insula | 773 | R | 42 | 32 | 18 | 8,01 | 5,42 | 0,002 | < 0.001 | < 0.001 | Salience / control / DMN |
|  |  | R | 32 | 20 | -2 | 7,35 | 5,17 | 0,01 | < 0.001 |  |  |
|  |  | R | 44 | 45 | 13 | 6,29 | 4,71 | 0,07 | < 0.001 |  |  |
| Superior parietal lobule / supramarginal gyrus | 465 | L | -38 | -40 | 40 | 7,71 | 5,31 | 0,005 | < 0.001 | < 0.001 | Dorsal attention / control / visual |
|  |  | L | -28 | -62 | 46 | 6 | 4,57 | 0,115 | < 0.001 |  |  |
| Precuneus | 143 | L | -8 | -68 | 53 | 6,84 | 4,96 | 0,026 | < 0.001 | 0,004 | DMN |
|  |  | R | 4 | -70 | 50 | 4,39 | 3,68 | 0,91 | < 0.001 |  |  |
|  |  | R | 14 | -70 | 43 | 4,02 | 3,44 | 0,99 | < 0.001 |  |  |
| Cerebellum | 73 | R | 26 | -75 | -50 | 6,68 | 4,89 | 0,034 | < 0.001 | 0,071 | NA |
|  |  | R | 39 | -68 | -50 | 3,77 | 3,28 | 0,999 | 0,001 |  |  |
| Superior frontal gyrus (SMA) / middle cingulate | 249 | L | -8 | 15 | 50 | 6,15 | 4,64 | 0,089 | < 0.001 | < 0.001 | Salience |
|  |  | R | 4 | 25 | 43 | 4,19 | 3,55 | 0,969 | < 0.001 |  |  |
| Cuneus / lingual / intracalcarine cortex | 691 | R | 2 | -72 | 6 | 5,96 | 4,55 | 0,123 | < 0.001 | < 0.001 | Visual |
|  |  | R | 6 | -65 | 13 | 5,57 | 4,35 | 0,239 | < 0.001 |  |  |
|  |  | L | -6 | -78 | 3 | 5,53 | 4,33 | 0,253 | < 0.001 |  |  |
| Cerebellum | 164 | R | 34 | -55 | -34 | 5,96 | 4,55 | 0,124 | < 0.001 | 0,002 | NA |
|  |  | R | 26 | -65 | -27 | 5,45 | 4,29 | 0,29 | < 0.001 |  |  |
|  |  | R | 24 | -58 | -34 | 4,43 | 3,71 | 0,893 | < 0.001 |  |  |
| Cerebellum | 71 | L | -26 | -78 | -47 | 5,8 | 4,47 | 0,164 | < 0.001 | 0,078 | NA |
|  |  | L | -34 | -75 | -50 | 5,43 | 4,28 | 0,297 | < 0.001 |  |  |
| Superior frontal gyrus (precentral / FEF) | 89 | L | -21 | -5 | 50 | 5,08 | 4,09 | 0,493 | < 0.001 | 0,036 | Control |
|  |  | L | -28 | 0 | 53 | 5,03 | 4,06 | 0,523 | < 0.001 |  |  |
| White matter | 31 | R | 32 | -65 | 6 | 5,04 | 4,06 | 0,522 | < 0.001 | 0,479 | NA |
| Caudate | 5 | L | -14 | 12 | -2 | 4,5 | 3,75 | 0,864 | < 0.001 | 0,992 | NA |
| Posterior cingulate | 21 | R | 9 | -25 | 23 | 4,42 | 3,7 | 0,898 | < 0.001 | 0,712 | DMN |
| Superior frontal gyrus (precentral) | 11 | L | -11 | -32 | 60 | 4,38 | 3,67 | 0,915 | < 0.001 | 0,931 | Somato-motor |
| Superior frontal gyrus (precentral) | 10 | R | 29 | 2 | 53 | 4,13 | 3,52 | 0,978 | < 0.001 | 0,946 | Somato-motor |
| White matter | 1 | L | -31 | -28 | 33 | 3,78 | 3,29 | 0,999 | 0,001 | 1 | NA |
| Cerebellum | 2 | R | 14 | -65 | -37 | 3,75 | 3,26 | 0,999 | 0,001 | 0,999 | NA |
| Putamen | 4 | L | -26 | -18 | 8 | 3,68 | 3,21 | 1 | 0,001 | 0,995 | NA |
| Caudate | 2 | R | 12 | 10 | 0 | 3,66 | 3,2 | 1 | 0,001 | 0,999 | NA |
| White matter | 2 | L | -21 | -58 | 0 | 3,56 | 3,13 | 1 | 0,001 | 0,999 | NA |
| White matter | 1 | L | -16 | 10 | 6 | 3,54 | 3,12 | 1 | 0,001 | 1 | NA |
| White matter | 1 | R | 46 | -40 | -10 | 3,53 | 3,11 | 1 | 0,001 | 1 | NA |
| White matter | 1 | R | 42 | -35 | -4 | 3,52 | 3,1 | 1 | 0,001 | 1 | NA |
| Superior parietal lobule / supramarginal gyrus | 1 | R | -46 | -38 | 56 | 3,51 | 3,09 | 1 | 0,001 | 1 | Dorsal attention / control / visual |

### 5.2. Networks associated with conscious versus non-conscious processing in the Active and Passive conditions

In the Active condition, the "Heard minus Not heard stimuli" at threshold intensity contrast was used to unravel the regions associated with conscious perception of the stimulus, at one fixed intensity level (Figure 4B, GLM 3a; see Methods). It revealed a network that was similar to the one found in the parametric modulation analysis for the same condition: a broad network of bilateral auditory, prefrontal, parietal, anterior insular, cerebellar and thalamic regions (Figure 4B, Table 5 and supplementary Figure S10). These findings partially survived multiple comparison correction although they were restricted to threshold intensity trials (i.e. 1/5^th^ of all trials, see Methods).

In the Passive condition, on the other hand, identifying the networks for conscious perception was more challenging, as statistical power was constrained by fewer probed trials (mind-wandering probe trials represented 1/4^th^ of all trials) and reduced signal-to-noise ratio compared to the Active condition. For this reason, all intensities were included in our first analysis contrasting answers "The sound" versus "Other" (GLM3c; see Methods). This mind-wandering - based contrast tentatively revealed a network encompassing bilateral PFC and anterior insular regions as in the Active counterpart although much weaker and less extended and, importantly, only in uncorrected statistics (Figure 4C, Table 6 and supplementary Figure S11).

**Table 6.**
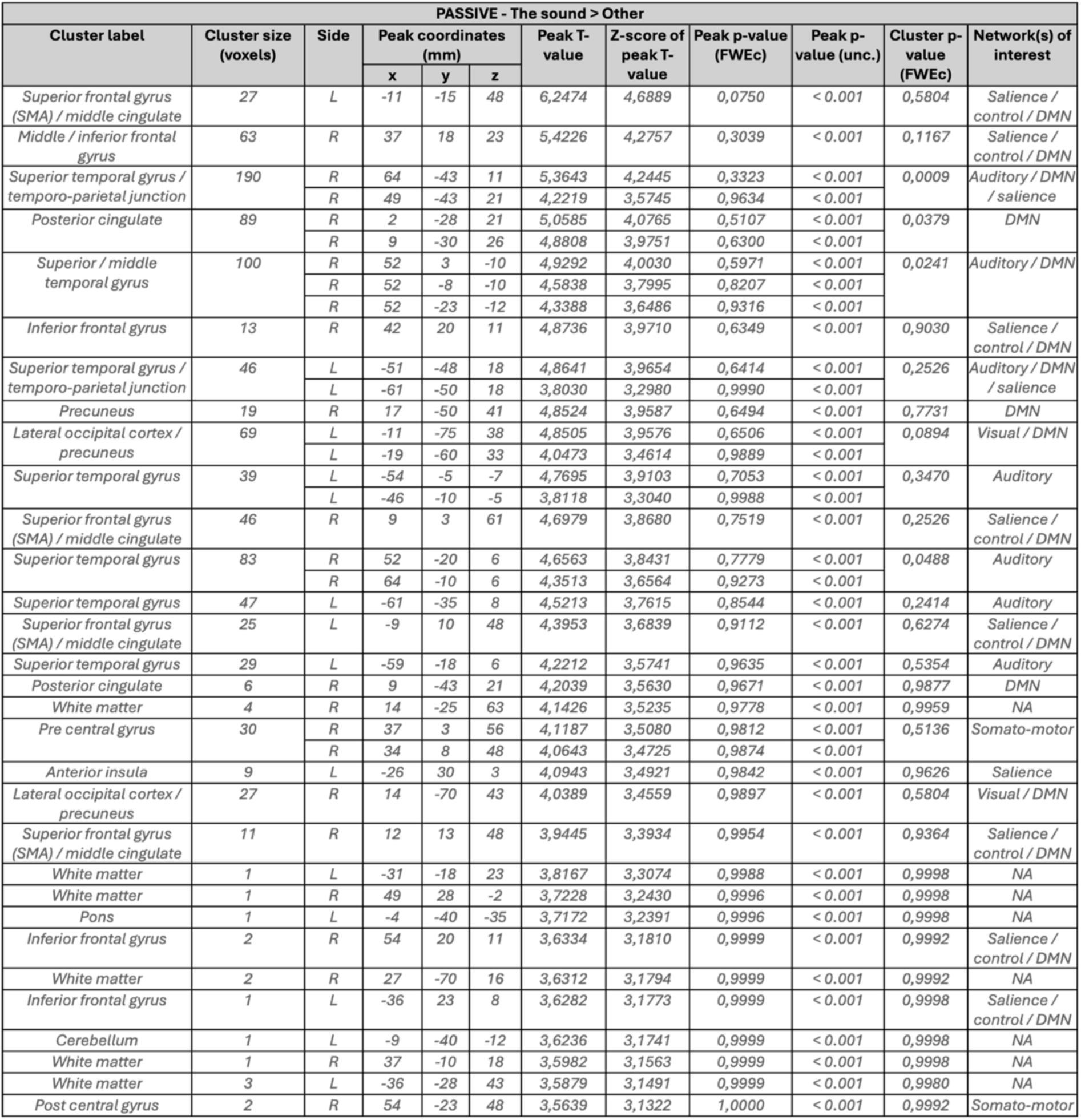
Regions associated with the conscious versus non-conscious contrast in the Passive condition. The clusters and sub-clusters peaks listed here show the full results of the whole-brain analyses shown in Figure 4C; the sub-clusters that did not survive cluster-level FWE correction (p < 0.05) are displayed in grey and italics (p < 0.001). X, y and z coordinates refer to the Montreal Neurologic Institute (MNI) space. The last column indicates network(s) the clusters belong to, according to Yeo et al. 2011^42^ unless specified otherwise; NA = Not applicable.

Since this contrast was performed with all the intensities, and the mind-wandering response correlates strongly with stimulus intensity, we next wanted to isolate the effect of conscious perception itself from the mere effect of stimulus intensity. We therefore built a composite model that included both the "Sound" / "Other" regressors for probed trials but also a stimulus intensity parametric modulation regressor to explain away the stimulus intensity effect (GLM3d; see Methods). With these exploratory contrasts, shown in Supplementary Figure S12, we found that extra-sensory regions encompassing PFC, cingulate and anterior insula were still observed in the “Sound” versus “Other” contrast after taking away the variance explained by the stimulus intensity. Conversely, we found strong auditory activations, with almost no extra-sensory regions, in the complementary parametric modulation contrast (see supp. Figure S12). This is in line with the main parametric modulation analysis (Figure 2B) revealing not only stimulus intensity effect, but also and mostly the conscious perception neural correlates.

## 6. Pupil size also reflects conscious report and mind-wandering responses

Finally, Figure 4D shows the pupil dilation response to the auditory stimuli as a function of awareness in the Active and Passive condition, as assessed with the same criterion than in the previous analyses. There was a marked increase of pupil dilation in "Heard" compared to "Not heard" stimuli in the Active condition, at threshold. Although less important, we also found a significant increase in pupil dilation mind-wandering trials with response "Sound" versus "Other" in the Passive condition (across all stimulus intensity), with a maximum difference between "Sound" and "Other" that seems to occur later than in the Active condition (around 2 versus 1 second(s) post-stimulus onset). These results obtained with an independent measure reinforce our fMRI observations that the stimulus is not processed in the same way across the Active and Passive conditions. They also provide very promising data for eye-tracking based markers of conscious auditory perception.

## 7. Diagnosing single-trial dynamics in primary auditory cortex and beyond

### 7.1. Distributions of single-trial activity in primary auditory and extra primary auditory Regions of Interest (ROIs)

As shown in Figures 1A & 5A, while average activity across trials can have very similar profiles between unimodal and bifurcation dynamics, these two types of dynamics can be clearly discriminated based on the distributions of activity across trials or on the profile of inter-trial variability as a function of stimulus intensity. For unimodal dynamics, average neural activity is expected to increase progressively with stimulus intensity with no or little change in inter-trial variability. For bifurcation dynamics on the other hand, when stimulus intensity is around threshold, it can lead to two types of equilibria: either activity remains at baseline, potentially corresponding to the absence of conscious access, or it leads to a higher equilibrium, potentially corresponding to conscious processing. The distribution of activity across trials is thus a mixture of two gaussians, in other words a bimodal distribution, with a low mode (baseline) and a high mode. As stimulus intensity increases, the probability of being in the "high" mode increases, until stimulus intensity is sufficient for the high mode to be reached on all trials (the stimulus is always conscious) (Figures 1A & 5A). This leads to a peak of variability around threshold intensity reflecting the spreading of the distribution of the values around threshold where the two gaussians are mixed.

With these two models in mind, we set out to characterize the trial-by-trial dynamics of neural activity within the brain-wide networks responding to the stimuli. More specifically, based on our previous EEG study^10^ we hypothesized that activity in the primary auditory regions, namely Heschl gyri, would primarily reflect initial sensory processing and thus display unimodal dynamics, while the rest of the network might reflect brain-scale activations associated with conscious access, and would therefore display bifurcation dynamics. The definition of the ROIs within and beyond primary auditory regions is described in the Methods section. We first performed a reduction of dimensionality to obtain, for each of these two ROIs (within or outside primary auditory regions), a single trial-by-trial neural value that summarizes the strength of the stimulus representation across the ROI. To do so, we used single-trial beta estimates (derived from a single trial GLM and using a dedicated toolbox, see Methods) to feed a linear multivariate pattern decoder trained on decoding maximal intensity stimulus versus no stimulus and tested on all intensity trials, following a leave-one-run-out cross-validation design, with the same logic that we used in our previous EEG study^10^. The decoding pipeline included an additional step of Recursive Feature Elimination to further select the most informative voxels (RFE; see Methods and supplementary Figure S15). The decoder provided one decision value (distance to the decision hyperplane) for each trial and ROI. These single-trial decision values are a proxy of how well the presence of the sound is represented in the ROI on each given trial; we next refer to them as "neural values".

As shown in Figures 5B & C, group-level distributions of trial-by-trial neural values revealed a marked difference across the two ROIs, both during the Active and during the Passive sessions. In the primary auditory regions, distributions across all intensity levels remained largely unimodal, with the mean response shifting incrementally as a function of stimulus intensity. In contrast, the other ROI, beyond primary auditory regions, exhibited a flattening of distributions at intermediate intensity levels. This structural deformation can be evocative of a mixture of two underlying modes - an "unconscious" and a "conscious" mode, rather than a single moving mean. The reason why one cannot expect a clearly bimodal aspect as presented in theoretical Figure 5A is that a group average of z-scored data with differing threshold intensity levels across participants (see individual behavioural data in supplementary Figure S13) blurs the two peaks of such bimodal distributions at threshold, if they exist. Next, we formally quantified this phenomenon using two independent metrics: the Variability Peak Ratio (VPR) and the Bifurcation Score (a model comparison metric based on the log-likelihood ratio of the two competing models).

**Figure 5.**
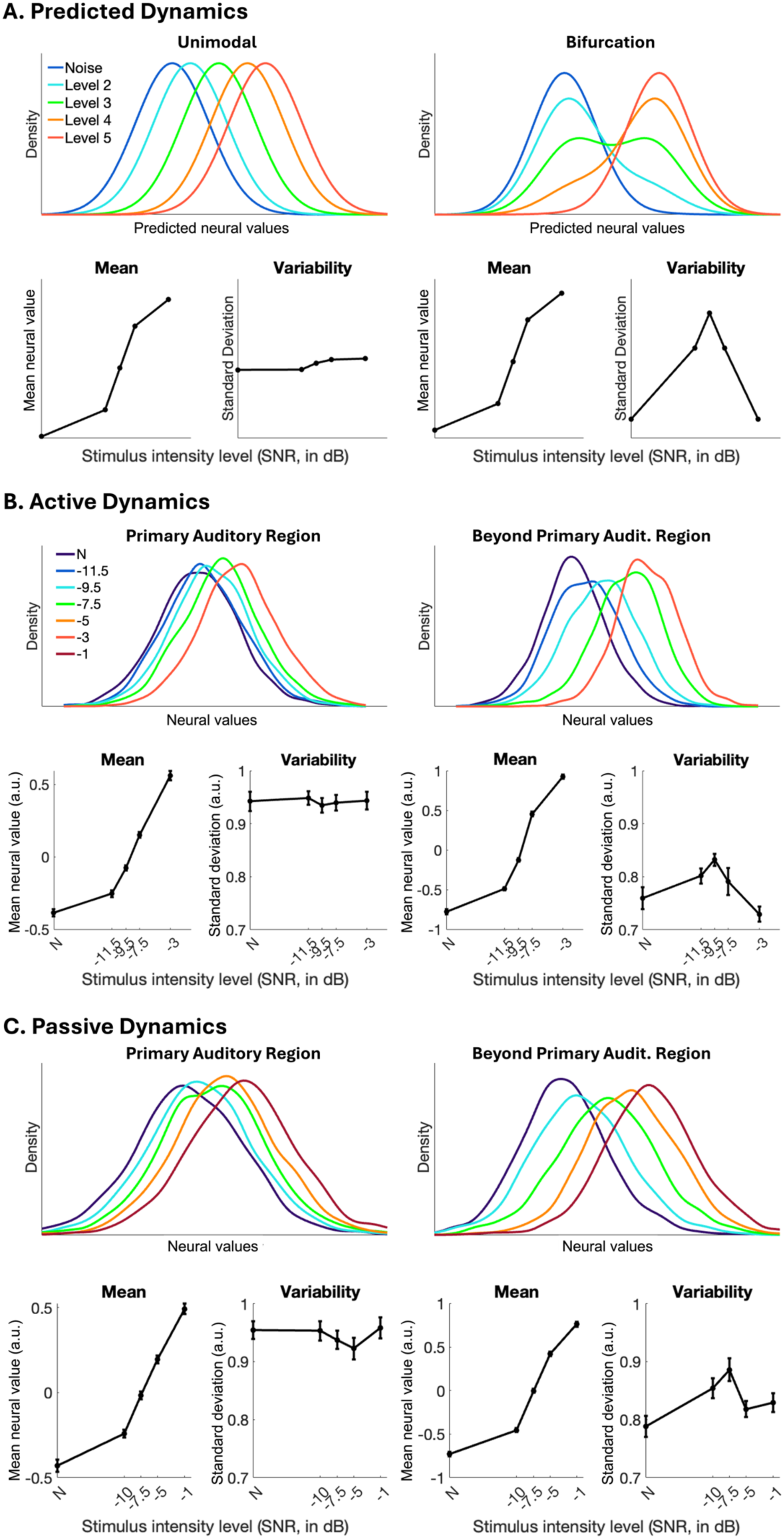
Dynamics of trial-by-trial neural values in primary auditory regions and beyond, in the Active and Passive conditions. A-C: Predicted dynamics of the unimodal and bifurcation models (A) and observed group-level dynamics of the neural values in the Active and Passive conditions, within versus beyond the primary auditory regions (B-C). Top panels display single-trial neural values distributions, color-coded as a function of stimulus intensity, and bottom panels display the profiles of average (on the left) and inter-trial variability (on the right) of the neural values as a function of stimulus intensity (values are in dB; N stands for Noise (SNR level 1)).

### 7.2. Formally testing bifurcation in single-trial data

#### The Variability Peak Ratio (VPR)

As a first approach, we tested the variability profiles (Figure 6A), predicting that bifurcation dynamics are associated with a peak of inter-trial variability around threshold intensity. The Variability Peak Ratio, or VPR, measured the ratio between the inter-trial standard deviation at intermediate intensity levels and the inter-trial standard deviation for the extremes, namely no stimulus or stimulus at maximal intensity (see Methods).

**Figure 6.**
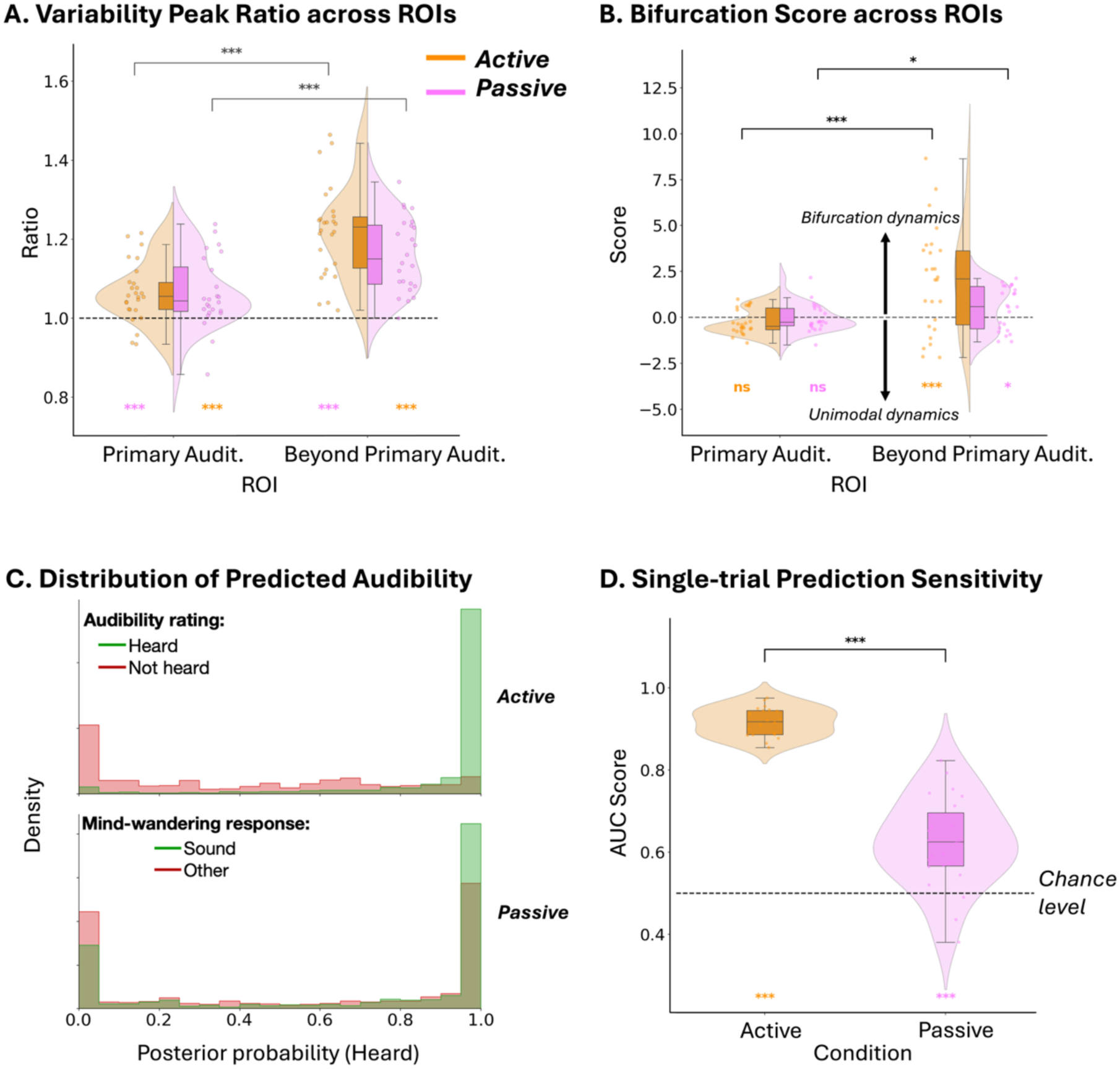
Single-trial dynamics show a bifurcation beyond the primary auditory regions and predict single-trial conscious report or mind-wandering content. A: Variability Peak Ratio (VPR; N = 27 participants) compared across both regions of interest and conditions. B: Bifurcation score (N = 24 participants) compared across both regions of interest and conditions. C: Group-level distributions of single-trial posterior probabilities derived from the bifurcation model, categorized by participants’ behavioural reports, displayed for the Active and Passive conditions, respectively. Green shaded areas represent trials where participants reported "Heard" or "Sound"; red shaded areas represent "Not Heard" or "Other". Data are pooled across all 20 participants included in the analysis and across all stimulus intensities. The x-axis represents the model’s confidence in a "Heard" state (0: "Predicted not heard"; 1: "Predicted heard"). D: Sensitivity for predicting single-trial conscious report (Active) or mind-wandering content (Passive) based on single-trial activity beyond the primary auditory cortex and the bifurcation model (AUC score; N = 20 participants). For panels A, B & D: in split-violin plots, the shaded areas represent the kernel density estimation of the underlying distribution for the Passive and Active conditions; in box plots, the central horizontal line indicates the median, while the box represents the interquartile range (IQR). Whiskers extend to the furthest data points within 1.5 * IQR. Each dot represents the score for an individual participant, jittered horizontally for visibility. Wilcoxon tests results for values against null are displayed with coloured asterisks or "ns" below the plots (*** p < 0.001, ** p < 0.01, * p < 0.05, "ns" non-significant). Dotted horizontal lines represent respectively "null" VPR value (1), null performance of the Bifurcation score (0) and AUC score chance level (0.5). Comparison tests (paired Wilcoxon signed-rank tests, one-tailed for B & D, two-tailed for A): the horizontal brackets spanning across ROIs or conditions indicate the results of comparisons (within condition, across ROIs for A & B, across conditions only for beyond primary auditory ROI for D). The asterisks above these brackets represent significant differences (*** p < 0.001, ** p < 0.01, * p < 0.05, "ns" non-significant).

Figure 6A shows that VPR was, as expected, significantly higher beyond primary auditory regions than in the primary auditory regions for both Active and Passive conditions (Wilcoxon tests p = 1.10^-7^ for Active condition, p = 0.00074 for Passive condition). However, VPR was significantly greater than 1 in both ROIs and both conditions, suggesting that there might be a deviation from purely gradual and unimodal dynamics also in the primary auditory region (primary auditory region: p = 0.00022 for Active condition, p = 0.00049 for Passive condition; beyond primary auditory region: p = 1.10^-8^ for Active condition, p = 1.10^-7^ for Passive condition).

#### Modelling the single-trial distributions and estimating the Bifurcation Score

We next formally tested the dynamics underlying the full distributions of neural values across trials by fitting the unimodal and bifurcation models to these empirical data and comparing their respective performance across both conditions and both ROIs. The equations and parameters of these models are detailed in the Methods section (equations 4, 5 and 6): the unimodal model reflects a progressive increase of neural value as a function of stimulus intensity, given by a unimodal distribution centred on a mean increasing with intensity, while the bifurcation model represents a Gaussian mixture of a low and high distribution, with the mean of the high distribution and the probability of being in the high distribution increasing with stimulus intensity.

The Bifurcation Score measured the relative goodness of fit of the bifurcation versus unimodal model based on their respective performance (log likelihood) measured in a leave-one-run-out cross-validation design and taking into account each model’s gain over the null model (see Methods and Figure 6B). In both the Active and Passive conditions, the Bifurcation score confirmed the existence of significant bifurcation dynamics in the regions beyond primary auditory cortex. In the Active condition the Bifurcation score beyond primary auditory regions was significantly higher than 0 (p = 0.0008, mean = 1.991, median = 2.09806, SD = 2.848;) and significantly higher than in primary auditory regions (p = 0.0004). In the Passive condition, the Bifurcation score beyond primary auditory regions was also significantly higher than 0 (p = 0.0228, mean = 0.520, median = 0.58885, standard deviation, SD = 1.178) and higher than in primary auditory regions (p = 0.0323).

Although the bifurcation score in primary auditory regions was not significantly lower than 0 (p = 0.07 in the Active condition, p = 0.3317 in the Passive condition), it reached a negative value in both conditions (in the Active condition: mean = -0.25, median = -0.477, SD = 0.681; in the Passive condition: mean = -0.034, median = -0.261, SD = 0.759), hence in favour of the unimodal model in that ROI.

#### Predicting report based on the single-trial bifurcation model

Finally, since the bifurcation model accounted well for the trial-by-trial dynamics in neural value beyond the primary auditory cortex, we tested if the split between two types of trials that it implies corresponded to the difference between conscious and unconscious processing assessed either by direct conscious report in the Active condition, or mind-wandering the Passive condition. Based on the fitted parameters of the bifurcation model for each participant, we computed the posterior probability of each single-trial neural value in the beyond primary sensory ROI, to belong to the high distribution (*x*; see Methods). Finally, we estimated whether these posterior probabilities correctly predicted conscious processing by comparing their distributions in the “Heard” versus “Not Heard” trials in the Active condition, and in the “Sound” versus “Other” mind-wandering contents in the Passive condition. Figure 6C shows these distributions at the level of the group. These posterior probabilities seemed to predict “Heard” versus “Not heard” trials very well in the Active condition, as attested by mostly low predictions for “Not heard”, and high predictions for “Heard”. In the Passive condition we also observed a clear separation, although not as neat as in the Active condition. This was to be expected for two reasons: first these trials only represent 1/4^th^ of all the trials, and are thus nosier, and second, it is possible that when participants reported “other” mind-wandering contents, they also had the sound in their mind but as a less salient content. In order to quantify the prediction performance, we calculated the Area Under the Receiver Operating Characteristic Curve (AUC-ROC) between the two distributions ("Heard” versus “Not heard” and “Sound” versus “Other”). This measured the sensitivity of the prediction (Figure 6D). Prediction sensitivity (AUC) was significantly higher than chance level in both Active and Passive conditions (Figure 6D; Active condition: mean AUC = 0.916, median = 0.918, SD = 0.035, p = 1.10^-6^; Passive condition: mean AUC = 0.623, median = 0.626, SD = 0.108, p = 1.10^-5^). These results suggest that the bifurcation observed in trial-by-trial neural values beyond the primary auditory cortex reflects the separation between conscious and non-conscious processing independently of the task.

## Discussion

A central difficulty in studying consciousness in the absence of report is the reliance on indirect markers of conscious perception. Here we invert the usual logic by assessing whether different ways of processing the same stimulus emerge naturally from the brain’s response to near-threshold stimulation and then test whether these dynamically defined modes correspond to the presence or absence of conscious perception when probed. As a first step, we used parametric modulation to delineate the networks whose activity scaled with stimulus intensity. This revealed a clear set of common areas at the conjunction between task-related and task-free networks, that spans not only sensory regions but also prefrontal, insular and cingulate regions (Figure 3A). Interestingly, while the BOLD temporal response profiles were very similar across conditions in the auditory regions, they displayed important differences in extra-auditory regions (Figure 3B), a pattern that could explain why some previous results might have appeared contradictory, as discussed below. The contrast between “Heard” and “Not heard” trials in the Active and Passive conditions, using mind-wandering probes for the latter, highlighted consciousness-related networks that were remarkably consistent with the conjunction network revealed by parametric modulation. However, this contrast was less sensitive due to the fact that mind-wandering probes concerned only ¼^th^ of the Passive trials. Hence, we adopted the reverse approach: we characterized the dynamics of trial-by-trial activations within the networks responding to the stimuli, distinguishing sensory and higher-level parts. This revealed that, in both Active and Passive conditions, primary sensory regions exhibited unimodal dynamics, while the rest of the network showed bifurcation dynamics, suggesting two possible modes of activation across trials for the same stimulation around threshold, irrespective of task presence. Finally, the probability of each single-trial activity to belong to one mode or the other accurately predicted trial-by-trial conscious experience as probed either by direct report or mind-wandering in the Active or Passive sessions respectively.

As discussed in detail below, these results suggest the existence of a shared network between task-related and task-free conscious perception, distinct both from sensory networks and from decision-making networks, and whose all-or-none engagement correlates with conscious perception or its absence irrespective of task demands. We discuss these results in relation to the global neuronal workspace model and argue that, while they are compatible with this framework, they advocate for an update that includes the notion of a global playground, i.e. a brain-wide network that shares critical nodes with the global workspace but can be disconnected from executive networks.

### A common network between task-related and task-free auditory processing

Thanks to the very good sensitivity and specificity of the parametric modulation approach, we could formally test the conjunction between networks associated with task-related and task-free processing, a direct statistical contrast that very few other studies on the topic actually perform (see Kronemer and colleagues 2022^30^ for another example using a visual protocol). This analysis clearly demonstrated that this common network includes both sensory and extra-sensory regions, and that the difference essentially relies on an extinction of the areas involved in executive and motor control during task-free processing (Figure 3A).

Interestingly, the temporal part of this intersection not only included the primary auditory cortex (Heschl’s gyrus) but also the lateral sulcus and superior temporal gyrus (STG). This seems quite remarkable given the simplicity of the stimuli and their subtle intensities compared to the surrounding noise, especially in the Passive condition where they should not benefit from top-down attention. Furthermore, some temporal activations extended beyond the STG to the middle temporal gyrus, typically belonging to the default mode network (DMN)^42^. This matches the observations described in another study investigating conscious auditory perception^5^. Posterior to the broad STG activations, we also found what could correspond to the temporo-parietal junction (TPJ), although it is an inter-individually varying region that is difficult to rigorously identify in pooled 3T fMRI data and is not defined in common atlases. Along with anterior insular and cingulate regions, TPJ stands at the junction between salience and default mode networks^42,44,45^ and is typically involved in detecting salient events in the sensory environment in both task-dependent and task-independent settings^46^.

Beyond the temporal cortex we found conjunction effects in anterior insula, middle and inferior frontal gyri (MFG and IFG) and middle cingulate cortex (Figure 3A). When putting these different areas in correspondence with the resting-state networks identified by Yeo et al.^42^, these extra-sensory areas seem to correspond to convergence zones between the salience network, the fronto-parietal control network and the DMN. Compared to the network observed in Active sessions, some key areas of the fronto-parietal control network simply cease to respond to the stimulus in Passive sessions, confirming several previous observations^5,13,26,28,31^: the more anterior part of the MFG, the more anterior part of the middle cingulate cortex as well as the intra-parietal sulcus and surrounding areas. Likewise, activations in the left motor cortex corresponding to response preparation with the right hand are extinguished in the Passive session and therefore absent from the conjunction.

Importantly, the temporal response profiles of the BOLD signal within these regions were very different according to task, contrasting with what we found in superior temporal regions (Figure 3B). In the Passive sessions, they showed a relative disengagement during the periods where the vowels were expected to be presented compared to the other periods of the same sessions, where participants were actively engaged with other tasks. Nevertheless, and crucially, they did not cease to respond to the stimulus: as stimulus intensity level increased, this relative disengagement was progressively lifted, explaining the strong parametric modulation effect within these regions even in the Passive condition.

Finally, we found bilateral medial occipital activations specific to the Active condition: this is probably due to a task-preparation effect, participants indeed expected a response screen to appear shortly after hearing the stimuli. It has been described in other studies exploring task-relevant auditory stimuli with similar experimental designs involving a change of visual screens sequentially after stimulus presentation and / or detection^4,7,13,47^. Beyond mere anticipation of the response screen, this occipital activation could also be due to a more general cross-modal effect with visual cortex activity being modulated by auditory stimuli in a task-relevant setting^48^.

### Resolving previous contradictions on extra-sensory activations to task-irrelevant stimuli

These results could help resolve apparent contradictions in previous studies and inform debates about the involvement of extra-sensory areas, notably the Prefrontal Cortex (PFC) in task-free conscious perception^32,49^. Many studies show that stimuli that do not have to be reported are associated with decreased activity in several areas of the PFC relative to task-relevant ones^26,28^. This has been taken to suggest that prefrontal activity might be exclusively related to post-perceptual processes such as decision making^24,32^. Here we show that a relative decrease of activity within these regions during stimulus presentation in task-free situations is not incompatible with them actually processing the stimulus: there is a relative change in baseline activity, but the area remains responsive to the stimulus. Furthermore, our results highlight the fact that the PFC is not a monolithic structure, as they establish a clear distinction between areas that cease to respond to the stimulus in task-free settings (e.g. anterior parts of the MFG) and other areas that respond to the stimulus regardless of whether it has to be reported (posterior IFG/MFG).

Another issue in this debate relates to the observation that, even when some prefrontal regions are shown to respond to the stimulus in task-free situations, these activations are typically weaker than in sensory areas^5,13,28,29^. This is also what we observe here: there is a clear modulation of the activity in extra-sensory ROIs by our stimuli even when they are task-irrelevant, but this modulation is attenuated compared to when they are task-related; in comparison, the attenuation seems less important in sensory regions (Figure 3B). In line with this, statistical significance is often lost for fine contrasts between conscious and unconscious processing in the absence of a task, and more sensitive types of analysis are often requested to uncover these activations^5,29^. According to some authors, this difference in intensity and extent of activation suggests that the role of the PFC and associated extra-sensory regions might be minor^5,25,29^. However, other elements can explain these discrepancies.

First, some aspects of the experimental protocol or analysis might undermine statistical power, such as comparing two different groups of participants in inattentional blindness protocols^5^. A particular drawback of these types of protocols, which are nevertheless central for studying task-free conscious perception, is that the “aware group” does not perceive the stimulus consciously on all trials, as verified in follow-up report performed after sessions, hence reducing the contrast with the “unaware group”. Furthermore, different types of distracting tasks may have various impacts on the processing of task-irrelevant stimuli, and might sometimes drastically reduce the chance that they become conscious^13^, especially when the stimulus of interest and the distracting task are in the same modality^5,29^. This highlights another advantage of the current approach, since we tested a range of different stimulus intensities and could directly anticipate and measure the shift in threshold across the Active and Passive conditions.

Second, these differences might also be inherent to the different ways of encoding stimuli along the gradient from sensory to extrasensory areas. Indeed, the PFC is a highly integrated network that is involved in many cognitive processes. While it is traditionally associated to behaviour guiding based on rules, task, environment and context^50^, we also know that the PFC encodes perceptual information such as visual category regardless of task relevance, and does so in an abstract way in the sense that it generalizes across modalities^51,52^. The PFC is highly connected to other brain areas upon which it exerts a top-down influence based on the context^53^. Finally, it plays an important role in information maintenance in working memory^53^. This vast region therefore not only underlies executive, task-related functions, but also plays a central role in integration and maintenance of sensory information, which are hallmarks of conscious access. The role of the PFC in such diverse yet concomitant high-level cognitive functions, reflects in the versatility of its encoding, and the higher variability of its organization across individuals^54^. This might underlie the greater difficulty to isolate its precise involvement compared to sensory areas, especially when performing subtle contrasts or comparing across participants. Indeed, even multivariate approaches have been described as weaker for decoding content in the PFC than in other brain regions using fMRI^55^. Accordingly, Hatamimajoumerd and colleagues pointed that they could decode visible versus masked trials above chance in the frontal lobe in both report and no-report conditions, although with weaker decoding performances in the no-report condition and with no cross-condition decoding across report and no-report due to a different encoding of stimuli in the PFC as a function of task relevance^29^.

In conclusion, along with these considerations our current results can help resolve what might have been previously interpreted as contradictory findings. Indeed, beyond clear divergences in interpretation, the patterns of activations are actually remarkably coherent across studies, even in extra-sensory areas: significant activations are found in the anterior insula and the IFG / MFG in response to the presence of task-irrelevant stimuli, be they auditory^5,13^, visual^25,27–30^ or even somatosensory^56^, even in studies which argue against the involvement of the prefrontal cortex in task-free perception^13,28^. Several studies also report middle / anterior cingulate activations to be shared between task-related and task-free conditions^5,30^. Finally, we also found precuneus activations in both task-related and task-free stimulus processing (Figure 3A), although not in overlapping regions, which likewise matches some previous observations^5,30^. These observations strongly argue in favour of the idea that conscious perception involves a brain-wide network encompassing both sensory and extra-sensory areas even when the stimulus is not relevant for a task.

### The global workspace and the global playground

These conclusions are in line with the central idea behind the global neuronal workspace framework^15,19,57,58^, which proposes that conscious perception specifically involves broadcasting sensory representations beyond the automatic routes of sensory processing, allowing them to be maintained and have an impact within a vast range of other functional networks. They also call for an important update relative to previous formulations of the global workspace. Indeed we show, in accordance with previous evidence, that the typical fronto-parietal control network and dorsal attention network^42^ that were highlighted as key areas of the global neuronal workspace are actually, in large parts, dedicated to the task and cease to respond when the stimulus is not task relevant. Importantly though, we demonstrate that other parts of the prefrontal, insular and cingulate cortices delineate an extra-sensory network that is shared between task-related and task-free processing, hence corresponding to the tentative neural correlates of conscious access independent of a task. This brings direct empirical support to the proposition that spontaneous conscious perception in the absence of a task might rely on a global playground, namely a wide network of sensory and extra-sensory areas within which the information is maintained and circulated, despite being decoupled from executive and motor functions^10^. Interestingly, the shared nodes we observed, at the conjunction of these tentative global workspace and global playground (Figure 3A) seem to stand at the cross-roads between several networks: default mode, salience and control networks^42^. We propose that they might correspond to essential nodes for the broadcasting process advocated by the model. In favour of this interpretation, we observe that they overlap with regions identified as important hubs or “rich-club” areas based on their particularly rich functional connectivity to the rest of the brain^59–61^.

Within the global playground, we could also expect to find activations that are unique to this more spontaneous way of processing the stimulus, beyond the shared hubs. One such “playground-specific” activation was found in the inferior part of the left precuneus. Unlike lateral posterior parietal regions, which are involved in visuospatial cognition and body movement control, the precuneus could be involved in a broader range of high-level cognitive functions including not only visuo-spatial processing but also episodic memory retrieval (including in the auditory modality), self-processing, and the default mode network^62–64^. However, such playground specific activations remain particularly modest here, which could be attributed to the fact that, when not relevant to the tasks, our vowel stimuli were not particularly rich or interesting. We might expect more interesting task-irrelevant stimuli, for example musical stimuli or more varied linguistic stimuli, to evoke broader activations within the DMN.

#### Bifurcation dynamics in extra-sensory areas underlie the difference between conscious and unconscious processing independently of task

Once the task-independent networks responding to the stimulus have been clearly identified, there remains the central question of understanding their specific role in the contrast between conscious versus non-conscious processing. This is where the most innovative aspect of our approach comes into play: not only did we use inferred labels of conscious versus non-conscious processing to perform this contrast, we also reversed the logic: we tested whether bifurcation dynamics emerge from the activity of some parts of the network, reflecting different ways of processing the same stimulus even in the absence of a task, and then tested whether they matched conscious experience when identified.

So far, studies on task-free conscious perception have relied on different indirect ways to label the processing as being conscious or not. As mentioned earlier, one popular way is to use inattentional blindness and perform a contrast between “aware” and “unaware” participants. Another original approach has been proposed by Kronemer et al^30^, who labelled task-free trials based on pupil data using a classifier trained on task-related trial, with the drawback that this possibly contaminates the decoding features with task-associated processes. We also tried a similar approach, by using responses to mind-wandering probes to assess whether participants had the task-irrelevant sound on their mind or not. This is an immediate single-trial measure, but it comes with the strong limitation that it only concerns one quarter of the trials, hence drastically reducing statistical power. Still, using them to perform contrasts confirmed that unconscious processing of the auditory vowels was essentially restricted to auditory regions in both Active and Passive sessions (Figure 4A), which is consistent with previous literature showing essentially sensory activations to unconscious stimuli^5,16^. When contrasting conscious versus non-conscious stimuli we found the same extra-auditory regions identified in the conjunction analysis, including PFC, anterior insula and middle cingulate cortex, even in the absence of a task and even when removing the stimulus intensity effect (Figures 4B & C and S12). However, statistical significance did not survive correction for multiple comparison in Passive sessions where we could only use ¼^th^ of the trials.

Inverting the logic by analysing the dynamics of activations across trials allowed overcoming this caveat. This builds on previous work with time-resolved neuroimaging techniques suggesting that conscious processing might be specifically associated with the all-or-none triggering of late activations, beyond the initial stages of sensory processing, typically after 250 milliseconds post stimulus^18,35–37,65^. Sergent and colleagues (2021) further demonstrated that these specific dynamics could be found even in the absence of report or task on the stimulus, and could be directly read out from brain activity, with no external conscious versus non-conscious labelling and no priors on the type of neural activity involved^10^. This approach leverages the distribution of activity across trials, either with simple measures of intertrial variability, or using model fitting and model comparison. Here we deployed this technique in fMRI using single-trial betas. While extracting single-trial betas in fMRI has been described for several types of analyses, including EEG-fMRI coupling or implementing multivariate pattern analyses ^43,66–72^, no previous study to our knowledge has examined how these values are distributed across trials and how these distributions might change with experimental conditions. Here we examined these distributions to better understand the dynamics of activation within the networks responding to the stimuli. Note that the region selection was based on parametric modulation analyses that were orthogonal to the specific single-trial dynamics under investigation. Although the temporal resolution of fMRI is very limited, we reasoned that the dynamics of activity within the primary auditory cortex might essentially reflect the initial stages of sensory processing, and hence follow unimodal dynamics, while the bifurcation dynamics observed at later stages in time-resolved recording techniques might be due to the all-or-none triggering of joint activations within a broader network, encompassing the extra-sensory nodes. The distributions of single-trial activations within the primary auditory region versus beyond primary auditory regions seemed to validate this prediction. Indeed, they displayed a clear broadening of the distribution for threshold intensities compared to extreme intensities specifically for regions beyond the primary auditory cortex in both Active and Passive conditions (Figures 5B & C), which might reflect a mixture of gaussians instead of a single gaussian at threshold stimulation. This was also well reflected in a peak of inter-trial variability specifically for stimuli at threshold (Figures 5B & C). The Variability Peak Ratio allowed quantifying this characteristic peak at threshold and confirmed that this signature of bifurcation was present beyond the primary auditory cortex in both Active and Passive sessions and significantly differing from primary auditory cortex (Figure 6A). Finally, we directly fitted the unimodal versus bifurcation models to the observed distributions. Model comparison confirmed that bifurcation dynamics were a better model than unimodal dynamics to explain activity in areas beyond the primary auditory cortex (Figure 6B). This validated our proposition that, for stimulation at threshold, areas beyond the primary auditory cortex might sometimes be involved and sometimes not across trials.

The next step was to test whether this differential involvement corresponded to the difference between conscious and non-conscious processing using an external behavioural benchmark. We tested how well the bifurcation model could predict single-trial conscious report based on the activity of the beyond primary auditory network during that trial. The bifurcation model provided an excellent trial-to-trial prediction of conscious report in the Active condition. In the Passive condition, while its performance was lower than for the Active condition given the fewer number of trials, it also predicted significantly better than chance whether, when prompted with a mind-wandering probe, participants would report that they had the sound on their mind versus anything else (Figures 6C & D).

In conclusion, these results suggest that regions responding to the stimuli in both Active and Passive settings, beyond the primary auditory cortex, form a key network that operates the switch between conscious and unconscious processing independently of report or task.

### Towards understanding the brain mechanisms of conscious access independent of task

The present results converge with other results obtained with time-resolved neuroimaging techniques, in our previous EEG study^10^, but also replicated in the visual domain by Cohen et al.^73^. They can help us envision the scenario of brain mechanisms leading to conscious access in the general case. Here we propose an interpretation within the global workspace framework, that includes the notion of a “global playground” for task-free conscious processing. We propose that the first stages of cortical auditory analysis, in the primary auditory cortex, are faithful to the physical parameters of the stimulus, scaling with its intensity in a unimodal manner, even for stimulation at threshold (Figures 5A & B). Such faithful encoding might propagate automatically along the hierarchy of auditory processing up until 200-300 milliseconds which is the critical period where we observe the change of dynamics for later activations with time-resolved techniques^10,18,36,73,74^. This critical period might correspond to the moment where this information reaches attentional nodes and competes with other stimuli for potential attentional amplification. We tentatively propose that the inferior frontal gyrus and anterior insula in particular might play a pivotal role in the engagement of the rest of the network, in line with the view of the insula as a "multimodal gate" for conscious processing^75,76^. Other studies using intracranial recordings suggest that some subcortical arousal regions such as the intralaminar thalamus might also play such a role^4,9,30^. In the present study there are some indications of thalamic activations in relation to stimulus processing (see label 7 in Figure 2A), but whole brain fMRI is probably not the best technique to reveal such subcortical activations. We propose that, at this point, the same near-threshold stimulus might, across spontaneous fluctuations of the system, follow two possible paths. It might either fail to gather sufficient attentional resources to engage a broader network, in which case its processing will remain unconscious and will essentially be confined within sensory areas (Figure 4A), or alternatively it might gain sufficient momentum within these gating areas to successfully engage a broader network. This would create a new equilibrium, reflected as a dynamical bifurcation, in which this specific sensory information is enhanced and stabilized across a broader network that includes both auditory and extra-auditory areas through reciprocal loops. We propose that the shared network outlined in Figure 3A might constitute a core set of nodes enabling this new equilibrium and that this core network might then flexibly connect different functional systems depending on current demands: either routing processing towards executive areas if a task is requested, or alternatively diffusing it towards the DMN, notably the inferior precuneus.

### The pupil reflects these processes and is a promising marker for future clinical use

Throughout these analyses we were also interested in testing whether the pupil reflected these complex brain-scale dynamics and could thus serve as a proxy potentially transferable to clinics. In fact, there have been reports that a pupil response can be elicited by auditory stimuli not only in healthy volunteers^77–79^ but also in patients with disorders of consciousness^80,81^. Such a simple marker would be very useful to help the diagnosis of these patients who show clinical signs of wakefulness but do not communicate reliably and clinical examination alone is often insufficient to establish whether a patient is self-aware or aware of their environment^82,83^. However, such auditory pupil response studies have largely focused on "oddball" paradigms and, while a recent study reported that task-irrelevant auditory stimuli could evoke a pupil response^84^, this was not related to conscious versus non-conscious processing.

Here we observed that auditory stimulation evoked a pupil dilation response that was modulated by both auditory intensity and perceptual report in both Active and Passive conditions (Figures 3C & 4D). The fact that the ’Heard’ vs. ’Not heard’ distinction was clearly and significantly reflected in the pupil response even during passive listening is particularly remarkable. This suggests that pupillary response to auditory stimulation around threshold might be a very promising physiological marker that mirrors the internal state of the listener, even in the absence of overt behavioural report. Further work could pursue these investigations by probing single-trial dynamics of the pupil dilation responses, which might reflect the same bifurcation dynamics as the brain activity, and could thus be used to probe signs of these dynamics in patients with disorders of consciousness with a simple recording system.

### Limitations and perspectives

The Heard versus Not heard comparisons should be interpreted with caution. By design, these contrasts rely on trial sorting based on subjective report and therefore lead to relatively small and potentially imbalanced trial numbers, espceially in the Passive condition. This limits statistical power and may increase susceptibility to noise-driven effects. Our new approach paves the way for potentially overcoming this caveat, as our model allows to estimate on each trial the probability that the stimulus was heard even when no mind-wandering probe was presented.

While our current analysis focused on specific ROIs, a primary goal for future refinement is to implement this framework with higher spatial precision. Moving toward a searchlight-based or voxel-wise bifurcation mapping would allow us to trace the emergence of bifurcation distributions across the entire cortical hierarchy with finer granularity, potentially pinpointing the regions where a sensory representation becomes a conscious mental content.

## Conclusion

In conclusion, rather than starting from predefined categories of conscious and non-conscious processing, we inverted the usual logic by examining the brain’s response function around perceptual threshold and the fundamental dynamical principles governing its activity. To this end, we developed a modelling framework capable of characterizing the distribution of single-trial neural responses and identifying bifurcation dynamics in fMRI data.

Our findings suggest that conscious perception is associated with a large-scale dynamical transition occurring within a distributed network that extends beyond sensory systems while remaining dissociable from executive and motor control circuits. In this respect, our results are broadly compatible with the global neuronal workspace framework but suggest a refinement of this account. We propose the notion of a global playground: a brain-scale network that shares critical nodes with the global workspace and supports the emergence of conscious contents, whilst being able to operate independently of the executive systems required for task performance and explicit report.

The present study introduces a general methodological framework for studying brain dynamics in response to stimulation. The modelling approach developed here provides a potential “bifurcation toolkit” that could be applied across neuroimaging modalities, sensory systems, and states of consciousness to identify latent dynamics that remain invisible to conventional analyses. By enabling the detection of signatures of conscious processing without requiring overt behavioural responses, this framework also opens unprecedented clinical perspectives, notably for assessing residual consciousness in non-communicating patients.

## Material and methods

### 1. Participants

This research received the approval of the Ethics Committee “Comité de Protection des Personnes Ouest IV”. All participants underwent a medical interview before enrolment to check for absence of MRI contraindications and to collect informed consent. They had to satisfy the following inclusion criteria: (1) signed informed consent, (2) having social security, (3) being 18 to 40 years old, right-handed, with normal or corrected-to-normal eyesight and hearing, (4) being a native French speaker and (5), having no MRI contraindication.

A total of 37 participants were enrolled between June 2022 and June 2024.

Among these, two scanning sessions were cancelled due to technical issues (one person would never hear the auditory stimuli in the scanner, another had a panic attack after entering the scanner); one participant’s data was lost at the scanning platform due to a computer dysfunction.

We therefore collected data for 34 participants (mean age was 27.53, median age 27 years old range 18 to 40 years), 21 (61.7%) were women, all were native French speakers and right-handed. Among them, 30 participants were included in the main experiment (the four remaining participants only underwent one control session). All were planned to undergo the Active and the Passive sessions on two different days. One participant could only undergo one session due to a scanner dysfunction, and two other participants had to be excluded from main analyses because their performance for the maximal intensity in the Active condition was too low (close to 75%, which is the threshold).

Among the 27 remaining participants that were included in the analyses, 14 performed the Passive session first and 13 the Active session first. Within these data, some runs were removed from the analyses due to diverse technical issues. Overall, in 22 participants we could keep all the acquired runs (9 runs in the Active and 11 runs in the Passive condition); in the other 5 participants, a maximum of 4 over 9 runs in the Active session, or a maximum of 4 over 11 runs in the Passive session had to be removed. The reasons for these runs being removed were the lack of time to finish one session scanning (participant 5 Passive), error in setting the staircase in some runs causing bad performance (participant 9 Active), or technical reasons like the scanner having to be restarted and getting desynchronized from behavioural data recording (participants 24 Passive and 25 Passive); finally, one subject experienced discomfort in the scanner, the session was therefore stopped prematurely after 6 blocks out of 11 (participant 29 Passive).

### 2. Stimulation

#### Auditory stimuli

The auditory stimuli consisted in the French vowels /a/ and /e/, pronounced by a female voice, lasting 200 milliseconds ^10^. They were synthesized using the MBROLA software with the following parameters: allophones ’a’ and ’e’ from the FR4 database (French female voice). These stimuli were embedded in a background noise, called threshold equalizing noise (TEN), which was continuous during each block. The TEN is designed to produce equal masked thresholds for all frequencies between 125 and 15000 Hz in individuals with normal hearing^85^.

The intensity of the vowel stimuli was varied relative to the intensity of the background noise to obtain five different levels, computed as Signal to Noise Ratio (SNR) in decibels (dB), ranging from minus Infinity (stimulus absent, just noise) to 0 (stimulus with the same intensity as the background noise, Figure 1B). Prior to the fMRI study, behavioural studies were conducted on 45 participants to determine the intensity levels of the “A” and “E” vowels leading to the desired points of the psychometric function for identification performance: one point below threshold, two points on the steep part of the psychometric curve with on close to threshold (75% correct), and one point clearly above threshold, close to ceiling (see supp. Figure S1). The SNR levels for the two vowels were adjusted so that the two psychometric curves overlapped. This led to SNR levels of minus Infinity, -11.5, -9.5, -7.5 and -3 for the vowel A in the Active condition (this corresponds to minus Infinity, -14, -12, -9 and -2 dB for the E vowel; for convenience, the matched SNR levels are referenced to the values for the A vowel throughout the article). These SNR levels were shifted of 2 dB in the Passive session anticipating an increase in the auditory threshold in the Passive session (minus Infinity, -10, -7.5, -5 and -1dB for vowel A ; of note, -10 was used instead of -9.5 due to a slight error but had no consequence on performance, see Figure 1C), corresponding to minus Infinity, -12.5, -10, -7 and 0 dB SNR levels for vowel E).

#### Stimulation apparatus

The auditory stimuli were played through MRI-compatible headphones (MR Confon, Magdeburg, Germany). The visual stimuli were projected on a screen and shown through an angled mirror to participants while lying into the MRI scanner. Light was always turned oO to ensure similar luminosity conditions across participants and sessions.

#### Trial structure

Stimulations were produced and presented using Matlab (R2018B) and Psychtoolbox^86^, based on the protocol previously designed by Sergent and colleagues^10^. As presented in Figure 1B, a continuous background noise (threshold equalizing noise or TEN) was played throughout the duration of each run. On each trial either no vowel was presented, or the vowel "A" or "E" was presented at one of the predefined SNR levels. Vowel identity and intensity conditions were randomly intermixed across trials within each run. Each trial started with the onset of a dot at the center of the screen that participants were instructed to fixate. The delay between trial start and stimulus onset was randomly jittered between 1 and 3 seconds. The delay between stimulus onset and the first response screen was jittered between 4 and 6 seconds. The response screens were different in the active and passive sessions (see below). These timing yielded long interstimulus intervals (including the response phases, average trial duration was 8.55 seconds for the active sessions (std 0.30), 8.96 seconds for the passive sessions (std 0.31)), a timing well adapted to the low temporal resolution of the BOLD response.

#### Staircase procedure

Before each session (active or passive), participants first underwent a staircase procedure in the scanner^87,88^ to take into account the high inter- and intra-individual variability of general auditory threshold in the scanner, due to earplugs use, headphones position, scanner noise, or individual sensitivity. This staircase aimed at determining the general volume level at which we delivered our stimuli for the rest of the session. To do so, we designed an alternative detection/identification task where participants had to click on the vowel they thought they heard (’A’ or ’E’) only if they heard one; otherwise, they were asked not to click on any vowel and let the response screen disappear. On each trial the randomly selected vowel A or E was played over the TEN at a fixed SNR level, the critical level aimed to be at threshold (always -9.5 dB for A and -12 dB for E, for both Active and Passive conditions). The staircase varied the general volume of the noise plus the stimulus. It was performed while the functional scanning sequence was running. Volume intensity was increased if participants did not detect the stimulus (hence did not click neither A or E) and decreased if they did (no matter what vowel was selected), with a progressively decreasing step. The procedure stopped when yielding a 50% detection rate. The general volume for the rest of the session was set as the average volume over the last 6 trials of the staircase.

#### Procedure and tasks for the Active and Passive sessions

During stimulus presentation, participants were asked to fixate a central point on the screen.

During the active sessions (see Figure 1B), stimulus presentation was followed by:

i. An identification task: a first response screen displaying two answer options “A” and “E” to the left and the right of the previously displayed fixation point (with the respective side of each option randomized on each block) and they had to select the one they thought they just heard. If they did not hear any letter, they were asked to choose randomly.
ii. An audibility rating: immediately after participants performed the identification task, or after a maximum reaction time of 3 milliseconds in case they did not, a second screen appeared and displayed four audibility options ranging from 1 to 4. Participants were instructed to use these options as follows:

- 1 = « I did not hear any letter, I chose randomly »
- 2 = « I think I heard something but very low »
- 3 = « I heard a letter clearly but I know it can be louder than this »
- 4 = « I heard a letter with the highest possible level ».

We collected audibility ratings using this “PAS-like” scale (“Perceptual Awareness Scale”) ^89^ with four options instead of a more graded one ranging from 0 to 10 like in the EEG study due to more drastic time constraints in fMRI as well as the necessity to use a 4-button keypad placed in participants’ right hand to answer questions, allowing them to respond quickly and minimise movements.

During the Passive sessions (see Figure 1B), each stimulus presentation was followed by either one of four possible visual tasks, with equal probability:

i. Mind-wandering probe: participants had to answer the question “What did you have in mind just now?” using four possible options: “The environment”, “The sound”, “Nothing, I am falling asleep”, or “My own thoughts”. Participants were instructed that “The sound” referred to any sound played in the headphones, and that the scanner noise was to be reported as “The environment”.
ii. General knowledge quiz: participants had to answer general knowledge questions with a varying level of difficulty by selecting one out of four possible answers and had feedback after responding or at the end of a maximum delay of 5 seconds, highlighting the right answer to keep them motivated.
iii. Visual detection task: when a green dot appeared at the centre of the screen, participants had to press any button as fast as possible; and then click again in response to a « Click to continue » message.
iv. Simple « Click to continue » message.

Active sessions included 9 blocks of 50 trials each, yielding a total of 45 trials per condition of SNR (5 levels) x Stimulus identity (2 levels: A or E) and passive sessions included 11 blocks of 40 trials, yielding a total of 44 trials per SNR x Stimulus identity condition. The total duration of each session was approximately 1h45 - 2h, depending on the duration of breaks, proposed to the participants between each block. All subjects included in the analyses underwent both sessions in a randomized order on different days (at least 24 hours apart).

It is to be noted that only one participant fell asleep during a scanning session (Active condition); participants reported a more pleasant aspect of the trivia question during the Passive sessions, while the Active session was considered more monotonous. Yet, except for two participants who had to be excluded from further analysis due to a poor performance (including the one falling asleep), all participants had very good identification performance as well as consistent audibility ratings (Figure 1C; supplementary Figure S13).

## 3. fMRI

### Data acquisition

Data were obtained using a 3-Tesla MRI scanner (Prisma fit, Siemens) at the Paris Brain Institute neuroimaging platform (CENIR). Anatomical imaging was performed using a T1-weighted sequence (voxel size 1 mm isotropic). Functional imaging was performed using multi-echo (ME-) fMRI (60 slices, field of view (FOV) 210 mm, TR 1.660 s, voxel size 2.5 mm isotropic) and several accelerating factors (multiband-3, GRAPPA-2) ^90^. ME-fMRI enables multi-TE acquisitions (various images were acquired at different echo times, TE) instead of using a single one like in single-TE EPI imaging, usually chosen as a trade-oO between BOLD weighting, overall signal-to-noise ratio, and signal loss due to magnetic susceptibility artefacts^91^.

To have enough space to use the adequate MRI-compatible headphones (MR Confon, Magdeburg, Germany), a 20-channel head and neck coil was used. Cardiac and respiratory frequencies were recorded using MR-compatible sensors during the whole acquisition for later denoising.

### Data pre-processing

The first steps of FMRI data pre-processing were performed using AFNI (https://afni.nimh.nih.gov/) for minimal pre-processing, using slice timing correction, motion correction and distortion correction.

Then, multi-echo data had to be merged into a single representative volume, which is called echo combination. The goal of echo combination is to take the various images acquired at different TEs and merge them into a single, high-quality volume that has a better signal-to-noise ratio than any single echo could provide on its own. Because T2* values vary across different brain regions, a voxel-wise T2* map was computed. These T2* values were then used to calculate weights for each echo, facilitating an optimal combination^92^. This procedure was performed with an opensource Python package, tedana (TE-Dependent ANAlysis)^91,93^.

After echo combination into a single volume, standard single-echo pre-processing was performed to normalize the volume into MNI space using SPM12 on Matlab.

Denoising was performed by regressing out 24 movement parameters (head motion artefacts modelled from shifts in brain images over time, and their derivatives) and physiology parameters^94^ using cardiac and respiratory recordings. PCA on white matter and cerebrospinal fluid masks was also used^95^. Finally, pre-processing also included optional smoothing with an 8 mm full width at half-maximum Gaussian kernel. Smoothed data were used for all univariate GLMs, while unsmoothed data were used for single-trial analyses.

### Analyses

#### I. General linear models (GLMs)

We designed three main types of GLMs for our univariate analyses:

- 1/ Model with stimulus intensity as parametric modulator (GLM1, Figures 2 & 3A, Tables 1, 2 & 3)

- first-level parametric modulation model in Active and Passive condition
- second-level contrasts for parametric modulation in Active alone, Passive alone, Active minus Passive, Passive minus Active, minus Active, minus Passive and Active / Passive conjunction
- 2/ Models with Finite impulse Response (GLM2, Figure 3B)

- first-level model with one distinct regressor per stimulus intensity
- second-level consisted in averaging beta estimates for each stimulus intensity and each timestamp across participants within predefined regions of interest
- 3/ Report-based models (GLM3)

- GLM3a: Active "Audibility" x "Stimulus intensity" (Figures 4A left & B, Tables 4 & 5)

`o` first-level model including regressors for distinct audibility rating and stimulus intensity
`o` second-level contrasts for "Not heard (audibility 1)" minus "No stimulus (stimulus intensity = SNR level 1)" and for "Heard (audibility > 1)" minus "Not heard (audibility 1)"
- GLM3b: Passive "Probed & answer Other / stimulus present" minus "Probed / stimulus absent" (Figure 4A right, Table 4)

`o` first-level model including regressors for "Probed - Sound - SNR levels 2-5", "Probed - Other - SNR levels 2-5", "Not probed" and "Probed - no stimulus (SNR level 1) with parametric regressors for SNR levels 2-5 for the three first regressors of interest
`o` second-level contrast for "Probed - Other - SNR levels 2-5" minus "Probed - no stimulus"
- GLM3c: Passive "The sound" minus "Other" (mind-wandering probes; Figure 4C, Table 6)

`o` first-level model including regressors of interest only for mind-wandering trials’ answer ("The sound" or "Other")
`o` second-level contrast for "The sound" minus "Other"
- GLM3d (additional model; in supplementary materials, Figure S12): Passive "The sound" minus "Other" and additional parametric modulation regressors based on stimulus intensity

`o` first-level model including regressors for "Probed (mind-wandering) -The sound", "Probed (mind-wandering) - Other", "Not probed", and parametric regressors on stimulus intensity for these three regressors
`o` second-level contrasts for "Probed - Sound" minus "Probed - Other" and for stimulus intensity parametric modulation on all trials (probed and not probed)

For all GLMs, first level analyses were performed on each subject individually (for a total of 27 participants), using 8mm-smoothed functional scans. We used SPM12 to estimate a GLM with a high-pass filter of 1/128 Hz (default value, so that slow signal drifts with a period longer than 128 seconds are removed) and the nuisance regressors described previously (see Data pre-processing section). The regressors of interest differed across analyses and are described below for each GLM. The regressors of non interest were included in all 1st-level GLMs and included the fixation point onset, the response screen onset (as a function of the question asked in the Passive condition because it changed the setting of the screen) and the keypress(es), all modelled as stick functions. Regression coefficients (for the regressors of interest) were estimated at the individual level. For GLM1 and GLM3, regressors for event-related analysis were obtained by convolving the timing of stimuli for our conditions of interest with a canonical hemodynamic response (HRF); and regression coefficients were taken to the group level random-effect analysis (see below). In these second-level analyses, we added covariates that were participants’ age, gender, education (either "neuroscience / cognitive science background" or other), and session order (either "Active then Passive" or "Passive then Active").

##### 1. Parametric modulation analysis (GLM1)

The first model (GLM1) included two boxcar functions of 200 milliseconds (one for each vowel type, A or E) modelling the signal at the vowel stimulus onset, which was parametrically modulated by the stimulus intensity. The regressor was convolved with a canonical hemodynamic response function (HRF). The aim of parametric modulation is to better model the effect of stimulus intensity on BOLD signal and unravel what regions of the brain show a significant modulation of their activity by auditory stimulus intensity^96^. The stimulus intensity values of the parametric modulators matched the progression of SNR levels (same interval between level 2 and 3 and between 3 and 4, and double interval between 4 and 5) and were mean-centred, yielding the values [-2.2; -1.2; -0.2; 0.8; 2.8] for the five respective levels of stimulus intensity, for both vowels and in both the Active and Passive sessions.

For second-level analyses, we constructed a flexible ANOVA design to facilitate both comparative and conjunction analyses. This allowed us to test for the effect of Active and Passive alone, respectively (Figures 2A & B, S2-3, Table 1), as well as minus Active and minus Passive (results in supplementary Figure S6 and Table S1). We then performed a conjunction analysis across the Active and Passive conditions, tested against the conjunction null hypothesis. Unlike the "global null", which could be significant if only one condition showed a strong effect, the conjunction null is a more rigorous "logical AND" test. It requires each constituent contrast to be independently significant, thereby identifying the regions significantly activated in both conditions (Friston et al. 1999; 2005; Nichols et al. 2005). We also performed Active minus Passive and Passive minus Active contrasts analyses. Results of these analyses are available in Figures 2 & 3A, Tables 1, 2 & 3 and supplementary Figures S4-5 & S7.

#### Statistical thresholding, multiple comparison correction, figure rendering, anatomical labelling

Correction for multiple-comparisons was performed using SPM-implemented Family Wise Error (FEW) correction at a p-value threshold of 0.05 whenever possible. For contrasts where the statistical power was weaker due to the small subset of trials used, we also considered activations at an uncorrected p-value threshold of 0.001; these are therefore only exploratory analyses. Regions and subregions that appeared activated in the different analyses were identified and labelled manually with the help of the Bioimage Suite 96, the Brainnetome atlas 97 and the FSLeyes and MRIcroGL implemented atlases (Harvard-Oxford and AAL atlases). The anatomical templates used for figures are MNI152 from MRIcro GL for T1 slices, and brainMESH ICBM152 from Surfice for 3D surfaces.

##### 2. Finite Impulse response (FIR) analysis (GLM2)

This GLM was used to model timeseries directly for each stimulus intensity; it therefore included five regressors of interest, one for each intensity. We used a FIR approach rather than assuming a canonical HRF, thus allowing a description of the temporal dynamic of the BOLD signal. The FIR GLM (GLM2) estimated the BOLD response at seven time points per trial, from -1 TR (so -1.66 seconds) before to +6 TR after stimulus onset. From this model, we extracted the regression estimates (beta values) for each time point across all trials for a given intensity level, for each subject. These beta estimates were then z-scored across SNR levels for each subject and then averaged across subjects, giving one value per voxel of the full 3D volume. Finally, we extracted these beta estimates from five predefined regions of interest that were extracted from the 2nd level FWE-corrected Active and Passive conjunction parametric modulation analysis (GLM1, Figure 3A), namely, left and right superior temporal gyri, supplementary motor area / middle cingulate cortex, and left and right prefrontal / anterior insular cortex. The main advantage of a FIR analysis is that it is not constrained by any predetermined hemodynamic response function (HRF); it therefore allowed us to compare not only differences of "amplitude" of BOLD signal (beta estimate values) across ROIs, but also differences of delays to reach the peak^97^. Results of these analyses are available in Figure 3B.

##### 3. Models including perceptual report and mind-wandering responses (GLM3)

The second type of GLMs aimed at investigating activations underlying subjective reports (Heard and Not heard trials in the Active sessions, and mind-wandering reports in the Passive sessions). For these models, regression coefficients were estimated at the individual level and then taken to group-level random-effect analysis using a one-sample t-test. In these analyses one major challenge was to avoid stimulus intensity-related confusion, since there is a correlation between the probability to report a stimulus as heard and its intensity.

#### (1) Active condition audibility-based analysis (GLM3a)

In the Active condition, the probability of reporting a stimulus as "Heard" and the stimulus intensity were not orthogonal. Thus, GLM3a included 40 boxcar functions of 200 milliseconds duration, modelling the signal at stimulus onset: these regressors modelled trials across all stimulus intensities (hence, 5 levels) and all audibility ratings (hence, 4 levels) for each vowel A or E (2 levels), for a total of 5x4x2 = 40.

We constructed two contrasts of interest:

- Stimulus not heard versus No stimulus: comparing trials with stimulus present but reported as not heard (intensity level > 1, audibility rating of 1) against trials with no stimulus (intensity level 1, irrespective of audibility rating).
- Stimulus heard versus Stimulus not heard at threshold: comparing trials reported as heard (threshold intensity level, audibility rating of 2, 3 or 4) against trials not heard (threshold intensity level, audibility rating of 1).

Importantly, to orthogonalize stimulus intensity and report, we limited this second contrast to the trials considered at threshold, i.e. either intensity levels 3 or 4 based on the behavioural results of each participant (to do so, we manually inspected individual intensity levels at which variability peak occurred and 75% identification was reached; see supplementary Figure S13). Hence heard versus not heard trials were contrasted for identical external stimulation. This analysis therefore only involved 1/5th of all trials in the Active sessions. Results of these analyses are available in Figures 4A & B, Tables 4 & 5 and supplementary materials S8 & 10.

#### (2) Passive condition mind-wandering - based analysis (GLM3b, c and d)

To try and probe participants’ conscious and unconscious perception during the Passive sessions, we used the subset of trials containing mind-wandering probes, considering that the answer "The sound" was a proxy for "Heard", while any other answer ("Other") was one for "Not heard". This analysis was therefore performed on only 1/4^th^ of all trials from the Passive sessions. We could not select trials at threshold intensity due to the absence of behavioural marker and the further lack of power that would be caused by selecting only 1/5^th^ out of the 1/4^th^ of trials with a mind-wandering probe. Of note, the following GLMs no longer distinguished between vowels A and E to simplify the models.

GLM3b was built to probe unconscious processing in the Passive condition and therefore included four boxcar functions of 200 milliseconds duration, modelling stimulus onset as being either (1) probed and heard ("Sound"), (2) probed and not heard ("Other"), (3) not probed, or (4) probed but absent (SNR level 1). Additionally, the three first regressors were parametrically modulated by the stimulus intensity, similarly to GLM1 except it only involved SNR levels 2 to 5, while SNR level 1 was individualized as "Stimulus absent". This design was made to (1) account for stimulus intensity with the parametric modulation regressors and (2) only subtract trials within the mind-wandering probed ones [distinguishing stimuli absent probed from unprobed trials to prevent unbalanced contrast, which would have been the case if subtracting regressors for Probed not heard and No stimulus (probed and not probed)]. The contrast of interest compared "Probed; answer Other" trials and "Probed; no stimulus" trials. Results of these analyses are available in Figure 4A, Table 4 and supplementary Figure S9.

To probe conscious processing in the Passive condition, we conducted a first analysis considering trials based only on mind-wandering probes, irrespective of their stimulus intensity. Although not accounting for differential stimulus physical intensity effect, this first approach had the advantage of being the simplest and therefore the most well-established method for an initial assessment of conscious perception in the Passive condition across a small number of trials. GLM3c therefore included four boxcar functions of 200 milliseconds duration: modelling the signal at vowel stimulus onset (vowel A or E, hence 2 levels) for trials containing mind-wandering probes, stratified by participants’ answer, "The sound" or "Other" (2 levels), hence 2x2 = 4 in total. The contrast of interest compared "The sound" with "Other" trials. Results of these analyses are available in Figure 4C, Table 6 and supplementary Figure S11.

To go further, we built a secondary, exploratory design that not only probed "Sound" versus "Other" answers, but also stimulus intensity, including unprobed trials. GLM3d was therefore performed as an exploratory analysis and was similar to GLM3c except that it also included unprobed trials and considered stimulus intensity using parametric modulation. It therefore contained three boxcar functions of 200 milliseconds durations modelling stimulus onset depending on being probed and heard, probed and not heard, or not probed (both vowels considered together to decrease the number of regressors overall); additionally, each of these three regressors was parametrically modulated by the stimulus intensity, identically to GLM1.

We then created two contrasts:

- "Probed - Sound" versus "Probed - Other" trials to investigate "pure" conscious perception neural correlates
- stimulus intensity (based on the three parametric modulators), to investigate the mere stimulus intensity effect.

Results of these analyses are available in supplementary materials S12.

Details of statistical thresholding, multiple comparison correction, figure rendering, and anatomical labelling were identical to those described for GLM1 (section I.1).

### II. Single-trial analysis

#### 1. Single-trial general linear model (GLM)

To investigate single-trial activity in an fMRI dataset, we needed to specify a dedicated GLM that considered each trial as a unique regressor instead of grouping all trials of a given condition as one regressor. Such an approach carries the risk of losing statistical power due to noisier results. We therefore used the GLMsingle toolbox^69^ that enables building a single-trial design matrix and improving signal-to-noise ratio thanks to HRF selection, a specific cross-validation denoising ("GLMdenoise"), as well as fractional ridge regression. The single-trial GLM was fitted to the unsmoothed data and the single-trial beta estimates were obtained for each subject and each voxel (for a total of 69x83x68 = 389.436 voxels per volume), resulting in a tremendous amount of data (389.436 voxels x 440 or 450 trials for 2 sessions for each of the 27 subjects, total = 9.3581.10^9^ values).

Next step was therefore to reduce dimensionality before investigating distributions and inter-trial variability of the single-trial values as a function of stimulus intensity, as was done in Sergent and colleagues’ 2021 paper with EEG data. To do so we designed a multivariate approach (see schematic pipeline in supplementary Figure S15), consisting in feeding single-trial beta-estimates to a linear decoder performing cross-classification for stimulus absence versus presence, across the five different levels of intensity and extract a decision value for each trial, hence having a single value for each region of interest, trial, condition and subject. This analysis is detailed below (section II.3).

#### 2. Defining regions of interest (ROIs) for single-trial analysis

The need for custom designed ROIs for analysing single-trial dynamics was motivated by two lines of constraints : (i) we wanted to analyse how activation dynamics might change across different regions of the brain, but had to reduce the dimensionality of the data; and (ii) we could not directly use ROIs derived from our group-level analyses without risking to contaminate each subject’s single-trial data with too many non-informative voxels, or conversely losing information when excluding some informative voxels of one given subject. This would have been particularly impactful knowing that single-trial beta estimates are much noisier than standard beta estimates. Hence, we used individualized 1st-level univariate results from the parametric modulation analysis described previously (GLM1; uncorrected statistical parametric maps, p < 0.001) to ensure using all voxels responding to the stimuli for each participant individually.

Since we predicted that bifurcation dynamics should be found in regions involved in conscious access and not in regions merely involved in primary sensory processing, we distinguished, among all responding voxels of each participant, two regions of interest: "Primary Auditory Region", i.e. the individual activation cluster voxels located in Heschl’s gyri and "beyond Primary Auditory Region", i.e. all remaining voxels. Anatomical ROIs corresponding to the primary auditory regions were defined using the Harvard–Oxford probabilistic cortical atlas distributed with FSLeyes (FMRIB Software Library (FSL) image viewer ^98^). Bilateral Heschl’s gyrus probability maps were used as an anatomical proxy for primary auditory cortex. Probabilistic atlas maps were resliced to each subject’s functional space (2.5 mm isotropic voxels) using trilinear interpolation in SPM12 to ensure voxel-wise correspondence with subject-specific activation maps. The resliced probabilistic maps were then thresholded at 10% or more probability and binarized to generate anatomically constrained ROI masks. Subject-specific activation clusters were then intersected with the binarized Heschl’s gyrus masks to obtain ROIs corresponding to cluster voxels located within primary auditory regions. Complementary ROIs, excluding primary auditory cortex, were generated by subtracting the Heschl’s gyrus mask from the activation clusters (beyond primary auditory regions). All ROIs were defined in individual functional space and used for subsequent single-trial analyses. Of note, to avoid inclusion of low-confidence voxels a probability threshold of 10% was chosen to balance anatomical specificity and inter-individual variability in Heschl’s gyrus morphology. Indeed, the Harvard–Oxford probabilistic atlas provides voxel-wise probabilities (0–100%) reflecting the proportion of subjects in whom each voxel belongs to a given anatomical label. Still, we must note that three participants were missing identifiable activity in Heschl’s gyri as defined in the atlas in some sessions. Subsequent ROI-based analyses were therefore conducted without those participants’ concerned sessions (participant 9, Passive session; participant 29, Passive session and participant 22, both sessions removed - Active session showed no primary auditory region cluster and therefore Passive session data were not used in the absence of an Active benchmark for that subject).

After overlapping individual univariate clusters with atlas-defined Heschl’s gyri we obtained ROIs containing from 18 to 403 voxels (mean 153, std 97) for Active condition primary auditory regions, 15 to 570 voxels (mean 191, std 131) for Passive condition primary auditory regions, 3890 to 18283 voxels (mean 9647, std 3825) for Active condition beyond primary auditory regions and 1184 to 7327 voxels (mean 2851, std 1248) for Passive condition beyond primary auditory regions.

#### 3. Extracting single-trial neural values using multivariate pattern analysis (supplementary Figure S15)

We fed a linear multivariate pattern classifier (Support Vector Machine; SVM) with the previously computed single-trial beta estimates from all the voxels of each ROI (see paragraphs 1&2 above). The classifier was trained to discriminate trials containing maximum-intensity stimulus from trials with no stimulus and then tested across all five stimulus intensity levels, in a similar way as in Sergent et al 2021^10^. Classification was implemented using a leave-one-run-out cross-validation scheme with the Decoding Toolbox, a MATLAB toolbox designed for multivariate pattern analysis and cross-classification. This toolbox also enabled the application of recursive feature elimination (RFE) to select voxels forming the most informative pattern for classifying maximal intensity versus absent stimuli^99–101^. From the cross-classification outcomes, we then extracted the decoding performance for each subject, condition (Active vs. Passive), and ROI. Importantly, we also customized the toolbox functions to extract trial-wise decision values, defined as the distance of each trial’s activation pattern to the decision hyperplane^10^. For clarity, we refer to these single-trial decision values as "neural values" in the rest of the article. Neural values were then used as a one-dimension proxy of each participant’s multivariate activation pattern for a given ROI, computed separately for each stimulus intensity. We z-scored these decision values within each participant so that they could be compared and averaged across participants for group-level analysis. See supplementary Figure S14 for a summary of selected voxels after applying RFE.

#### 4. Modelling distributions of single-trial neural values

We investigated single-trial dynamics using single-trial neural values for each participant, each stimulus intensity, each condition and each ROI (cluster restricted to primary auditory regions / cluster beyond primary auditory regions), obtained through the processes described in section II.3.

We first visually assessed the distribution, averaged values as a function of stimulus intensity and inter-trial variability profiles both individually and at group level (Figures 5B & C) for sanity check and subjective assessment of unimodal versus bifurcation aspects and inter-trial variability peak at threshold.

Subsequently, we searched for ways to mathematically quantify, if existing, the differences between conditions and ROIs of inter-trial variability profiles and unimodal versus bifurcation dynamics. All subsequently described analyses were performed using Python and open-source toolboxes^102–107^. The optimization scripts and visualization parameters were refined through iterative consultation with Gemini 3 Flash^108^.

A first, coarse approach, to distinguish between unimodal and bifurcation dynamics is to examine the profile of inter-trial variability (standard deviation across trials) as a function of stimulus intensity and use the variability peak ratio as an index of bifurcation dynamics. Then a more thorough approach consists in fitting the two models to the observed distribution of responses across trials and perform a model comparison yielding a “bifurcation score”. Both approaches are described below.

##### 1) Variability peak ratio (VPR, Figure 6A)

This ratio provides a measure of increased inter-trial variability at intermediate stimulus intensity stimuli (the three middle SNR levels) compared to extreme intensity stimuli (lowest, namely no stimulus (noise, SNR level 1), or maximal intensity (SNR level 5)); the higher the VPR, the higher the relative variability at threshold.

It can be formalized equivalently by the formula that follows:

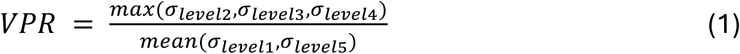

Where σ_*level i*_ are the inter-trial variability computed for intensity level *i*.

To assess whether the variability peak at threshold differed between sensory and higher-order areas, we performed planned comparisons between ROIs (primary auditory regions versus beyond primary auditory regions) separately for each task condition (Passive and Active): for each condition, we compared the distribution of VPR scores across participants using an independent samples t-test (or paired t-test, depending on participant matching) between the primary auditory cortex (primary auditory regions) and the non-primary auditory area (beyond primary auditory regions). Statistical significance was determined using a standard alpha level of p < 0.05. To account for multiple comparisons across the two task conditions, p-values were adjusted using the Bonferroni-Holm correction.

##### 2) Model comparison

###### a) Models

We modelled single-trial neural activations (*y*) across the five levels of stimulus intensity using two competing models, a unimodal one and a bifurcation one. Both models were anchored to the participants’ individual extreme intensity levels’ statistics (mean and standard deviation of minimal and maximal intensity levels, respectively *μ*min, *μ*max, σmin, and σmax). We also defined a null model to compute each of the competing model’s gain compared to noise. The number of optimized parameters was different across models (see section below), which did not raise validity issues as long as a cross-validation design was respected for model comparison to prevent overfitting (see section *Leave-One-Run-Out Cross-Validation*). Below the definition of each model is detailed.

***Null Model*** *(for comparison).* This model assumes that the activity does not change with stimulus intensity, and hence that the probability *P* of a neural activity value of *y* for a given trial at each stimulus intensity can be described by a single normal distribution whose parameters are those computed across all trials:

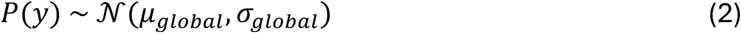

Where *μ_global_* is the averaged value across all intensity levels trials, and σ*_global_* is the standard deviation across all trials as well. These two parameters are directly measured on training data and do not need any gradient descent to be fitted.

***Unimodal Model.*** This model assumes that the probability *P* of a neural value of *y* for a given trial follows a normal distribution whose mean shifts progressively with stimulus intensity. The distribution at each stimulus intensity is defined as:

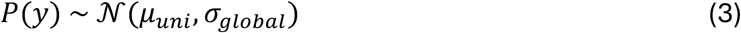

Where:

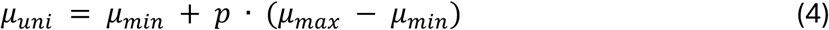

And σ*_global_* is the mean of the two extreme standard deviations, σmin and σmax.

- Parameters σ*_global_*, *μ_min_*_)_and *μ_max_* are therefore directly measured on training data.
- The parameter *p* in [0, 1] represents the "mean position" and is optimized via gradient descent during the cross-validation fitting process for each stimulus intensity simultaneously.

So this model had a total of eight estimated parameters, three directly estimated from the training data, and five optimized via gradient descent.

a. ***2. Bifurcation Model***. This model assumes that, around threshold, the brain switches between two possible equilibria corresponding to conscious or non-conscious processing of the same stimulus, and yielding Gaussian Mixtures defined as follows:

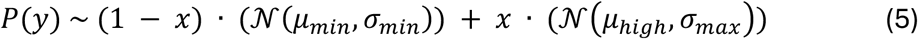

Where:

- The parameters *μ_min_*, σ*_min_* and σ*_max_* are measured from training cross-validation data but not optimized.
- Ten parameters are optimized via gradient descent during the cross-validation fitting process (two for each stimulus intensity):
- `o` *x* in [0, 1] represents the probability of being in the “high” state
- `o` *μ_high_* in [*μ_min_*, *μ_max_*+ 2] is the mean activity of the “high” state

So a total of thirteen parameters were estimated for this model, three directly from the training data, and ten via gradient descent optimization.

##### b) Constrained model fitting

###### 1/ Monotonic constraint

A critical requirement for biological plausibility in these models is the assumption of monotonicity: as the physical signal strength increases, the latent parameters representing (i) for the unimodal model, the mean position (*p*) and (ii) for the bifurcation model, the probability of "high” state (*x*) and the expected mean of the "high” state (*μ*_21-2_) should also increase. To enforce this, we utilized Sequential Least Squares Programming (SLSQP) for parameter estimation^109^. Unlike standard unconstrained solving algorithms such as L-BFGS-B (limited Broyden Fletcher Goldfarb Shanno) that can only take into account box constraints (i.e., minimizing a cost function within predetermined bounds)^110,111^, SLSQP allowed the implementation of linear inequality constraints such that:

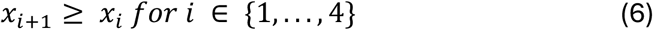

Where *x*_1_ represents the model parameter (*p* for the unimodal model, *x* and *μ_high_* for the bifurcation model) optimized for the *i*^th^ stimulus intensity level. This constraint prevents the models from overfitting to the local noise that would otherwise result in non-monotonic, hence physiologically aberrant, psychometric functions.

###### *2/* *VPR-based penalty*

Additionally, given that the variability peak ratio (VPR) emerged as a highly discriminant metric for our neural data, we incorporated a VPR-based penalty into the model fitting procedure. This constraint was designed to penalize the bifurcation model if it failed to capture this characteristic variability peak observed at perception threshold. While this penalty was applied uniformly to both Unimodal and Bifurcation models to ensure a balanced comparison, it exerted no practical influence on the unimodal fit, given that the standard deviation parameter in that model is fixed across all stimulus levels.

The VPR-based penalty (*Pen_VPR_*) was computed as the absolute difference between predicted VPR, computed with the model’s fitted parameters and the empirical VPR, computed on training data within each cross-validation fold like the rest of the fixed parameters, weighted by the parameter λ:

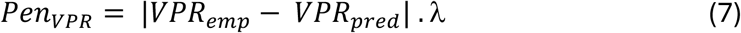

Where:

- ë was set at 10
- *VPR_emp_* was computed with formula (1), based on training data’s standard deviation values for each stimulus intensity level
- *VPR_emp_* was computed with formula (1), based on each stimulus intensity level’s calculated variance from the fitted parameters *μ_high_* and *x* as described in formula (6) and according to the total mixture variance defined by:

`o` *μ_pred_* = (1 − *x*) . *μ_min_* + *x* . *μ_high_*; which is predicted mean of a given stimulus intensity level after fitting all parameters
`o` 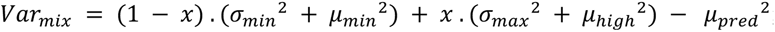; which gives the variance of a gaussian mixture as defined by formula (6)

The VPR-based penalty *Pen_VPR_* is then subtracted from the computed log-likelihood of the model, computed for each cross-validation fold, as described below.

#### c) Optimization and model comparison

The cost function we minimized was the negative log-likelihood of the data given the model, supplemented by the VPR penalty term (λ = 10) to further regularize the fits toward empirical individual variability patterns and by the monotonic constraint, forcing both models to predict higher neural activity given a higher stimulus intensity. To ensure model generalizability and prevent overfitting, particularly for the more flexible Bifurcation model which has more parameters, we employed a Leave-One-Run-Out cross-validation scheme. For each participant, the data were partitioned by experimental blocks. In each fold, the models were trained on *N-1* blocks to estimate (1) fixed fitted parameters (*μ*min, *μ*max, σmin, and σmax) and (2) optimized parameters (Unimodal model’s *p* and Bifurcation model’s *x*, *μ*_21-2_) by maximizing the log-likelihood (LLH; or equivalently minimizing the negative LLH) of the training subset. The predictive power of these parameters was then evaluated by calculating the LLH of the held-out test run. Of note, the VPR-based penalty was only applied for LLH maximization and not for evaluation LLH, since its goal was to help fitting the parameters during model training. This process was repeated for all *N* runs, and the resulting test log-likelihoods were averaged across folds to provide measure of model fit that is immune to overfitting and naturally corrects for the different number of fitted parameters in our three models.

#### d) The Bifurcation Score (*S_bif_*, Figure 6B)

The comparison between models was quantified using a weighted Bifurcation Score. This metric combines the raw difference in predictive power between the Bifurcation (*LLH_bi_*) and Unimodal (*LLH_uni_*) models with a scaling factor representing the overall quality of fit relative to the Null baseline (*LLH_null_*).

First, we computed the model delta (Δ*L_comp_*) and the mean improvement over null (Δ̅):

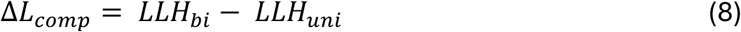

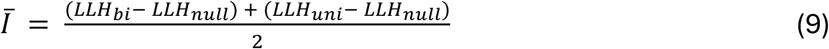

And the bifurcation score was then defined as:

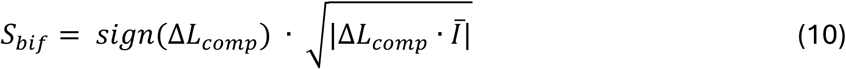

This transformation ensured that the score remained in the units of log-likelihood while penalizing datasets with low signal-to-noise ratios, by pondering the log-likelihood difference between Unimodal and Bifurcation models with their gain against the null model.

### e) Statistical Analysis

All statistical analyses were performed in Python using the scipy.stats, pandas, and numpy libraries. Due to the non-parametric nature of the neural metrics, statistical significance was assessed using one-tailed Wilcoxon signed-rank tests. The use of one-tailed tests was justified by our specific, pre-defined directional hypotheses. We indeed predicted that (1) the primary auditory regions would exhibit a unimodal profile (Bifurcation score < 0), while regions beyond the primary auditory regions would show a bifurcated profile (Bifurcation score > 0) and (2) the bifurcation strength would be significantly greater in beyond primary auditory regions compared to primary auditory regions, and significantly enhanced in the Active condition compared to the Passive condition. Statistical significance was defined at an alpha level of alpha = 0.05. Because our primary interest lay in the directionality of these neural effects, only one-sided p-values are reported. All paired comparisons (Active vs. Passive and NoA1 vs. A1) were performed only on subjects with data available for both conditions to ensure internal consistency.

## 3) Single-trial prediction of report based on the Bifurcation model

### a) Bayesian approach

To investigate the relationship between extra primary auditory clusters neural values and subjective perception report we used a Bayesian approach. For each participant and task (Active and Passive) we modelled the distribution of single-trial neural value (*y*) using the bifurcation Gaussian mixture model previously described. We used the fitted parameters (this time without cross-validation, since the aim was not to assess the model’s accuracy) to decode the behavioural report on a trial-by-trial basis. As previously defined:

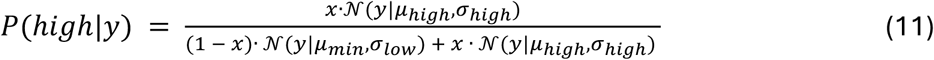

where *y* represents the neural value for a given trial, and *x* is the fitted mixing proportion representing the prior probability of a "high” state (corresponding to predicted heard trials).

To determine the likelihood of "Heard" for a given trial, we calculated the posterior probability that the observed neural value *y* originated from the "high” distribution. This was derived using the following formula:

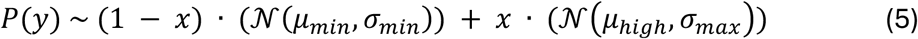

The posterior probability *P*(*high|y*) gives values between 0 and 1, with values approaching 1 indicating a "high" state, hence a "Heard" trial, and values near 0 corresponding to a "low" state, hence "Not heard" trial.

### b) Prediction evaluation using AUC (Figures 6C & D)

Rather than using raw accuracy percentages, which can be biased by class imbalances at extreme stimulus intensities, we evaluated prediction performance using the Area Under the Receiver Operating Characteristic Curve (AUC-ROC). The AUC provides a threshold-independent measure of how well the bifurcation model applied to neural values discriminates between the "Heard" and "Not heard" behavioural categories. We further validated this approach by examining the distributions of the calculated posterior probabilities. As previously described for other analyses, "Heard" / "Not heard" reports were considered either audibility > 1 versus audibility = 1 or mind-wandering answer "The sound" versus other answers for the Active and Passive conditions, respectively.

### c) Participant exclusion

Because there was no guarantee that mind-wandering probes were a reliable measure of "Heard" reports in the Passive condition, although as shown in Figure 1C, at group level it was, we screened individual mind-wandering responses to identify potential outliers, i.e. participants whose proportion of answer "The sound" to mind-wandering probes did not correlate positively with stimulus intensity. We computed the Spearman correlation coefficient ρ between the physical stimulus intensity level and the proportion of "The sound" answers for each subject (supplementary Figure S16). We excluded four participants from prediction evaluation, who exhibited negative correlation (ρ < 0), suggesting that their mind-wandering probes were not reflecting their stimulus perception, hence not suitable as a true label for prediction performance measure.

## 4. Eye-tracking and pupillometry analyses

### Data acquisition

Eye-tracking was performed during all sessions using an MRI-compatible device, Eyelink 1000. Calibration was performed at the beginning of each experimental session asking the participants to fixate successively nine points distributed across the screen, then gaze coordinates and pupil area were recorded at a sample rate of 1000 Hz. The first objective was to verify correct fixation, with feedback sent to the experimenter at the end of each block, giving the mean performance of fixation during all the trials of the block. The experimenter could therefore give feedback to the participant at the end of a given block if an improvement of fixation was required. A secondary objective was to study the pupil diameter as a function of the stimuli, as it has been described to be related to perception of not only visual^30^ but also auditory stimuli^79,81^, in order to confront electrophysiological signal with fMRI BOLD signal. To do so, we used Matlab and Fieldtrip functions^112,113^ to preprocess the pupil area time-series and plot them time-locked to the stimuli as a function of perception or SNR level in both active and passive conditions.

Due to technical difficulties during recording (loss of eye-tracking while scanning or necessity to restart the stimulation protocol during a scanning session), we had to deal with an important loss of data. Consequently, several subjects’ data were removed from analysis for one condition or another, when not enough recorded trials were available to compute pupil area time-locked to the stimulus (four subjects from the active group, and two subjects from the passive group).

### Pre-processing and event-related analysis

Raw data were segmented into trials time-locked to stimulus onset (-1 s to +5 s) and resampled at 100 Hz using FieldTrip. Preprocessing followed a fixed order to ensure signal integrity: first, blinks were detected (using both Eyelink-defined blinks and additional automated detection) and linearly interpolated. Second, we regressed out the residuals of blink- and saccade-linked pupil responses and Z-scored the continuous signal per participant to account for inter-individual variability in pupil size and reactivity. Subsequent analyses were performed on the EyePupil "channel" (recorded pupil diameter only, not eye gaze position coordinates).

To remove slow instrumental and physiological drift without contaminating the stimulus-evoked response, we applied a linear "bridge" detrending procedure on the epoched data. For each trial, a linear ramp was fitted between the mean pupil value in the pre-stimulus window (-0.5 to 0 s) and the trial end-window (4.5 to 5 s) and subtracted from the entire trial. Following detrending, trials were baseline-corrected by re-zeroing the signal to the pre-stimulus window (-0.5 to 0 s). To further clean auditory stimuli-driven signal, we performed a subject-level baseline correction. For each participant, the mean pupil trace of the noise-only condition (intensity level 1) was subtracted from the traces of all other stimulus intensity levels.

After single-trial cleaning and normalization, trials were averaged within each participant according to experimental condition: (i) stimulus intensity (Figure 3C): trials were grouped by intensity level and averaged per participant, and (ii) behavioural report (Figure 4D): trials were grouped as Heard versus Not heard. In the Active condition, this split was based on trial-by-trial audibility ratings at the participant’s specific threshold intensity level; in the Passive condition, it was based on responses to mind-wandering probes (grouped as "Sound" versus "Other") for all trials where a stimulus was present (stimulus intensity levels 2 to 5).

Group-level time courses were obtained by averaging participant-level waveforms across subjects. Variability across participants was quantified as the standard error of the mean (SEM) at each time point. For visualization only, group mean time courses were lightly smoothed using a moving average (250 milliseconds window). Statistical significance was assessed using point-by-point paired t-tests (e.g., testing if traces significantly differed from zero, or comparing Heard vs. Not heard trials). Significance clusters were defined as contiguous time points where p < 0.05.

## Acknowledgements

We thank Jean Lorenceau for helping with the pupillometry analysis and interpretation. We thank Paolo Bartolomeo for helpful discussions on anatomical interpretations, Jacob Prince for helping with the GLMsingle toolbox, Kai Görgen, Martin Hebart & Carl Allefeld for helping with the Decoding Toolbox, and Martina Corazzol for schematic figures of experimental design.

This work was supported by the Fondation pour la Recherche Médicale (FRM), grant number « FDM202306017112 », to Julie Boyer, by an ERC grant to Claire Sergent: 101044362 (CONSCIOUSBRAIN), and by a grant from the Agence Nationale de la Recherche to Claire Sergent: ANR-17-CE37-0004-01 (FlexConscious).

All scripts used for experiments and analyses can be found on GitHub: https://github.com/JulieB1234/soundfmri_public.

