## Supplementary Figures & Tables for "Bifurcation dynamics in a shared network beyond sensory areas characterize conscious auditory perception independently of report"

**SUPPLEMENTARY INFORMATION**

**Authors:** J Boyer[1], N Beraud[1], B Beranger[2], T Hardy[1], B Türker[1], H Gouyette[1], A Lopez-Persem[3], C Sergent[1]

*1. Université Paris Cité, INCC UMR 8002, CNRS, F-75006 Paris, France*

*2. Centre de NeuroImagerie de Recherche - CENIR, Institut du Cerveau - ICM, Paris, France*

*3. FrontLab, Sorbonne University, Institut du Cerveau - Paris Brain Institute - ICM, Inserm, CNRS, AP-HP, Hôpital de la Pitié Salpêtrière, Paris, France*

**Table of contents**

Figure S1: Behavioural pilots psychometric curve

Figure S2-5: Parametric modulation full fMRI maps: Active, Passive and Subtractive contrasts

Figure S6: Parametric modulation full fMRI maps: Active and Passive deactivations

Table S1: Table of results for Figure S6

Figure S7: Parametric modulation full fMRI maps: Active and Passive conjunction

Figure S8-9: Full fMRI maps: unconscious processing, Active and Passive

Figure S10-11: Full fMRI maps: conscious processing, Active and Passive

Figure S12: Alternative model for Passive conscious processing (GLM3c)

Figure S13: Individual behavioural results

Figure S14: Voxels selected by RFE (recursive feature elimination) during single-trial multivariate analysis

Figure S15: Single-trial multivariate processing pipeline

Figure S16: Correlation between mind-wandering answer "The sound" and stimulus intensity

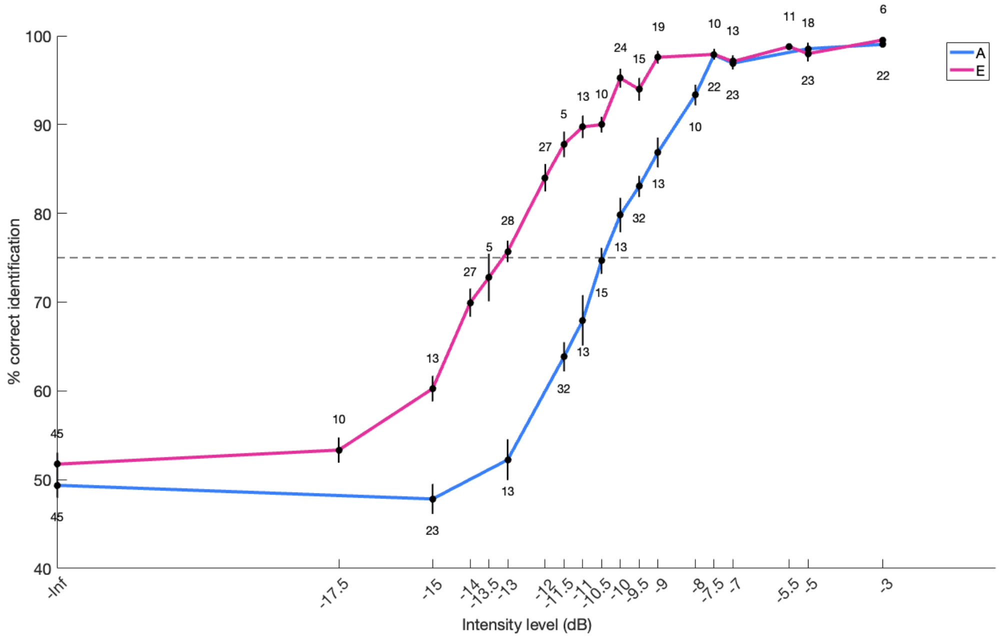


**Figure S1. Behavioural Pilots Performance Results.** This figure displays identification performance (in % correct) across all of the 45 pilot participants tested across 18 different levels of intensity separately for both A and E vowels. Threshold level is represented by a dotted line at 75% correct identification. The plot illustrates the shift across both psychometric curves (chance level reached for a weaker intensity level for vowel E than A) as well as a slight difference of slope (steeper for vowel A), justifying the use of different SNR levels for A and E vowels in our main experiment.

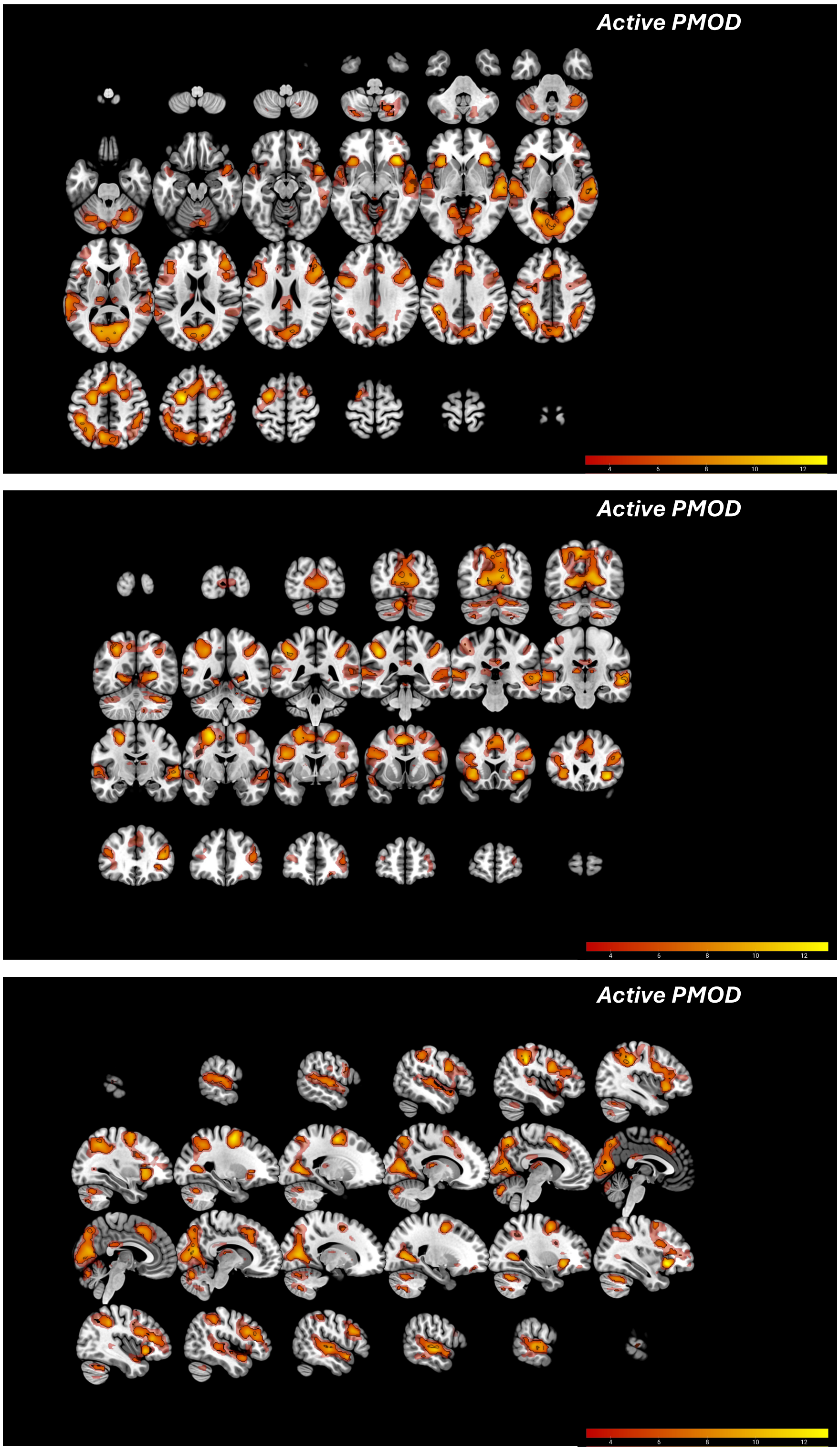


**Figure S2.** Full activation maps (top panel: axial slices from left to right; middle panel: coronal slices from anterior to posterior; bottom panel: sagittal slices from left to right) for the Active condition parametric modulation analysis (PMOD). Details for correction and display are the same as principal Figure 2A.

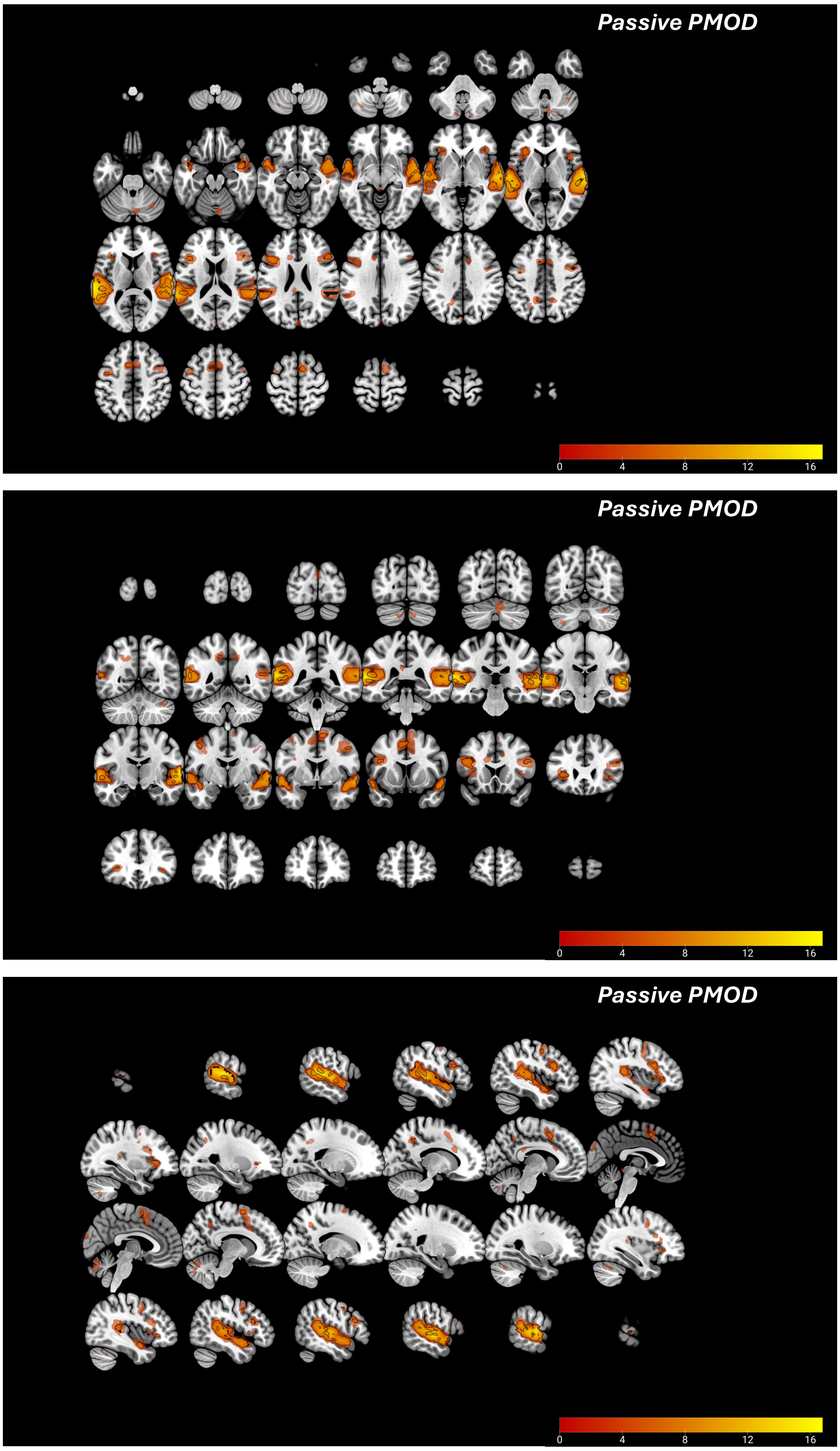


**Figure S3.** Full activation maps (top panel: axial slices from left to right; middle panel: coronal slices from anterior to posterior; bottom panel: sagittal slices from left to right) for the Passive condition parametric modulation analysis (PMOD). Details for correction and display are the same as principal Figure 2B.

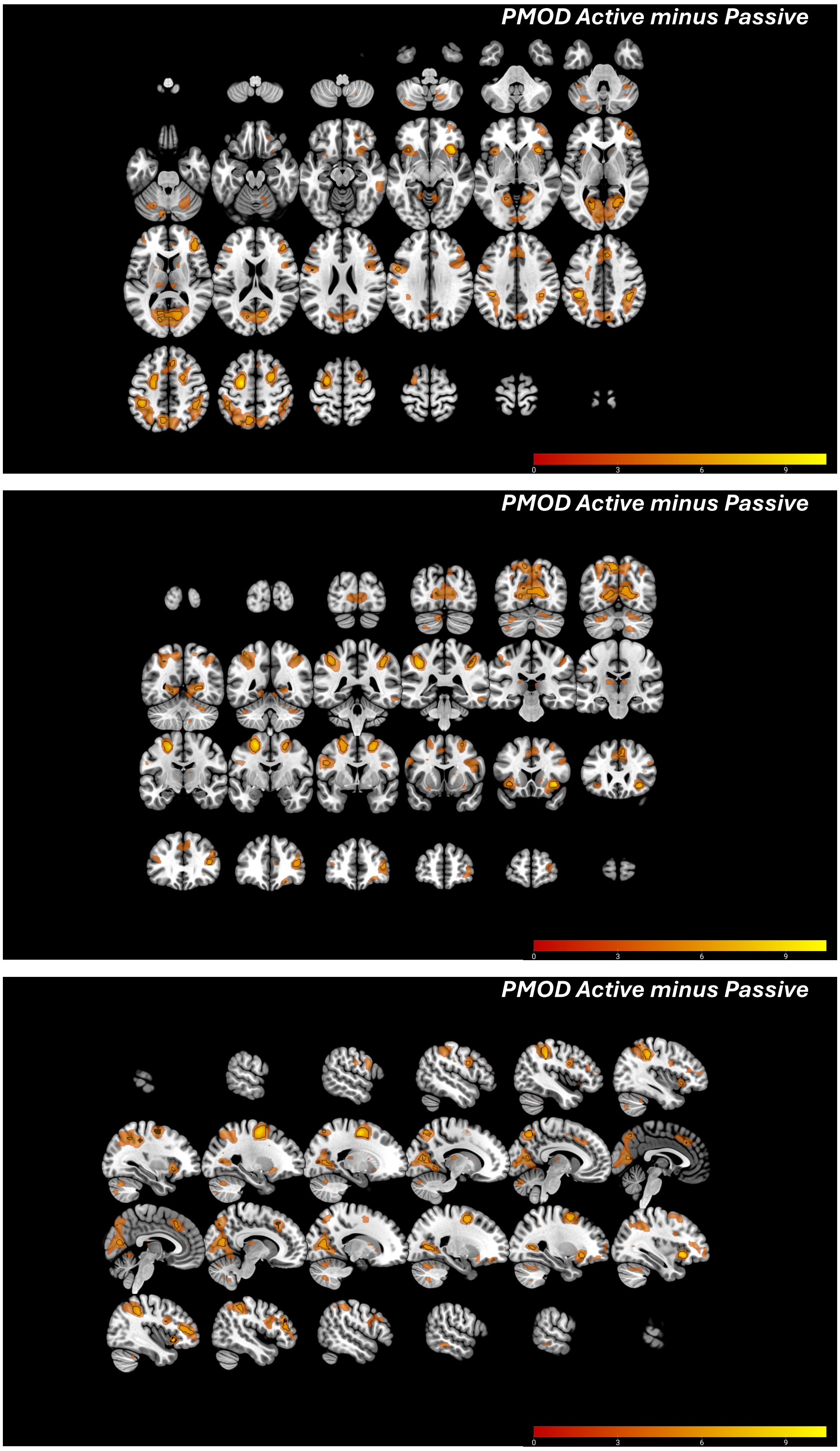


**Figure S4.** Full activation maps (top panel: axial slices from left to right; middle panel: coronal slices from anterior to posterior; bottom panel: sagittal slices from left to right) for the Active minus Passive contrast parametric modulation analysis (PMOD). Details for correction and display are the same as principal Figure 2C.

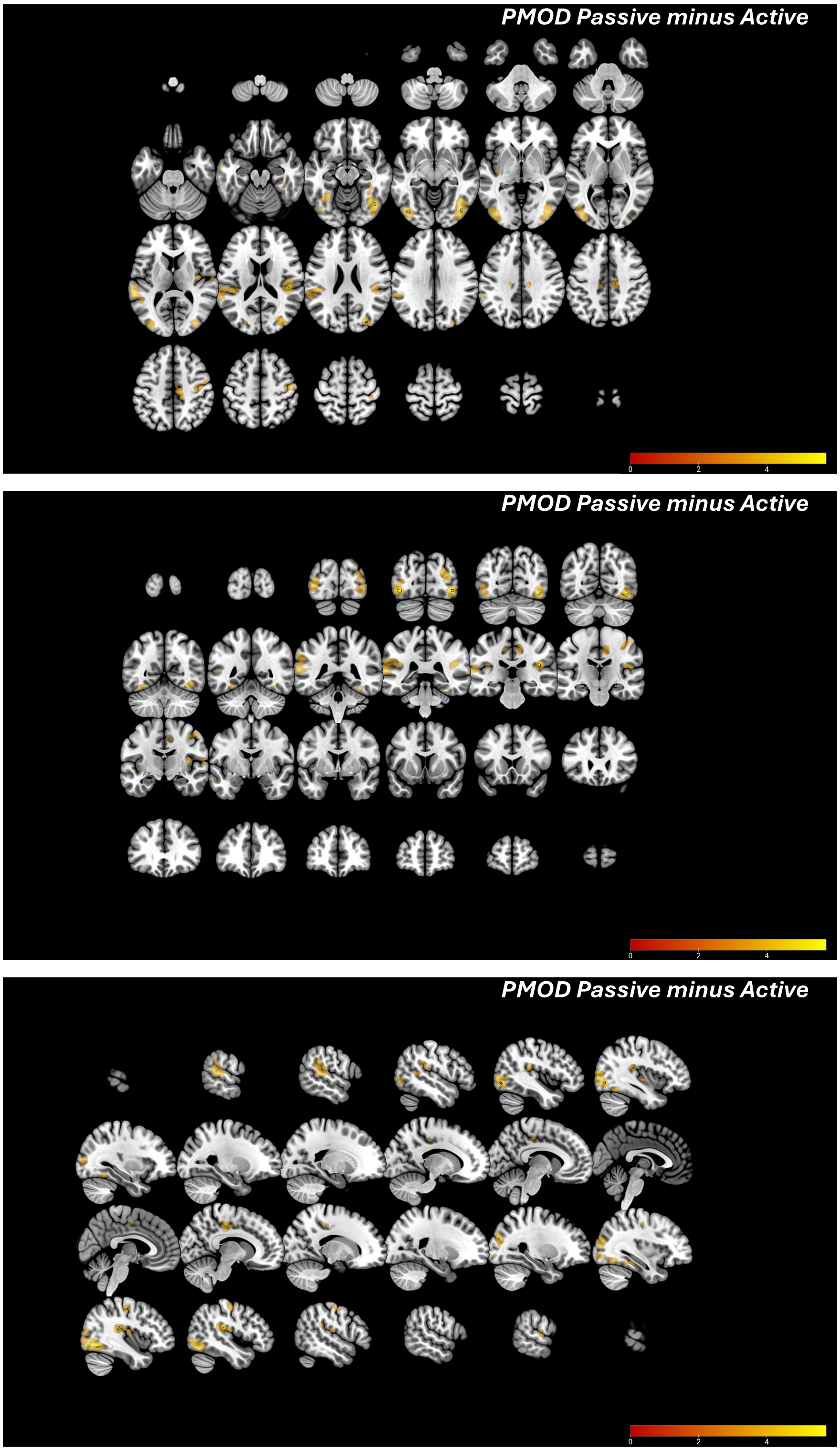


**Figure S5.** Full activation maps (top panel: axial slices from left to right; middle panel: coronal slices from anterior to posterior; bottom panel: sagittal slices from left to right) for the Passive minus Active contrast parametric modulation analysis (PMOD). Details for correction and display are the same as principal Figure 2C.

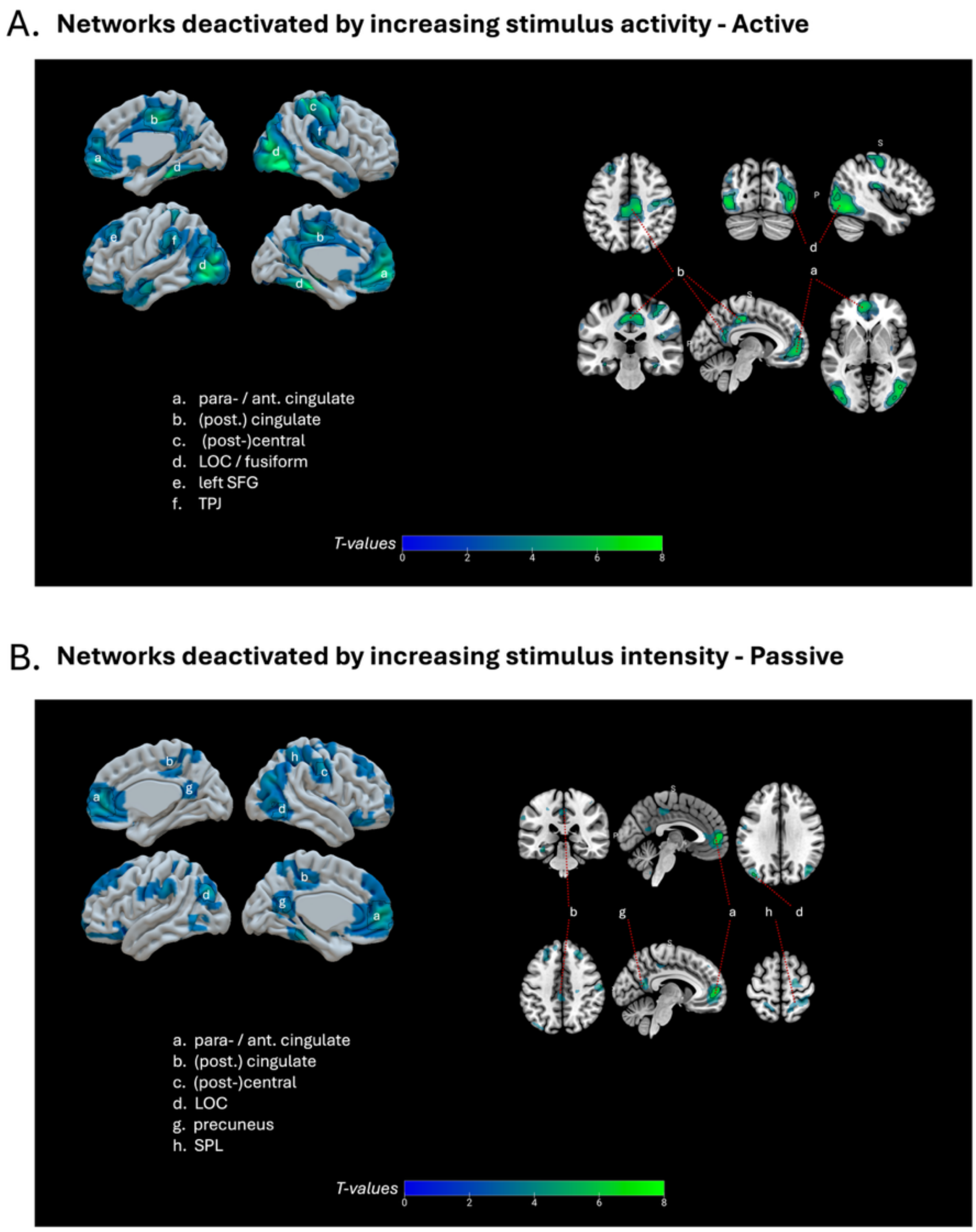


**Figure S6. Networks deactivated by increasing auditory stimulus intensity, with and without a task.**

A: Active condition deactivation parametric modulation t-values. B: Passive condition deactivation parametric modulation t-values. All slices are displayed in neurological convention. All slices are displayed in neurological convention. 3D surfaces and slices display corrected activations delimited by a black contour (FWEc, $p<0.05$), and uncorrected ones ($p<0.001$) in a less opaque shade. Ant. cingulate = anterior cingulate cortex; post. cingulate = posterior cingulate cortex; LOC = lateral occipital cortex; SFG = superior frontal gyrus; SPL = superior parietal lobule.

| MINUS ACTIVE | | | | | | | | | | |
|---|---|---|---|---|---|---|---|---|---|---|
| **Cluster label** | **Cluster size (voxels)** | **Side** | **Peak coordinates (mm)** | | | **Peak T-value** | **Z-score of peak T-value** | **Peak p-value (FWEc)** | **Cluster p-value (FWEc)** | **Network(s) of interest** |
| | | | **x** | **y** | **z** | | | | | |
| Lateral occipital cortex | 1287 | R | 44 | -62 | -10 | 9,77 | 7,22 | < 0.001 | < 0.001 | Visual |
| | | R | 36 | -82 | 13 | 9,05 | 6,88 | < 0.001 | | |
| | | R | 32 | -85 | 23 | 8,61 | 6,66 | < 0.001 | | |
| Lateral occipital cortex / temporal fusiform / parahippocampal | 758 | L | -48 | -75 | 0 | 8,85 | 6,78 | < 0.001 | < 0.001 | Visual |
| | | L | -31 | -32 | -14 | 8,41 | 6,56 | < 0.001 | | |
| | | L | -38 | -90 | 10 | 8,33 | 6,52 | < 0.001 | | |
| Posterior cingulate (superior part) | 398 | R | 12 | -25 | 48 | 8,74 | 6,73 | < 0.001 | < 0.001 | DMN |
| | | L | -6 | -25 | 46 | 8,27 | 6,49 | < 0.001 | | |
| | | R | 6 | -12 | 48 | 6,63 | 5,56 | < 0.001 | | |
| Para- / anterior cingulate | 395 | L | -11 | 52 | 0 | 8,1 | 6,4 | < 0.001 | < 0.001 | DMN |
| | | L | -8 | 58 | 20 | 5,94 | 5,12 | 0,01 | | |
| Post central gyrus / superior parietal lobule | 408 | R | 29 | -42 | 60 | 7,92 | 6,3 | < 0.001 | < 0.001 | Somato-motor |
| | | R | 42 | -18 | 53 | 7,89 | 6,29 | < 0.001 | | |
| | | R | 52 | -15 | 53 | 7,55 | 6,1 | < 0.001 | | |
| Parietal operculum / (temporo-parietal junction) | 47 | R | 44 | -22 | 20 | 7,16 | 5,88 | < 0.001 | < 0.001 | Salience / DMN |
| Post central gyrus | 22 | L | -24 | -40 | 58 | 6,42 | 5,43 | < 0.001 | < 0.001 | Somato-motor |
| Superior / middle frontal gyrus | 37 | L | -24 | 28 | 40 | 6,42 | 5,43 | < 0.001 | < 0.001 | DMN |
| Posterior cingulate / precuneus | 34 | L | -8 | -55 | 16 | 6,22 | 5,3 | < 0.001 | < 0.001 | DMN |
| | | L | -6 | -52 | 26 | 5,76 | 5 | 0,01 | | |
| Middle temporal gyrus | 3 | L | -61 | -5 | -20 | 6,08 | 5,21 | < 0.001 | 0,013 | DMN |
| Posterior insula | 13 | R | 36 | -12 | 6 | 6,05 | 5,19 | 0,01 | 0,001 | Somatosensory* |
| Frontal pole | 9 | L | -34 | 32 | -14 | 6 | 5,16 | 0,01 | 0,003 | Control / DMN |
| Pre central gyrus | 3 | R | 24 | -12 | 68 | 5,86 | 5,06 | 0,01 | 0,013 | Somato-motor |
| Supramarginal gyrus | 23 | L | -64 | -35 | 33 | 5,79 | 5,02 | 0,01 | < 0.001 | Dorsal attention / control |
| Para- / anterior cingulate | 6 | R | 6 | 52 | 13 | 5,76 | 5 | 0,01 | 0,006 | DMN |
| Lateral occipital cortex | 1 | L | -44 | -75 | 33 | 5,28 | 4,66 | 0,05 | 0,027 | Visual |
| **MINUS PASSIVE** | | | | | | | | | | |
| **Cluster label** | **Cluster size (voxels)** | **Side** | **Peak coordinates (mm)** | | | **Peak T-value** | **Z-score of peak T-value** | **Peak p-value (FWEc)** | **Cluster p-value (FWEc)** | **Network(s) of interest** |
| | | | **x** | **y** | **z** | | | | | |
| Para- / anterior cingulate | 195 | L | -6 | 55 | 6 | 6,61 | 5,55 | 0,001 | 0 | DMN |
| Lateral occipital cortex | 6 | R | 54 | -65 | -7 | 5,97 | 5,14 | 0,006 | 0,006 | Visual |
| Lateral occipital cortex | 13 | L | -41 | -80 | 30 | 5,76 | 5 | 0,012 | 0,001 | Visual |
| Para- / anterior cingulate | 10 | R | 12 | 48 | -4 | 5,64 | 4,91 | 0,017 | 0,003 | DMN |
| Temporal fusiform / parahippocampal | 4 | L | -31 | -32 | -14 | 5,55 | 4,86 | 0,022 | 0,01 | Visual |
| Precuneus | 5 | L | -11 | -58 | 18 | 5,46 | 4,79 | 0,029 | 0,008 | DMN |
| Post central gyrus | 1 | R | 62 | -15 | 43 | 5,3 | 4,68 | 0,046 | 0,027 | Somato-motor |

**Table S1. Parametric modulation fMRI results: negative contrasts.** The clusters and sub-clusters peaks listed here result from whole-brain parametric modulation analyses and survived cluster-level FWE correction ($p < 0.05$). X, y and z coordinates refer to the Montreal Neurologic Institute (MNI) space. The last column indicates network(s) the clusters belong to, according to Yeo et al. 2011 (Yeo et al., 2011) unless specified otherwise (* Garcia-Larrea (Garcia-Larrea, 2012); NA = Not applicable). Uncorrected p-values are not displayed here because they are all < 0.001 since it is only corrected results.

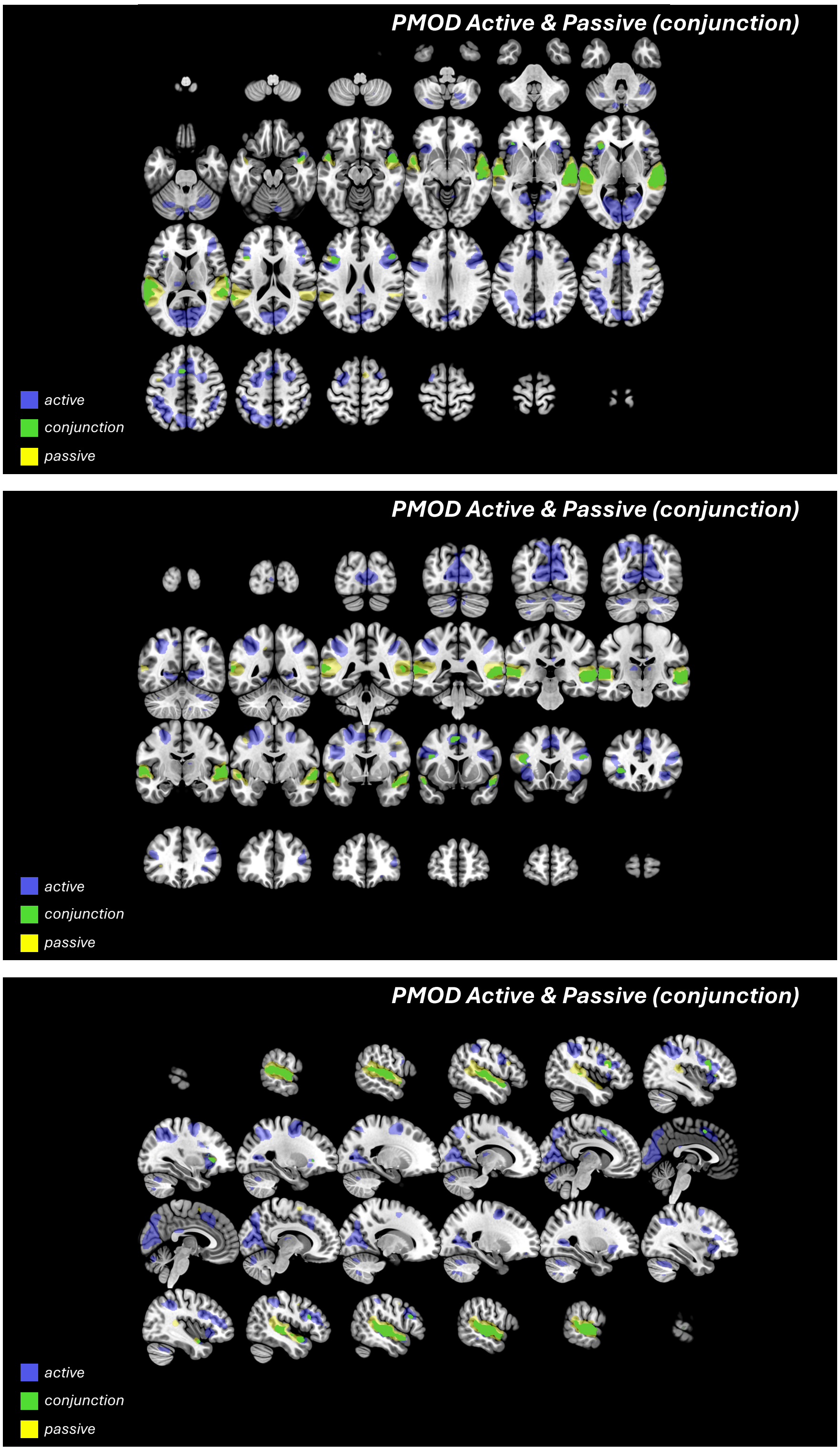


**Figure S7.** Full activation maps (top panel: axial slices from left to right; middle panel: coronal slices from anterior to posterior; bottom panel: sagittal slices from left to right) for the Active / Passive conjunction contrast parametric modulation analysis (PMOD). Details for correction and display are the same as principal Figure 3A.

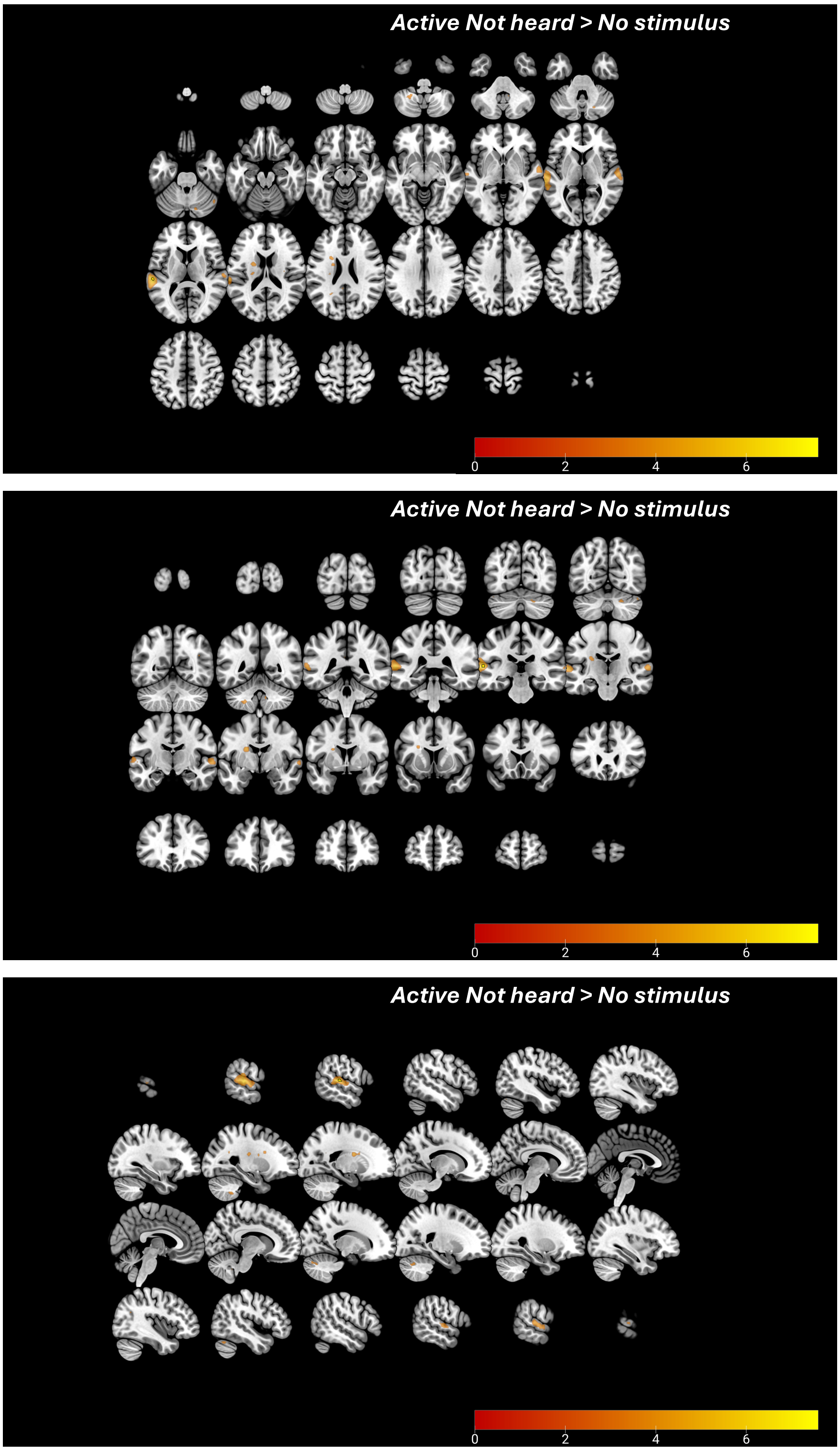


**Figure S8.** Full activation maps (top panel: axial slices from left to right; middle panel: coronal slices from anterior to posterior; bottom panel: sagittal slices from left to right) for the Active Not heard minus No stimulus analysis. Details for correction and display are the same as principal Figure 4A.

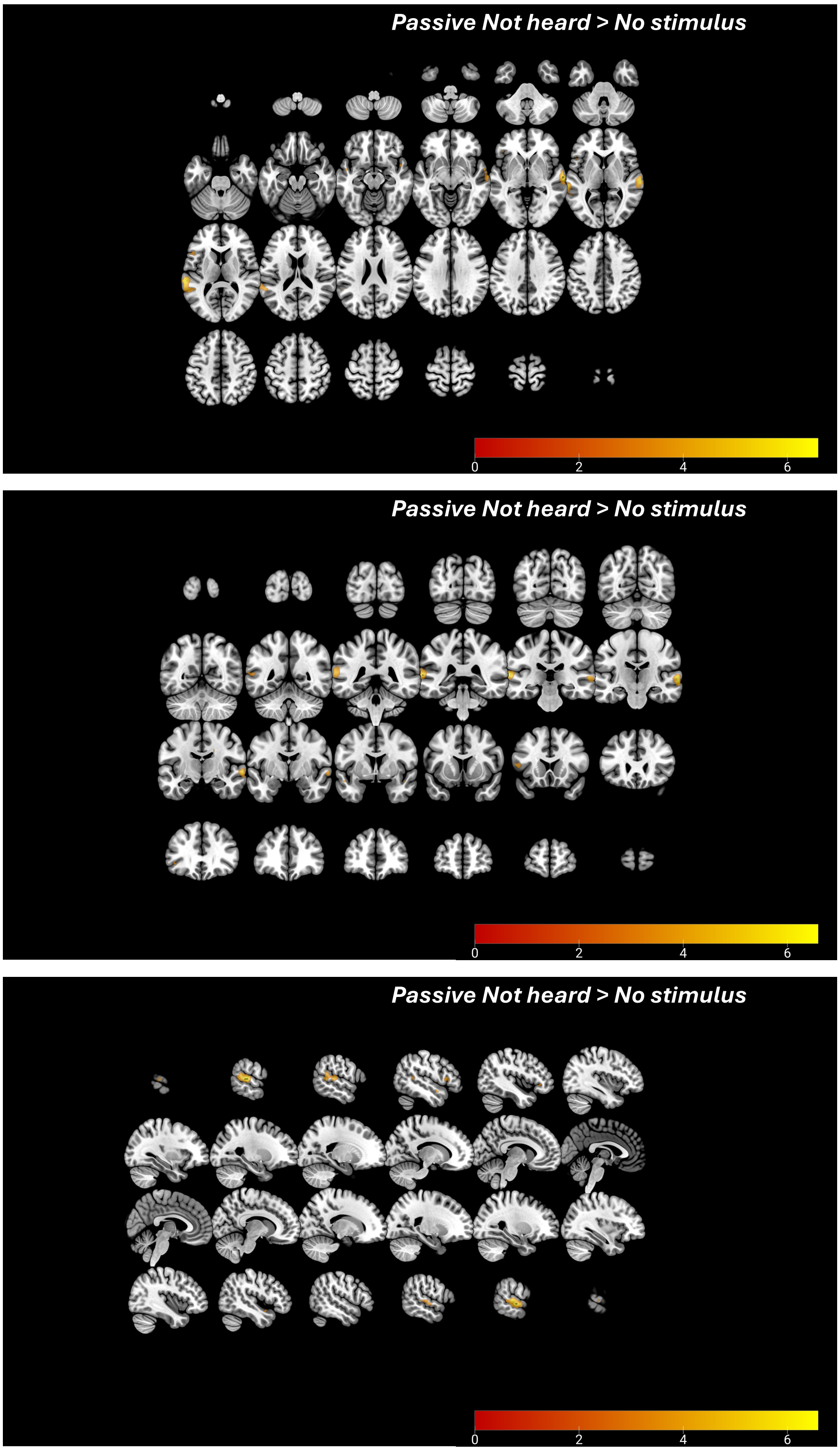


**Figure S9.** Full activation maps (top panel: axial slices from left to right; middle panel: coronal slices from anterior to posterior; bottom panel: sagittal slices from left to right) for the Passive Not heard (answer "Other") minus No stimulus analysis. Details for correction and display are the same as principal Figure 4A.

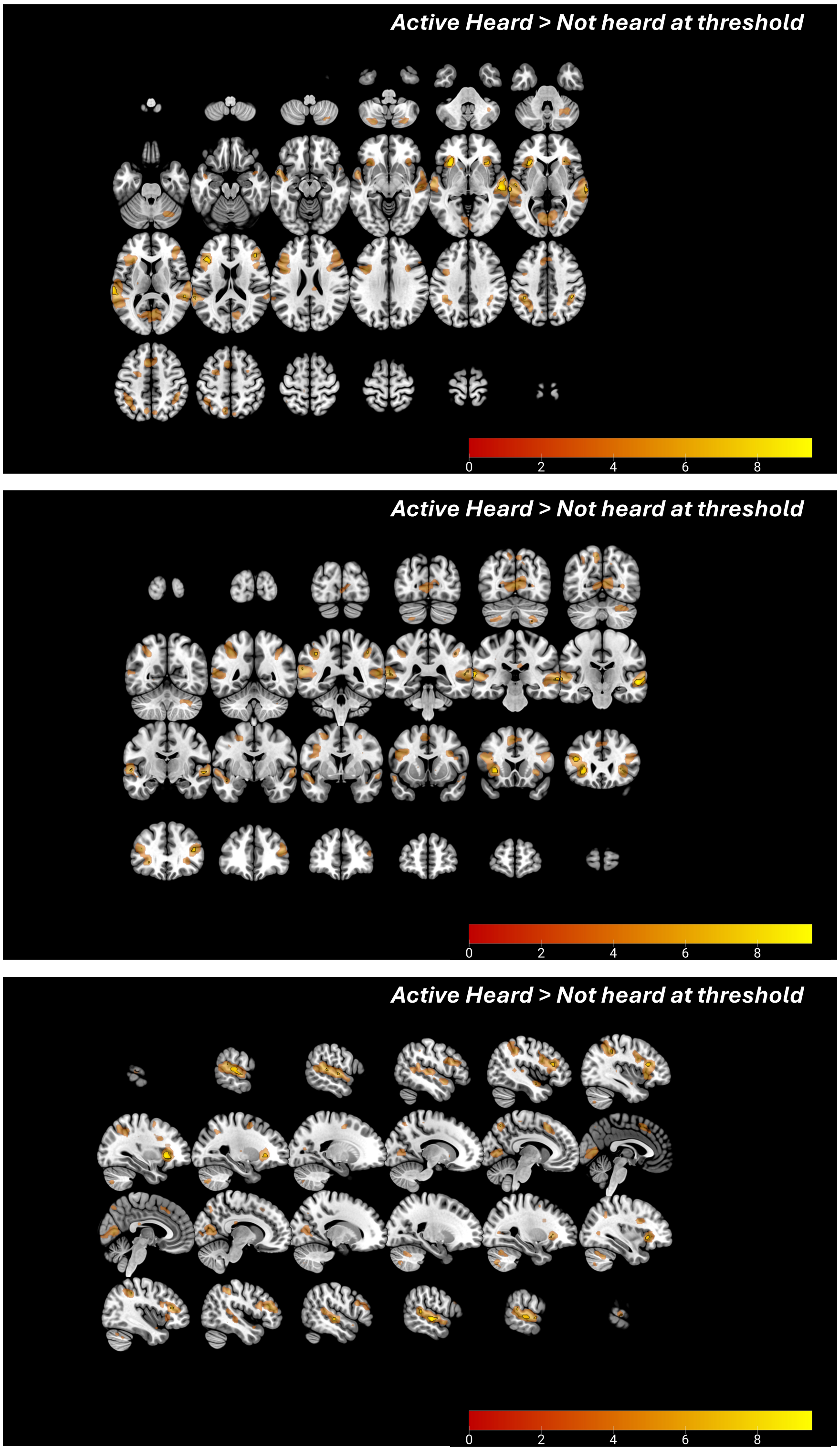


**Figure S10.** Full activation maps (top panel: axial slices from left to right; middle panel: coronal slices from anterior to posterior; bottom panel: sagittal slices from left to right) for the Active Heard minus Not heard at threshold analysis. Details for correction and display are the same as principal Figure 4B.

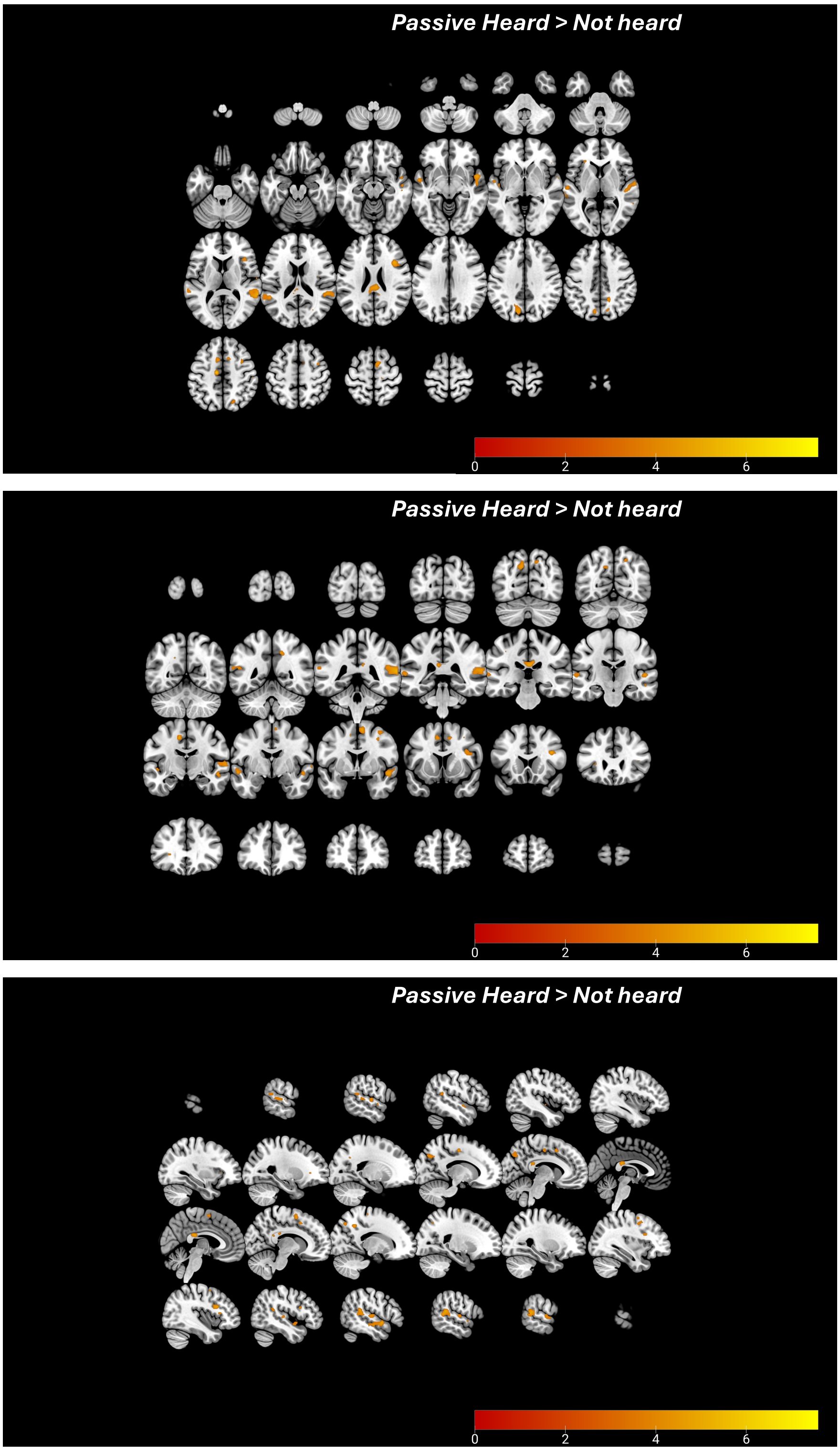


**Figure S11.** Full activation maps (top panel: axial slices from left to right; middle panel: coronal slices from anterior to posterior; bottom panel: sagittal slices from left to right) for the Passive Heard minus Not heard (answer "Sound" minus "Other") analysis. Details for correction and display are the same as principal Figure 4C.

A. **Composite model for Passive Report: The sound > Other**

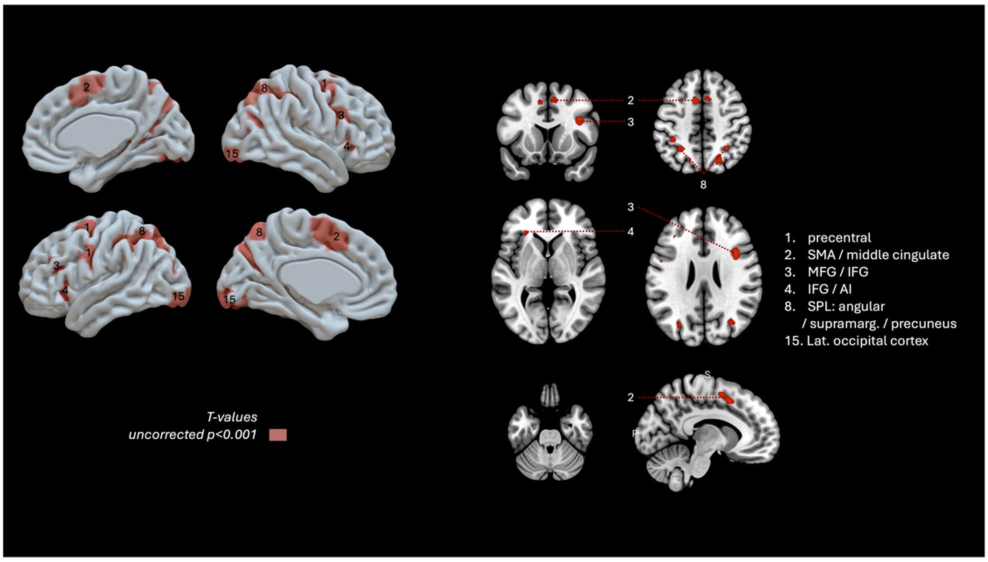


B. **Composite model for Passive Report: stimulus intensity parametric modulation**

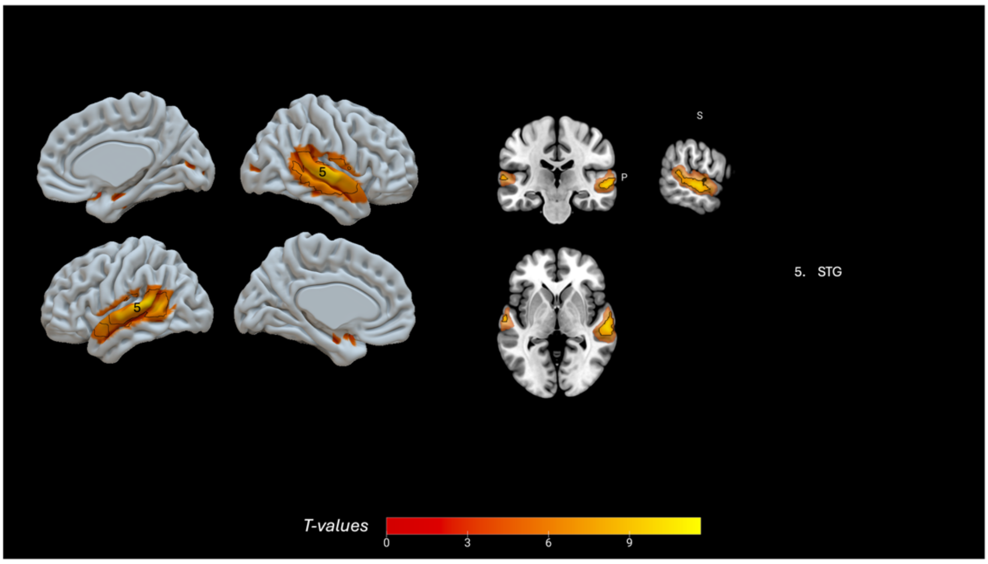


**Figure S12. Composite model for Passive conscious perception.** A: "The sound" minus "Other" contrast t-values. B: Stimulus intensity parametric modulation t-values. All slices are displayed in neurological convention and show corrected activations (if existent) delimited by a black contour (FWEc, p<0.05), and uncorrected ones (p<0.001) in a less opaque shade. 3D surfaces display corrected (FWEc, p<0.05) and uncorrected (p<0.001) binarized t-maps. SMA = supplementary motor area; MFG = middle frontal gyrus; IFG = inferior frontal gyrus; AI = anterior insula; SPL = superior parietal lobule; IPS = intraparietal sulcus; STG = superior temporal gyrus.

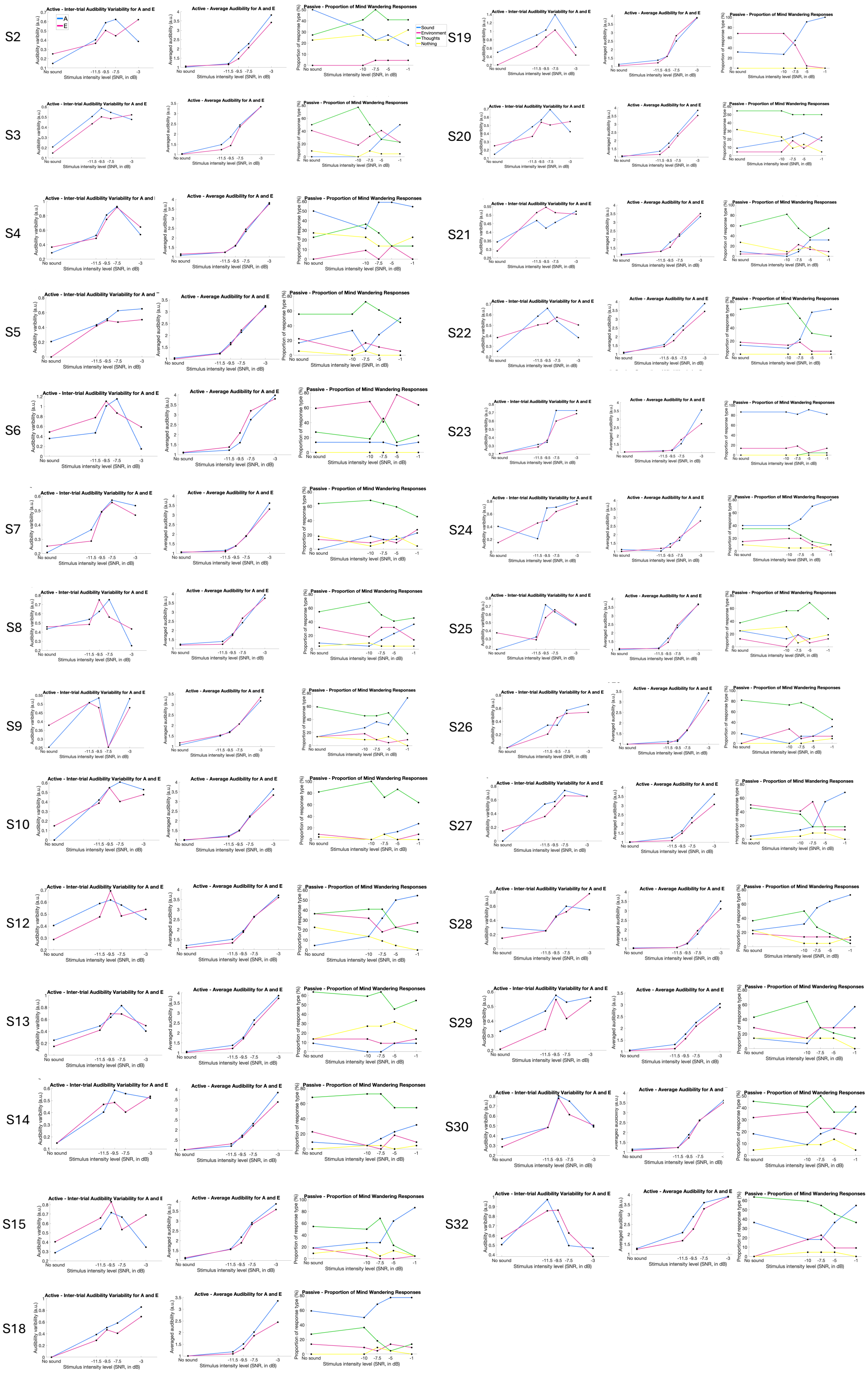


**Figure S13. Individual behavioural results.** Each row corresponds to one subject included in the main analysis; from left to right: inter-trial variability as a function of stimulus intensity (active condition), audibility rating as a function of stimulus intensity (active condition) and mind-wandering responses as a function of stimulus intensity (passive condition).

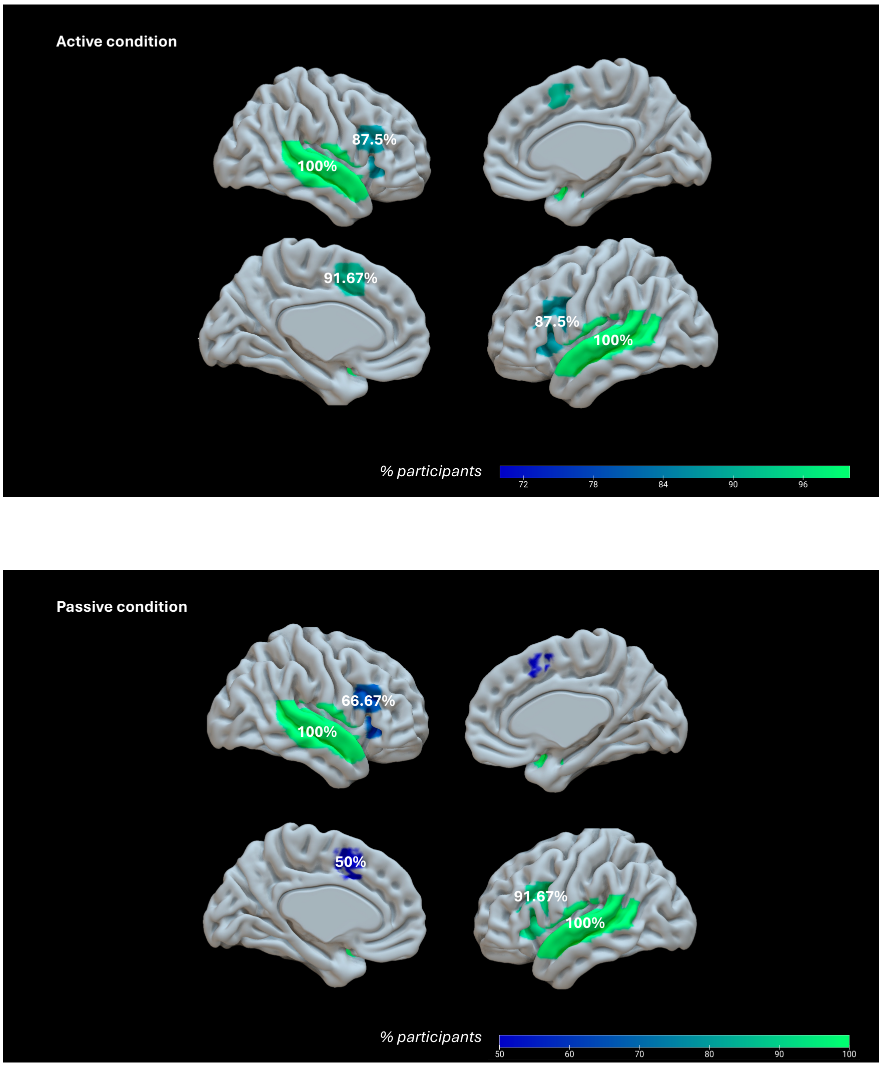


**Figure S14. Proportion of participants' RFE-selected beyond primary auditory ROI voxels in the conjunction ROIs.** Each panel represents the percentage 3D map of participants that have at least one voxel included in the conjunction ROIs (superior temporal gyrus, anterior cingulate cortex, PFC / anterior insula) after applying RFE (recursive feature elimination) to define extra-primary auditory individual clusters from which single trial activity is extracted, for each condition (top panel: Active; bottom panel: Passive). Of note, Heschl's gyrus were not included but for reasons of representations, we displayed the whole STG regions.

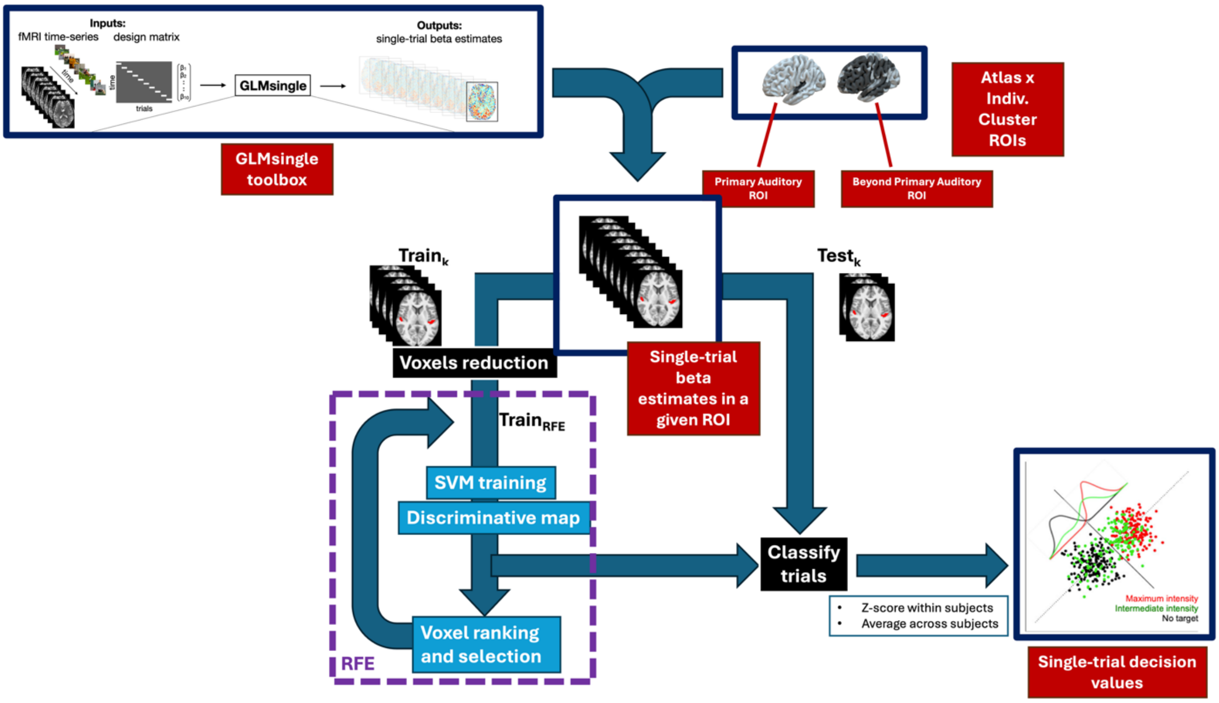


**Figure S15. Schematic representation of the pipeline used for multivariate single-trial extraction.** As a first step, we designed a single-trial GLM to the fMRI data and applied GLMSingle functions (GLMdenoise, HRF selection, fractional ridge regression) to improve the signal to noise ratio of the extracted single-trial Beta estimates across the whole brain. Next, we extracted those single-trial Beta estimates within two ROIs. Those ROIs were previously defined by overlapping each subject's indivual univariate cluster of activation to the stimulus intensity (parametric modulation analysis) with atlas-defined bilateral Heschl's gyri to create (1) primary auditory ROIs (overlapping Heschl's gyri) and (2) beyond primary auditory ROIs (the remaining voxels of the cluster after extracting Heschl's gyri). We then fed a linear decoder (SVM, support vector machine) with each of these individual ROI-extracted single-trial Beta estimates clusters and trained it to decode maximum intensity level stimulus versus no stimulus; at each cross-validation fold, it therefore defined a decision hyperplane to distinguish between these two categories, and we then tested it on the remaining trials, this time including all stimulus intensities, in order to extract one decision value (ie, distance from the decision hyperplane) for each single trial's cluster of betas. Additionally, we performed RFE (recursive feature elimination) nested into the decoder's training in order to previously select each participant's and each ROI's most informative subset of voxels. As an output we obtained one single-trial "decision value" or neural value for each trial, each ROI (primary auditory and beyond primary auditory), each subject and each condition (Active / Passive), which we z-scored. These values were then averaged across participants to obtain the group-level single-trial dynamics for each ROI and condition.

Adapted from Prince et al. eLife 2022, De Martino et al. NeuroImage 2008 and Sergent et al. Nature Communications 2021.

| Participant number | Spearman Rho | P-value |
|---|---|---|
| ***2*** | ***-0,9*** | ***0,083*** |
| 3 | 0,975 | 0,033 |
| 4 | 0,564 | 0,4 |
| 5 | 0,5 | 0,45 |
| ***6*** | ***-0,354*** | ***0,8*** |
| 7 | 0,667 | 0,267 |
| 8 | 0,9 | 0,083 |
| 9 | 0,9 | 0,083 |
| 10 | 0,9 | 0,083 |
| 12 | 1 | 0,017 |
| 13 | 0,289 | 0,8 |
| 14 | 0,9 | 0,083 |
| 15 | 0,975 | 0,033 |
| 18 | 0,872 | 0,1 |
| 19 | 0,9 | 0,083 |
| 20 | 0,564 | 0,367 |
| 21 | 0,791 | 0,133 |
| 22 | 0,9 | 0,083 |
| ***23*** | ***-0,264*** | ***0,667*** |
| 24 | 0,975 | 0,033 |
| ***25*** | ***-0,316*** | ***0,6*** |
| 26 | 0,359 | 0,633 |
| 27 | 1 | 0,017 |
| 28 | 1 | 0,017 |
| 29 | 0,872 | 0,067 |
| 30 | 0,821 | 0,133 |
| 32 | 0,527 | 0,467 |

**Figure S16. Correlation coefficient between each participant's proportion of mind-wandering answer "The sound" and the stimulus intensity level in the Passive condition.** In red, participants with a negative correlation, whose mind-wandering probes were therefore not informative regarding their perception, were removed from the prediction performance evaluation.
